# From peptides to proteins: coiled-coil tetramers to single-chain 4-helix bundles

**DOI:** 10.1101/2022.08.04.502660

**Authors:** Elise A. Naudin, Katherine I. Albanese, Abigail J. Smith, Bram Mylemans, Emily G. Baker, Orion D. Weiner, David M. Andrews, Natalie Tigue, Nigel J. Savery, Derek N. Woolfson

## Abstract

The design of completely synthetic proteins from first principles—*de novo* protein design—is challenging. This is because, despite recent advances in computational protein-structure prediction and design, we do not understand fully the sequence-to-structure relationships for protein folding, assembly, and stabilization. Antiparallel 4-helix bundles are amongst the most studied scaffolds for *de novo* protein design. We set out to re-examine this target, and to determine clear sequence-to-structure relationships, or *design rules*, for the structure. Our aim was to determine a common and robust sequence background for designing multiple *de novo* 4-helix bundles, which, in turn, could be used in chemical and synthetic biology to direct protein-protein interactions and as scaffolds for functional protein design. Our approach starts by analyzing known antiparallel 4-helix coiled-coil structures to deduce design rules. In terms of the heptad repeat, ***abcdefg***—i.e., the sequence signature of many helical bundles—the key features that we identify are: ***a*** = Leu, ***d*** = Ile, ***e*** = Ala, ***g*** = Gln, and the use of complementary charged residues at ***b*** and ***c***. Next, we implement these rules in the rational design of synthetic peptides to form antiparallel homo- and heterotetramers. Finally, we use the sequence of the homotetramer to derive a single-chain 4-helix-bundle protein for recombinant production in *E. coli*. All of the assembled designs are confirmed in aqueous solution using biophysical methods, and ultimately by determining high-resolution X-ray crystal structures. Our route from peptides to proteins provides an understanding of the role of each residue in each design.

## INTRODUCTION

*De novo* protein design is advancing rapidly,^1, 2^ indeed our ability to design proteins from scratch is said to have come of age.^3^ Protein-design processes have evolved over the past four decades from minimal design that uses straightforward chemical principles, through rational design that incorporates sequence-to-structure relationships learnt from natural proteins, to computational design that builds proteins from fragments or parametric templates and scores them using statistical and physical forcefields.^4^ Today, the field has also progressed to include state-of-the-art computational methods such as Artificial Intelligence and Machine Learning that allow protein designers to “hallucinate” proteins in the computer ahead of making and characterizing in the laboratory.^5, 6^ Of course, a possible downside of these new innovations is that we no longer understand what we are—that is, *what the computer is—* designing, and one of the original motivations for the field is potentially being lost; namely, the notion of testing our understanding of the chemical and physical principles of protein folding, assembly, and stability.^4^ That aside, the field has matured from being structure-centric to one committed to developing synthetic proteins with useful functions that mimic or augment natural protein functions both *in vitro* and in biological contexts.^6–11^ Whilst many challenges remain,^2, 4^ these are truly exciting and promising times for *de novo* protein design.

Over the past 4 decades, 4-helix bundles (4HBs) have been one of the go-to targets for *de novo* peptide and protein design.^1, 12–14^ Historically, 4HB design began with minimal approaches employing patterns of hydrophobic (leucine) and polar (glutamate and lysine) residues to design single short amphipathic a helices that self-associate due to the hydrophobic effect, or to program libraries of single-chain 4-helix proteins that fold through hydrophobic collapse.^15–18^ Again, these design approaches have evolved by incorporating biological information and rational design, which led more readily to high-resolution X-ray crystal structures.^4, 19–22^ Most recently, interest has shifted to using computational methods involving backbone and fold parametrization, optimization of core packing, and specific interaction networks between core residues.^23–28^ Furthermore, 4HBs present a variety of assembly modes for protein designers to target, including: single peptides that associate to tetramers, helix-loop-helix constructs that can dimerize, and self-contained single-chain proteins.^17, 29–31^ However, they also present pitfalls—or alternate states—that designers must learn to navigate away from using negative design principles.^22, 32, 33^ For instance, for the tetramers, adjacent helices can have all-parallel, antiparallel, or mixed arrangements; and for helix-loop-helix and single-chain systems various topologies are possible.^31, 34, 35^ These different architectures, the relatively large hydrophobic cores, and the apparent robustness to modification have been exploited to functionalize 4HBs and introduce small-molecule binding,^36^ catalysis,^37–39^ allostery,^40^ and the control of protein-protein interaction including regulation of gene expression.^41, 42^

One specific type of 4HB form α-helical coiled coils (CCs). In CCs, tight and regular packing between side chains of neighboring helices—known as knobs-into-holes (KIH) packing—specifies the structure, including defining oligomer state, partner preferences, and helix orientation. This has proved extremely powerful in rational and computational design of CCs.^43–45^ In more detail, CCs are supercoiled assemblies of amphipathic a helices. Generally, CC assembly is programmed by sequence repeats of hydrophobic (***h***) and polar (***p***) residues, ***hpphppp***, often called heptads and denoted ***abcdefg*** (Figure 1A).^44, 46^ Many sequence-to-structure relationships, especially at the hydrophobic ***a/d*** interface, have been learnt from analysis of natural structures and empirical studies.^19, 47^ In turn, these have been used to deliver a wide range of structured and increasingly functional CC designs.^4, 43, 44^ For example, our own basis set of *de novo* CCs currently comprises parallel assemblies from dimer to nonamer,^20, 48, 49^ and these are being used increasing by us and others in various applications.^41, 50–55^ That all said, designing antiparallel CC assemblies from first principles has been more challenging.^33, 56–58^ Moreover, subtle changes in primary sequence or even experimental conditions can induce switches from energetically close parallel assemblies to antiparallel conformations.^33, 59–61^ For example, recently, we reported the rational redesign of an antiparallel CC tetramer, apCC-Tet, following the serendipitous discovery of up-down-up-down tetramers adopted by a point mutation of our original parallel hexamer, CC-Hex.^33^

**Figure 1.**
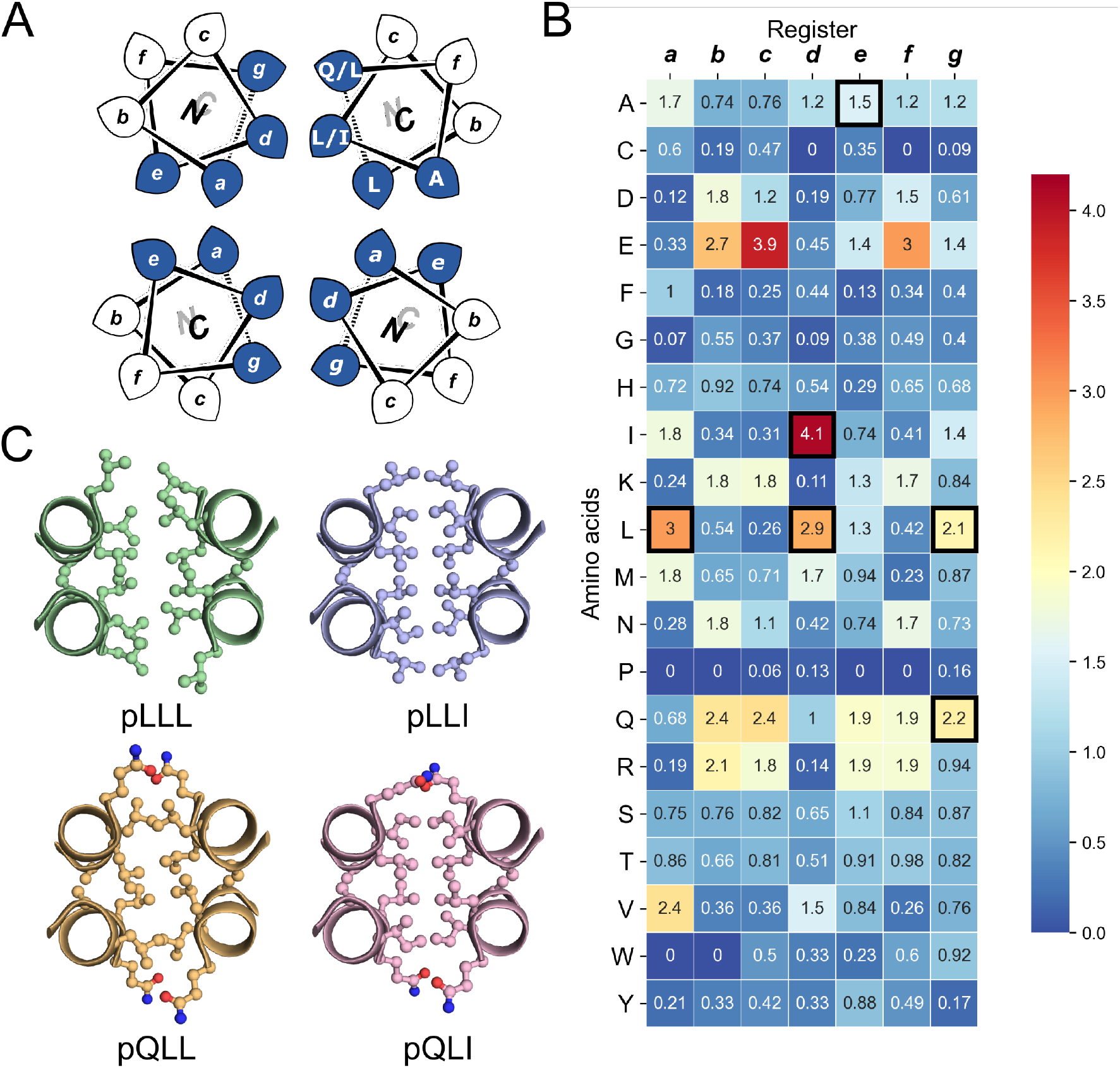
The rational design of new sequences to form antiparallel CC tetramers. (A) Helical-wheel representation of an antiparallel four-helix CC. Sequences have heptad repeats, ***abcdefg***. The interfacial positions, ***a***, ***d***, ***e***, and ***g***, where our designs focused are highlighted in blue. Selected residues in our designed sequences are shown in the top right helix. The *N-to-C*-terminal directions of the helices are indicated with the ‘N’ or ‘C’ with the darker font indicating that that end is closer to the viewer. (B) Propensity table of residues for each amino acid at each position of the heptad repeat for antiparallel 4-helix CCs found in CC+.^65^ Raw counts (Table S1) were normalized using the amino-acid frequencies in SWISS-PROT to give the propensity scale shown as a heat map (high, red; low, blue). A propensity of 0 indicates that no examples of that amino acid were found at that position in the database. Residues identified for the design of the new antiparallel tetramer sequences are highlighted with dark square boxes. (C) Heptad-repeat slices through the AlphaFold2-multimer^67–69^ models for each designed sequence: pLLL (top left), pLLI (top right), pQLL (bottom left), and pQLI (bottom (right). Images for panel C were generated in PyMOL (pymol.org).

Establishing clear principles for *de novo* design, such as sequence-to-relationships for a given target, would help navigate the complex energy landscape of helical assemblies. Moreover, it would deliver *design rules* to direct the assembly of different helical states to improve on and expand toolkits such as our own CC basis set and similar sets from others.^62– 64^ In turn, these would be provide platforms for redesign and applications where the impact of modifications required for functionalization could be anticipated.

Here, we elaborate a set of sequence-to-structure relationships for the design of CC-based antiparallel 4HBs. By inspecting the structural database of CCs (CC+),^65^ we deduce clear design rules for this target. In turn, these are used to deliver three *de novo* structures: an antiparallel homotetramer, apCC-Tet*, a heterotetramer, apCC-Tet*-A2B2, and a singlechain 4HB, sc-apCC-4. All three designs are characterized fully in solution, and to high-resolution by determining X-ray crystal structures. The designs are hyperstable with respect to thermal and chemical denaturation, and they fold, assemble, and function in *E. coli*. These properties make them ideal scaffolds for functionalization and for future *in vitro* and subcellular applications.

## RESULTS AND DISCUSSION

### Rational designs based on analysis of known coiled-coil structures

To garner sequence-to-structure relationships for designing of our target antiparallel CC tetramers and bundles, we analyzed relevant structures in the CC+ database.^65^ We selected sequences for all antiparallel, four-helix, homo- and heteromeric CC assemblies with ≤50% sequence redundancy. The sequences were used to compile an amino-acid profile for the heptad repeats, ***abcdefg***, of these structures (Figure 1A); ^44, 46^ the raw counts are available in Table S1. The profile was normalized using expected amino-acid frequencies from SWISS-PROT to give propensities for each residue at each position of the ***a – g*** repeat (Figure 1B). Next, we used these propensities to deduce new *de novo* repeat sequences. We focused on the interfacial positions, ***g***, ***a***, ***d*** and ***e***, as these contribute most to CC folding, stability, and oligomer state specification (Figure 1A).^44^ For the ***a*** site, we selected leucine (Leu, L) as this was overwhelmingly preferred in the profile with a propensity of 3; *i.e*., it occurred three times more frequently than expected by chance at this site (Figure 1B). We did not consider the next most prevalent residues at ***a***, the β-branched isoleucine (Ile, I) and valine (Val, V), as these are known to favor both parallel and antiparallel dimers when placed at this site.^19^ Two residues, Ile and Leu, had high propensities for ***d***-occurring at ≈4x and ≈3x the expected frequency, respectively, so we considered both in our initial designs. At ***e***, no residues appeared above twice the expected frequency. Therefore, we opted for Ala at this site, as it occurred frequently in the dataset (Table S1) and is known to promote antiparallel tetramers, *via* Alacoil formation.^33, 66^ Finally, Leu and glutamine (Gln, Q) had propensity values exceeding 2 for the ***g*** position. Therefore, both residues were investigated at ***g*** in our initial designs.

Consequently, our analysis led to four distinct sequence combinations with the potential to form antiparallel tetramers: namely, for ***g*** ➔***f*** repeats, *L/Q-L-**b**-**c**-I/L-A-**f.*** Following previous CC designs,^20, 33, 48, 49^ we concatenated 4 copies of each repeat into each of 4 designed homomeric peptide sequences. We used the unspecified ***b*** and ***c*** sites to direct antiparallel assemblies further, specifically in homomers. Our rationale was to create a ‘bar-magnet’ charge pattern in the sequences by placing negatively charged glutamic acid (Glu, E) at the ***b*** and ***c*** sites of the first two heptad repeats, and positively charged lysine (Lys, K) at these sites in the two *C*-terminal repeats.^33^ The sequences were completed with the remaining 4 ***f*** sites filled with Gln, Lys, tryptophan (Trp, W), and Gln, respectively. The final sequences were capped with glycine (Gly, G) at both ends and *N-*terminally acetylated and *C-*terminally amidated (Table 1). Initially, we named the sequences after the residues at the ***g***, ***a*** and ***d*** sites, *i.e*., pLLL, pLLI, pQLL, and pQLI.

Ahead of experiments, we modelled the four new sequences using the AlphaFold2-multimer predictor (Figure 1C and Figures S1-S4).^67–69^ Encouragingly, the AlphaFold2 predictions for both Q@***g*** sequences, pQLL and pQLI, gave antiparallel tetramers as designed and with high confidence (Figure 1C) even when an oligomeric state larger than 4 was provided as a target to AlphaFold2 (Figures S3-S4). By contrast, although the L@***g*** sequences, pLLL and pLLI, could be predicted to form antiparallel 4HBs by AlphaFold2 (Figure 1C), this was not consistently observed when higher chain numbers were used; in these cases, higher-order α-helical assemblies were predicted (Figure S1 and S2).

**Table 1.** Designed sequences and summary of biophysical data for the principal analogues.

| Peptide name | Sequence and register<br>*gabcdef gabcdef gabcdef gabcdef loop* | CD Helix (%)<sup>b</sup> | SV (mass/monomer mass)<sup>c</sup> | XRD oligomeric state<sup>d</sup> |
| --- | --- | --- | --- | --- |
| pLLL | Ac-G LLEELAQ LLEELAK LLKKLAW LLKKLAQ G-NH<sub>2</sub> | 75 | 6.0 | - |
| pLLI | Ac-G LLEEIAQ LLEEIAK LLKKIAW LLKKIAQ G-NH<sub>2</sub> | 82 | 6.3 | - |
| pQLL | Ac-G QLEELAQ QLEELAK QLKKLAW QLKKLAQ G-NH<sub>2</sub> | 87 | 4.3 | - |
| pQLL<sup>3</sup> | Ac-G QLEELAK QLQQLAW QLKKLAQ G-NH<sub>2</sub> | 77 | 4.0 | - |
| pQLI<br>apCC-Tet* | Ac-G QLEEIAQ QLEEIAK QLKKIAW QLKKIAQ G-NH<sub>2</sub> | 94 | 4.5 | 4<br>(8a3g,<br>0.96 Å) |
| pQLI<sup>3</sup><br>apCC-Tet*<sup>3</sup> | Ac-G QLEEIAK QLQQIAW QLKKIAQ G-NH<sub>2</sub> | 89 | 4.3 | 4<br>(8a3i,<br>1.42 Å) |
| pQLI<sup>3</sup>-A<sub>2</sub>B<sub>2</sub><br>apCC-Tet*<sup>3</sup>-<br>A<sub>2</sub>B<sub>2</sub> | Ac-G QLEEIAK QLEEIAW QLEEIAQ G-NH<sub>2</sub><br>Ac-G QLKKIAK QLKKIAY QLKKIAQ G-NH<sub>2</sub> | 81 | 4.3 | 4<br>(8a3j,<br>2.1 Å) |
| sc-QLI-4<br>sc-apCC-4 | QLEEIAQ QLEEIAK QLKKIAW QLKKIAQ <u>GEPSAQG</u><br>QLEEIAQ QLEEIAK QLKKIAW QLKKIAQ <u>GPDSV</u><br>QLEEIAQ QLEEIAK QLKKIAW QLKKIAQ <u>GGTSGG</u><br>QLEEIAQ QLEEIAK QLKKIAW QLKKIAQ | 71 | 0.9 | 1<br>(8a3k,<br>2.0 Å) |

<sup>a</sup>Complete biophysical analysis can be found in the Supporting Information. <sup>b</sup>Fraction helix determined using the MRE<sub>222</sub> from circular dichroism spectroscopy. Conditions: 50 µM peptide concentration, phosphate buffered saline (PBS, 8.2 mM sodium phosphate dibasic, 1.8 mM potassium phosphate monobasic, 137 mM sodium chloride, 2.4 mM potassium chloride) solution, pH 7.4; or 25 µM single-chain protein in sodium phosphate buffer (50 mM sodium phosphate, 150 mM NaCl), pH 7.4. <sup>c</sup>Oligomeric states from sedimentation-velocity measurements in analytical ultracentrifugation. Conditions: 150 µM peptide concentration, PBS solution, pH 7.4; or 25 µM single-chain protein in 50 mM sodium phosphate, 150 mM NaCl, pH 7.4. <sup>d</sup>Oligomeric state from X-ray crystallography. PDB IDs and diffraction resolutions are specified in brackets.

### Experimental characterization of a robust antiparallel coiled-coil tetramer, apCC-Tet*

The four sequences pLLL, pLLI, pQLL and pQLI (Table 1) were synthesized by solidphase peptide synthesis (SPPS) and confirmed by MALDI-TOF mass spectrometry (Figures S5-S8). First, circular dichroism (CD) spectroscopy was used to assess the secondary structure and stability of the designs in aqueous buffer near neutral pH. All four peptides were highly a helical with characteristic minima at 208 and 222 nm (Figure 2A and Figures S11-S12). Furthermore, the structures resisted thermal denaturation up to 95 °C (Figures 2B and Figures S11-S12). Next, sedimentation-velocity (SV) experiments using analytical ultracentrifugation (AUC) revealed monodisperse oligomers in all four cases (Figures 2C and Figures S13-S16). Interestingly, and consistent with the AlphaFold2 modelling, both L@***g*** peptides, pLLL and pLLI, formed hexamers in solution (Figure S13-S14). This was despite the amino-acid profiles, and the precedence of L@***g*** in other 4HB designs, notably from computational design.^27, 70^ With hindsight, the hexamers that we observed might have been anticipated, as other *de novo* peptides with predominantly hydrophobic residues at ***g, a, d*** and ***e*** are Type-II CC sequences, which can form oligomers of >4, including a-helical barrels.^44, 48, 49^ Therefore, these two peptides, pLLL and pLLI, were not investigated further in this study, which aimed to deliver antiparallel 4HBs. By contrast, and consistent with the design target, both Q@***g*** peptides, pQLL and pQLI, returned tetrameric molecular weights in AUC by both sedimentation equilibrium (SE) and SV experiments (Figures S15-S17). Both of these designs were taken forward.

**Figure 2.**
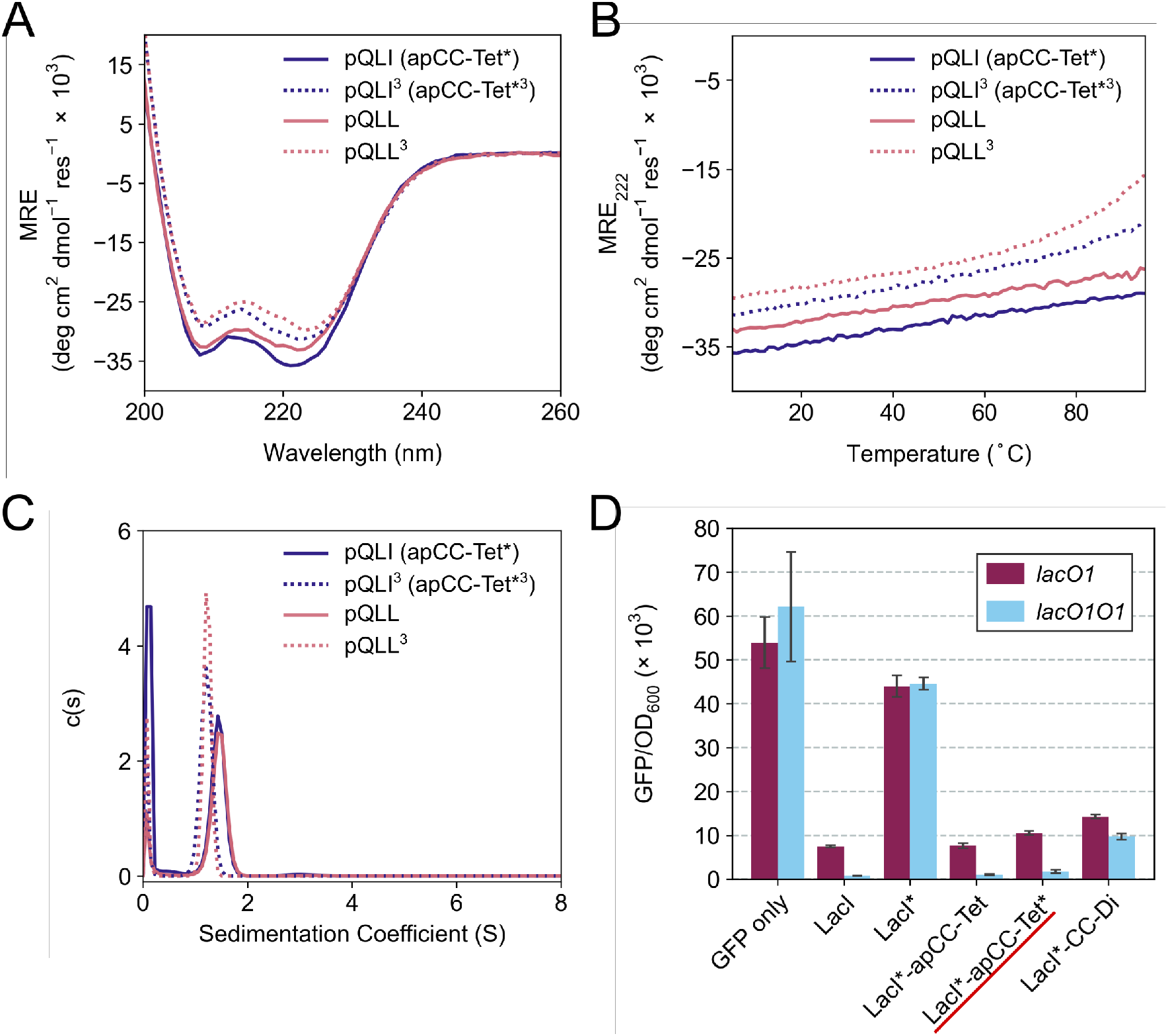
Biophysical and in-cell characterization of the homotetrameric peptides pQLI (apCC-Tet*) and pQLL. (A) CD spectra at 5 °C and (B) thermal responses of the CD signals at 222 nm (ramping up from 5 to 95 °C) of pQLI (apCC-Tet*), pQLI^3^ (apCC-Tet*^3^), pQLL, and pQLL^3^. Conditions: 50 μM peptide, PBS, pH 7.4. (C) Sedimentation-velocity data from AUC of pQLI, pQLI^3^, pQLL, and pQLL^3^. Fits returned masses of 4.5, 4.3, 4.3 and 4.0 × monomer mass, respectively. Conditions: 150 μM peptide, PBS, pH 7.4. (D) Transcription repression assay in *E. coli*. CC peptides were fused to LacI*, a destabilized variant of the Lac repressor. The reporter gene, GFP, was expressed from the *lacUV5* promoter with or without an additional *lac O1* operator placed upstream of the *lacUV5 O1* operator. Results are shown for LacI*-apCC-Tet* (underlined in red) and for controls LacI*-apCC-Tet and LacI*-CC-Di. GFP fluorescence was normalized to the OD_600_ of the cell culture and is an average of three repeats shown with standard error.

In an attempt to access an unfolding transition for one of the Q@***g*** designs, we measured CD spectra of the pQLI peptide in guanidinium hydrochloride, Gn·HCl. Surprisingly, neither the equilibrium spectra recorded at 5 °C nor the mean residue ellipticity at 222 nm (MRE_222_) signal recorded over 5 – 95 °C changed appreciably in the range of 0 – 6 M Gn·HCl (Figure S17-S18). Thus, pQLI is another hyperstable *de novo* peptide assembly. To probe this further, we truncated both pQLL and pQLI to 3-heptad repeats, yielding pQLL^3^ and pQLI^3^, respectively (Table 1). The overall charge pattern was preserved, though only the first and the last heptads had charged residues at ***b*** and ***c*** positions and the central repeat had Gln at these sites. Both truncated designs retained stable a-helical structures by equilibrium and variable-temperature CD measurements (Figures 2A and 2B). However, reducing the peptides concentrations to 5 μM accessed reversible thermal unfolding transitions, which were sigmoidal indicative of cooperativity, with estimated midpoints of 91 °C and 76 °C for pQLI and pQLL, respectively (Figure S19). Moreover, tetrameric assemblies for both peptides were confirmed by SV and SE experiments in AUC consistent with the target assemblies (Figure 2C and Figures S20-S21).

Next, we screened the 3- and 4-heptad variants of pQLL and pQLI for crystallization. Interestingly, only the pQLI peptides yielded crystals (Table S3). Both peptides gave good-quality X-ray diffraction data. These allowed structures to be determined by molecular replacement using ideal a helices implemented in Fragon^71^ for pQLI, or using the AlphaFold2^67– 69^ model for pQLI^3^ to resolutions of 0.96 and 1.42 Å, respectively (Figures 3A&B and Table S4). The solved structures confirmed the pQLI designs as antiparallel 4-helix CCs with knobs-into-holes packing identified by SOCKET2 (Table S5).^72, 73^ Inspection of a one-heptad slice through either structure (Figure 3C) immediately illustrates this packing and highlights the selection rules used in the design, namely: i) a *core* of Leu@***a*** that pack into holes on neighboring helices; ii) a *wide helix-helix interface* formed by the bulky Ile@***d*** residues and flanked by Gln@***g***; and a *narrow helix-helix interface* with Ala@***e*** allowing close helical contacts consistent with Alacoils^33, 66^ and flanked by Glu@***b***➔Lys@***b’*** salt bridges. In the two narrow interfaces of pQLI, 4 such salt bridges are made with C_δ_ ➔ N_ζ_ distances of 3.5 Å. Finally, the new X-ray crystal structures aligned closely with AlphaFold2 model for both pQLI analogues (RMSD_all-atom_ = 0.359 Å and 0.584 Å for pQLI and pQLI^3^, respectively, Figure S22).

**Figure 3.**
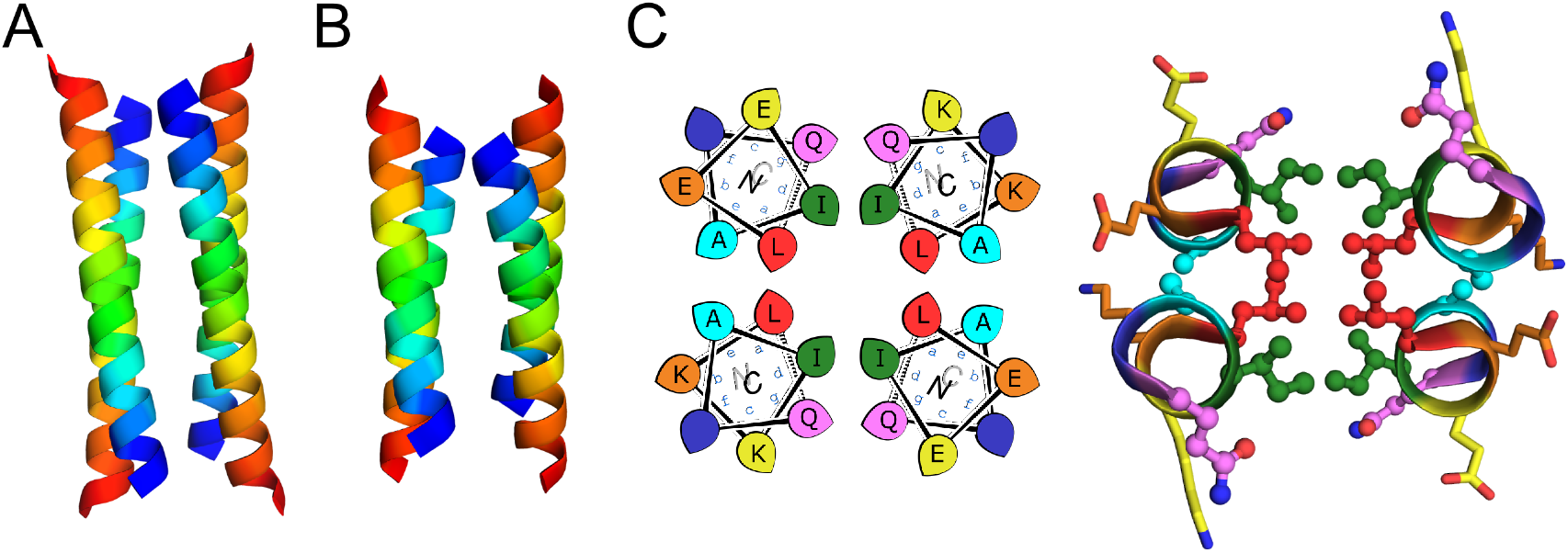
X-ray crystal structures for the antiparallel homotetrameric assemblies of pQLI analogues. (A) pQLI (apCC-Tet*, PDB ID: 8a3g) with 4 heptad repeats. (B) The shorter pQLI^3^ (apCC-Tet*^3^, PDB ID: 8a3i). The chains of both structures are colored in chainbow from the *N* (blue) to the *C* termini (red). (C) Left: helical wheel for the heptad repeats of pQLI. Right: slice through a heptad of the X-ray crystal structures for pQLI. Each position of the heptad is depicted in different color following the SOCKET2 scheme. Amino acids that compose the design rules are depicted in ball-and-stick representation.

We posit that the new designs with their clear and interpretable sequence-to-structure relationships offer stable modules for future applications in protein design and for chemical and synthetic biology. Therefore, we rename pQLI as apCC-Tet* to add to our basis set of robust and fully characterized *de novo* CCs. To demonstrate its potential utility, next we developed the design in a number of different assemblies as described below. The pQLL sequences were not taken forward from this point.

### apCC-Tet* assembles efficiently and functions in E. coli

To investigate the portability of apCC-Tet* into cells, we tested it as a component in an established transcriptional assay based on the oligomeric Lac repressor, LacI, in *Escherichia coli* (*E. coli*).^53^ In this assay, the repressor targets the *lac* promoter to control expression of a GFP reporter gene introduced on a plasmid: competent LacI complexes bind the promoter and repress the GFP gene (Figure 2D). We used a monomeric Lac repressor variant, LacI*, in which the wild-type (WT) tetramerization domain is removed, and the LacI dimer interface is disrupted.^74, 75^ We have demonstrated previously that *de novo* designed CCs can substitute for the natural oligomerization domain, which is an antiparallel homotetramer.^41, 53^ LacI* does not repress GFP production when expressed at low levels. However, when apCC-Tet* was fused to the *C* terminus of LacI*, repression was restored (Figure 2D). Moreover, the level of repression achieved was comparable to the parent LacI and the previous apCC-Tet design.^33, 41^ As WT LacI is a dimer of dimers, it can contact one or two *lacO1* operator sites in the promoter and, with the latter, repression is extremely tight. Importantly, we found that the level of repression induced by LacI*-apCC-Tet* was much greater with two *lacO1* operators in the reporter plasmid than with a single copy, indicating that tetramerization of the LacI* fusion protein occurred (Figure 2D). This contrasts with a control, where LacI* is fused to the dimeric CC-Di,^20^ which gave similar levels of GFP repression with one or two operators present. Overall, these data indicate that the designed apCC-Tet* efficiently tetramerizes in *E. coli* to restore the fully active LacI* complex and its DNA-binding function.

### apCC-Tet* can be adapted to build antiparallel heterotetramers

Breaking the symmetry of *de novo* CC homo-oligomers to make heteromeric systems has many advantages and potential applications.^37, 52, 76^ We sought to expand the utility of apCC-Tet* by redesigning it to make an A2B2 heterotetramer, apCC-Tet*-A_2_B_2_. For this, we maintained the ***g, a, d*** and ***e*** sites as Gln, Leu, Ile, and Ala, respectively, and made two potentially complementary peptides: an acidic peptide, apCC-Tet*-A, with Glu at all ***b*** and ***c*** positions; and a basic peptide, apCC-Tet*-B, with Lys at those sites. The only other change was the subtle use of Trp or Tyr at one ***f*** position in the A and B peptides, respectively. These designs were made in two lengths of 3 and 4 heptads to give two potential pairings, apCC-Tet*^3^-A_2_B_2_ and apCC-Tet*-A_2_B_2_ (Tables 1 and S2). The four peptides were synthesized by SPPS, purified, verified by mass spectrometry (Figures S23-S26), and characterized alone and as equimolar paired mixtures as follows.

Equilibrium and variable-temperature CD spectra revealed that the individual 4-heptad acidic and basic peptides were both folded and stable in PBS (Figures S27), and AUC-SV experiments showed that these isolated peptides formed tetramers like the parent homo-assembly despite the lack of complementary charges (Figures S28). An equimolar mixture of apCC-Tet*-A and apCC-Tet*-B spontaneously aggregated. Annealing the sample by heating up to 90 °C and then slowly cooling at room temperature resulted in soluble complexes, which were characterized as a folded and stable heterotetramer (Figures S27-S28). However, the annealed mixture had a lower a-helical content than the respective isolated peptides. Overall, these properties are far from ideal for a *de novo* designed module that can be used in other contexts and applications. Therefore, we turned to the 3-heptad pair, apCC-Tet*^3^-A plus apCC-Tet*^3^-B. Although fully or partly folded (Figure 4A), the individual acidic and basic peptides had accessible thermal unfolding transitions with midpoints of 61 °C and 42 °C, respectively (Figure 4B). *N.B*. The helicity of the basic peptide increased upon cooling back to below 20 °C. When mixed at 20 °C, the acid and basic peptides formed a partly helical assembly (Figure 4B). Moreover, upon heating between ≈40 – 55 °C, the mixture folded to a more-helical and hyperthermally stable assembly without an observable melting transition up to 95 °C (Figure 4B). AUC-SV experiments of annealed samples confirmed the presence of monodispersed tetramers in solution, consistent with an apCC-Tet*^3^-A2B2 design (Figure S29).

**Figure 4.**
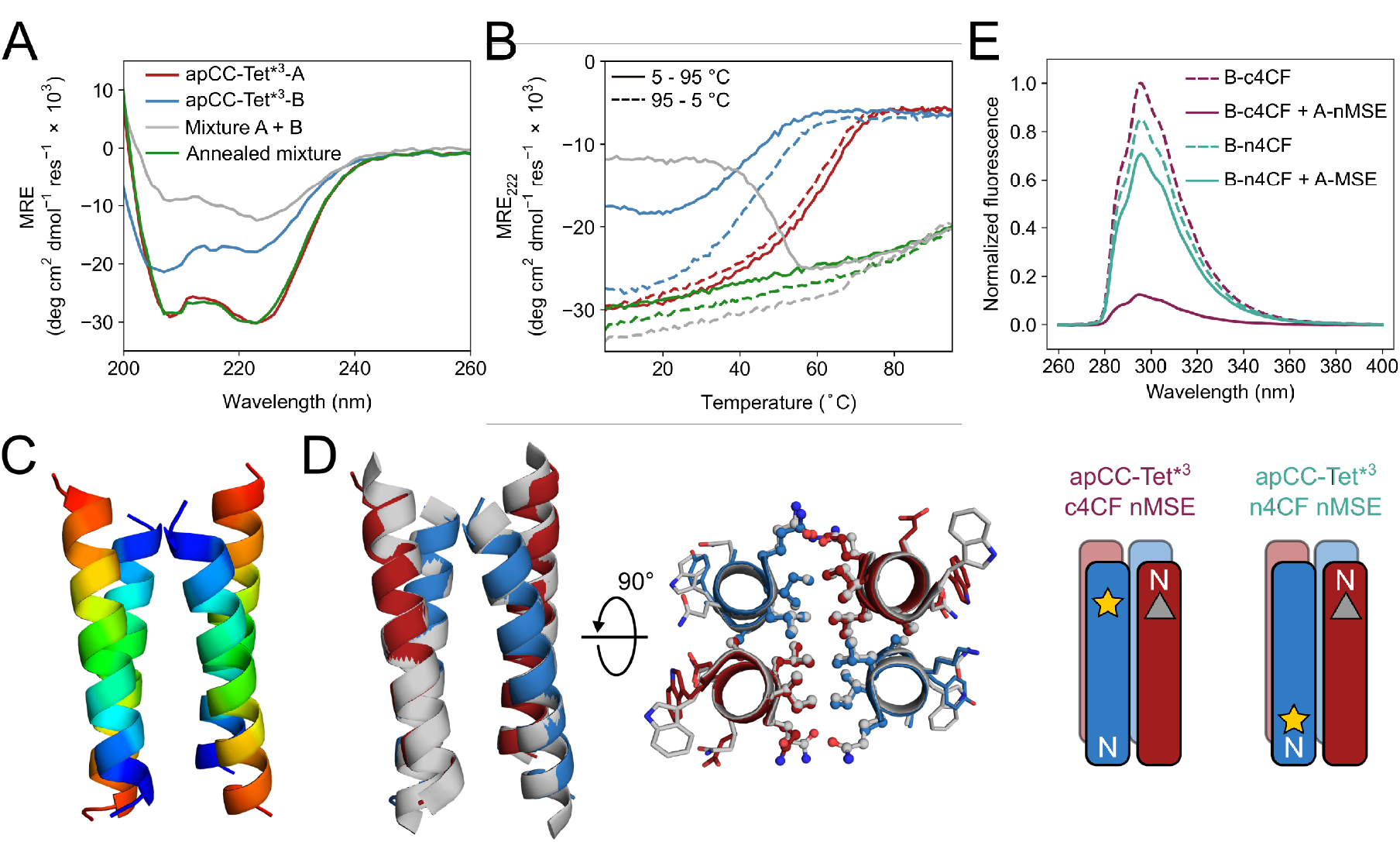
Biophysical and structural characterization of the heterotetrametric complex apCC-Tet*^3^-A_2_B_2_. (A) CD spectra at 5 °C and (B) thermal response curves (ramping up, solid lines; and ramping down, dashed line) for apCC-Tet*^3^-A (red), apCC-Tet*^3^-B (blue), the pre-annealed mixture apCCTet*^3^-A_2_B_2_ (grey), and the annealed mixture apCCTet*^3^-A_2_B_2_ (green). Conditions: 50 μM peptide, PBS, pH 7.4. (C) X-ray crystal structure of the heteromeric assembly apCCTet*^3^-A_2_B_2_ (PDB ID: 8a3j) with the chains colored from the *N* (blue) to the *C* termini (red). (D) Alignment of the crystal structures of apCCTet*^3^-A_2_B_2_ (apCC-Tet*^3^-A, red; apCC-Tet*^3^-B, blue) and the related homotetramer apCC-Tet*^3^(grey). (E) Fluorescence-quenching assay for labelled apCCTet*^3^-A_2_B_2_ peptides. 4CF is the 4-cyano-L-phenylalanine fluorophore (yellow star) and MSE is the L-selenomethionine fluorescence quencher (grey triangle). ‘n’ and ‘c’ indicate mutations near the *N* and *C* termini, respectively. In this panel only, peptide names are shortened for clarity. Conditions: 50 μM concentration of each peptide in phosphate buffer (8.2 mM sodium phosphate dibasic, 1.8 mM potassium phosphate monobasic), pH 7.4.

We were able to crystallize a mixture of apCC-Tet*^3^-A and apCC-Tet*^3^-B near neutral pH, and to obtain X-ray diffraction data out to 2.1 Å resolution (Table S3-S4). The resulting crystal structure revealed an antiparallel hetero-tetramer confirming the target apCC-Tet*^3^-A_2_B_2_ complex (Figure 4C). Like those for apCC-Tet* and apCC-Tet*^3^, the structure of apCC-Tet*^3^-A_2_B_2_ had a well-packed hydrophobic core with narrow and wide interfaces. Indeed, the heterotetramer overlaid well with the 3-heptad homotetramer (RMSD_all-atom_ = 0.390 Å, Figure 4D). Again, the design rules—***a*** = Leu, ***d*** = Ile, ***e*** = Ala, and ***g*** = Gln—are readily identifiable from visual inspection of the structure (Figure 4D).

Despite this experimental structure revealing an antiparallel orientation, there is one potential issue in moving from the ‘bar-magnet’ charge pattern of the homomeric system to the all-acidic plus all-basic design of the hetero-tetramer: the latter opens the possibility of accessing a parallel arrangement of helices in solution. To test this, we probed the arrangement of the assembled helices in solution using fluorescence-quenching experiments introduced by Raleigh.^77^ Guided by the X-ray crystal structure, we inserted the fluorescent 4-cyanophenylalanine (4CF) at the *C*-terminal ***e*** site of the B peptide to give apCC-Tet*^3^-B-c4CF (Table S2, Figure S30) and its quencher, selenomethionine (MSE), at the *N*-terminal ***b*** position of the A peptide (apCC-Tet*^3^-A-nMSE, Table S2, Figure S31). As a control, we also introduced the 4CF residue at the *N-*terminal ***c*** position of the B peptide (apCC-Tet*^3^-B-n4CF, Table S2, Figure S32), which should be too distant from the MSE residue for quenching in an antiparallel assembly with apCC-Tet*^3^-A-nMSE. Indeed, this control combination fluoresced comparably to the apCC-Tet*^3^-B-n4CF peptide alone (Figure 4E). Conversely, fluorescence was substantially quenched when apCC-Tet*^3^-B-c4CF was mixed with apCC-Tet*^3^-A-nMSE, indicating that the 4CF and the MSE groups were proximal, and confirming the formation of antiparallel tetramers in solution (Figure 4E).

### Constructing a single-chain de novo proteins from apCC-Tet*

As a final demonstration of the utility of the apCC-Tet* system in protein design, we targeted the construction of a single-chain protein that could be expressed from a synthetic gene in *E. coli*. This route of taking peptides into full-length proteins — *from peptides to proteins* — has been demonstrated for some time^14, 30, 78^ and discussed recently.^79–81^ The advantages of revisiting this approach are: i) that well-understood *de novo* peptide assemblies like apCC-Tet* should provide a strong basis for constructing robust *de novo* proteins; and ii) that the resulting proteins should be functionalizable through mutations to break symmetry whilst maintaining the design specification. Therefore, encouraged by the consistency of the apCC-Tet* peptides and the clear design rules underpinning them, we sought to make a singlechain 4-helix protein by looping together peptide sequences of the apCC-Tet* tetramer.

We hypothesized that helix packing would drive folding of the single-chain protein with only minor influences from the loops, and that no extensive design of the latter should be required. Therefore, we searched for loop sequences of reasonable composition that matched distances between the termini of the helices in the apCC-Tet* structure, while avoiding extended structures that can have unfavorable entropy contribution in the folding.^82, 83^ From the apCC-Tet* structure, we calculated end-to-end inter-helix distances of 17.4 – 18.5 Å and 12.5 – 15.0 Å for the wide and narrow faces, respectively. We treated these distances similarly to find loops in the PDB and from the literature to span both interfaces. The selected loops^70, 82, 84^ were arbitrarily incorporated into apCC-Tet*, but Alphafold2 predictions indicated that the sequence (Table 1) should form the desired single-chain 4HB (Figure S41). We called this single-chain protein sc-apCC-4.

A synthetic gene for sc-apCC-4 was expressed in *E. coli*, and the protein product was purified in sodium phosphate buffer (Figures S42-S43). Biophysical characterization by CD spectroscopy showed a highly α-helical structure that was fully resistant to thermal denaturation like the parent apCC-Tet* peptide (Figures 5A&B). Moreover, sc-apCC-4 was hyperstable to chemical denaturation, i.e., up 6 M Gn·HCl (Figure S44-S45). AUC-SV and SE experiments indicated that the *de novo* protein was a monodispersed monomer in solution (Figures 5C and S46). Finally, an X-ray crystal structure for sc-apCC-4 was obtained at 2.0 Å resolution. The structure was solved by molecular replacement using apCC-Tet* as starting model. It confirmed a monomeric four-helix CC bundle with an antiparallel (up-down-up-down) topology (Figure 5D). The sc-apCC-4 structure is consistent with all of our designs in this series: it has a well-packed hydrophobic core, wide and narrow faces, and the sequence-to-structure relationships are clear from visual inspection (Figure 5E). Moreover, and interestingly, it overlaid extremely well with the AlphaFold2 model with all all-atom RMSD of 0.475 Å (Figure S47). This suggests that core packing drives the folding over the loops demonstrating that design rules for apCC-Tet* are robust and transposable to build larger and well-defined proteins with analogous biophysical and structural properties.

**Figure 5.**
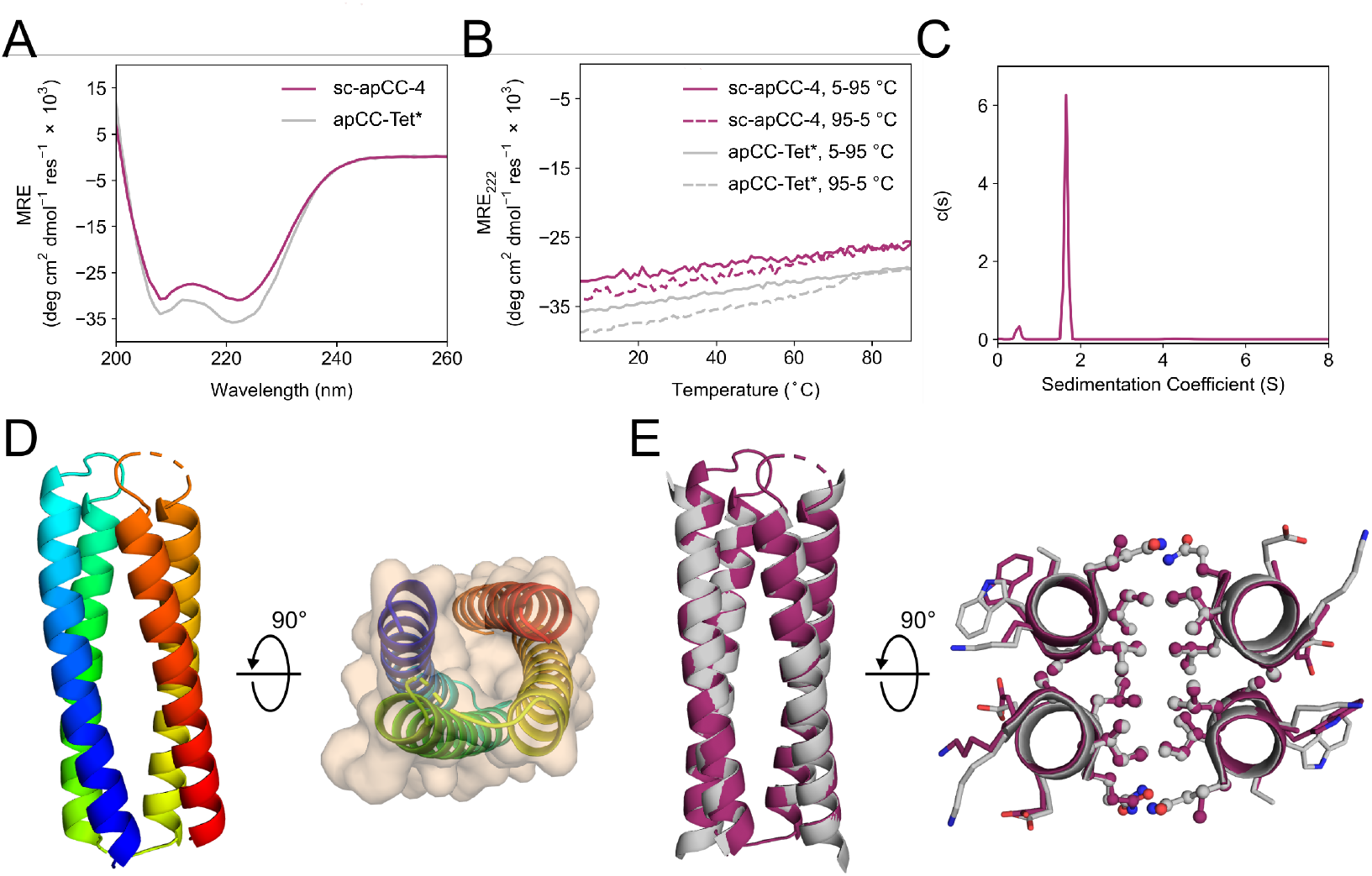
Characterization of the single-chain *de novo* protein, sc-apCC-4. (A) CD spectra at 5 °C and (B) thermal response curves (ramping up, solid lines; and ramping down, dashed line) for sc-apCC-4 (purple) in comparison with apCC-Tet* peptide (grey). Conditions: 25 μM protein in 50 mM sodium phosphate, 150 mM NaCl, pH 7.4 for the single-chain analogue; and 50 μM peptide, PBS, pH 7.4 for apCC-Tet*. (C) Sedimentation-velocity data from AUC for sc-apCC-4. The fit returned a weight of 0.9 × monomer mass. Conditions: 25 μM protein in 50 mM sodium phosphate, 150 mM NaCl, pH 7.4. (D) Left: X-ray crystal structure of sc-apCC-4 (PDB ID: 8a3k) colored chainbow from the *N* (blue) to the *C* terminus (red). Right: sc-apCC-4 structure viewed from the termini with chainbow coloring and surface representations. (E) Orthogonal views of the overlay between the structures of sc-apCC-4 (purple) and apCC-Tet* (grey) with a RMSDall-atom of 0.447 Å.

## CONCLUSION

We have combined bioinformatic analysis and rational protein design to determine a set of rules for the design of antiparallel four-helix coiled-coil bundles. Specifically, the rules are Gln@***g***, Leu@***a***, Ile@***d***, and Ala@***e*** in the heptad repeats, ***abcdefg***, of coiled-coil sequences. Using these, we have built a new homotetramer, apCC-Tet*, with ‘bar-magnet’ patterning of charged residues at ***b*** and ***c*** to help direct antiparallel helices. apCC-Tet* is hyperstable with respect to heat and chemical denaturation, and to truncation down to 3 heptad repeats. We have also used the rules to design heterotetramers comprising two different peptide chains, one with completely acidic residues and the other with basic residues at the ***b*** and ***c*** sites. Thus, in apCC-Tet*-A_2_B_2_, only the ***g***, ***a***, ***d*** and ***e*** sites are needed to direct antiparallel assembly. Finally, we show that the apCC-Tet* sequence can be concatenated to construct a single-chain 4-helix coiled-coil protein, sc-apCC-4, with loops taken from the PDB or the literature, and which can be expressed recombinantly in *E. coli*. All of the designs are fully characterized experimentally in solution and to atomic resolution by X-ray crystallography. Simple visual inspection of the resulting structures reveals that the design rules can be *read* straight from these structures. Thus, the rules are interpretable, robust, and transferable.

From the success of this rational approach, we contend that we now understood the contribution made by each amino acid in our designed sequences for 4-helix bundles. In turn, we anticipate that the newly designed peptides and protein will provide robust modules for further protein design to introduce function; and in chemical and synthetic biology as synthetic oligomerization domains. Such studies will be facilitated by the biophysical and structural characterizations that we provide here. Moreover, the different designs—of homo- and heterotetrameric peptides, and a monomeric protein—present opportunities to target and fine-tune different functions and uses. As an example of this potential, the relatively large and well-defined hydrophobic cores of tetrameric coiled coils and 4-helix bundles have been exploited by others to introduce cavities, small-molecule-binding pockets, and catalytic functionalities.^37, 39, 85–87^ Moreover, because our designed peptides and protein assemble efficiently in cells, such as *E. coli*, we anticipate applications to intervene in and to augment natural sub-cellular processes.^10, 53, 58, 62, 88, 89^

In short, we believe that our work adds fundamental understanding of the structural principles and sequence-to-structure relationships for coiled coils generally and 4-helix bundles specifically; and that our new designs provide platforms for future *de novo* design, and chemical and synthetic biology programs.

Of course, many others have designed *de novo* 4-helix bundles and coiled coils over the past four decades.^1, 4^ These have been achieved by modifying natural protein domains (e.g., the GNC4 leucine zipper, and the tetramerization domain of the Lac repressor),^19, 90, 91^ through rational approaches that focus on designing amphipathic helices,^18, 29, 30^ and by taking computational approaches.^26, 27, 70^ This has led to many different sequences for similar design targets. Therefore, to place our work in this broader context and to explore the potential sequence variation for the target, we examined other engineered and *de novo* designed sequences that (i) have been confirmed with high-resolution structures, and (ii) contain knobs-into-holes packing as detected by SOCKET2 (Table S6).^73^ Interestingly, we found that most of the foregoing sequences have no clear residue fingerprints at the ***g***, ***a***, ***d*** and ***e*** sites that we have focused on. Thus, there is no discernible consensus from these sequences. Those with the most regular hydrophobic cores and most similarity to our own designs are based on Harbury’s GCN4-pLI sequence.^19^ These have Leu@***a*** and Ile@***d***, but less regularity at the flanking ***e*** and ***g*** positions, which can be Leu, charged, or other residues (Table S6).^60^ Clearly, these and the other sequences ‘work’ and are solutions to the 4-helix-bundle design problem. However, we suggest that the heterogeneity in sequences and the lack of pinpointable sequence-to-structure relationships may make them less attractive as robust and mutable modules for future redesign and design studies.

Finally, it is interesting to speculate on the broader implications and applications of the approach of transforming self-assembling peptides to single-chain proteins as we demonstrate here and others have elsewhere.^18, 30, 78^ This can be likened to a possible evolutionary process in which primitive proteins might have assembled from the association and subsequent concatenation of smaller peptides,^80^ similar to the oligomerization of apCC-Tet* peptide to form robust tetramer and then the single-chain protein. The ease of looping the four helices together while maintaining the core folding provides some support to such a mechanism.^92^ Our future research aims to apply this approach to transform other well-understood multi-chain *de novo* coiled-coil peptides^20, 48, 49^ into single-chain proteins with clear sequence-to-structure features. We anticipate that the resulting synthetic proteins will be robust and stable, and, therefore, highly mutable to allow the incorporation of residues for binding, catalysis, and other functions.^36, 51, 52, 78, 88, 93, 94^

## ACKNOWLEDGMENTS

EAN, AJS, NJS and DNW are supported by a Biotechnology and Biological Sciences Research Council (BBSRC) grant (BB/S002820/1). KIA, ODW and DNW are supported by a BBSRC-NSF grant (BB/V004220/1 and 2019598). BM and DNW are supported by a BBSRC grant (BB/V006231/1). We are also grateful to the Max Planck-Bristol Centre for Minimal Biology, which supports KIA, BM, and DNW. DNW was also supported by BrisEngBio, a BBSRC-funded Engineering Biology Research Centre (BB/L01386X/1), and a Royal Society Wolfson Research Merit Award (WM140008). ODW is grateful for a National Institutes of Health grant (GM-118167). We thank the University of Bristol, School of Chemistry, Mass Spectrometry Facility for access to the EPSRC-funded Bruker Ultraflex MALDI-TOF instrument (EP/K03927X/1) and to the Synapt G2S nanospray instrument. We would like to thank Diamond Light Source for access to beamlines I04 and I24 (Proposal mx23269) and the European Synchrotron Radiation Facility (ESRF) for access to beamline ID30B (Proposal mx2373). We thank Will Dawson, Prasun Kumar, Freddie Martin, and members of the Woolfson laboratory for helpful discussions.

## TOC GRAPHIC

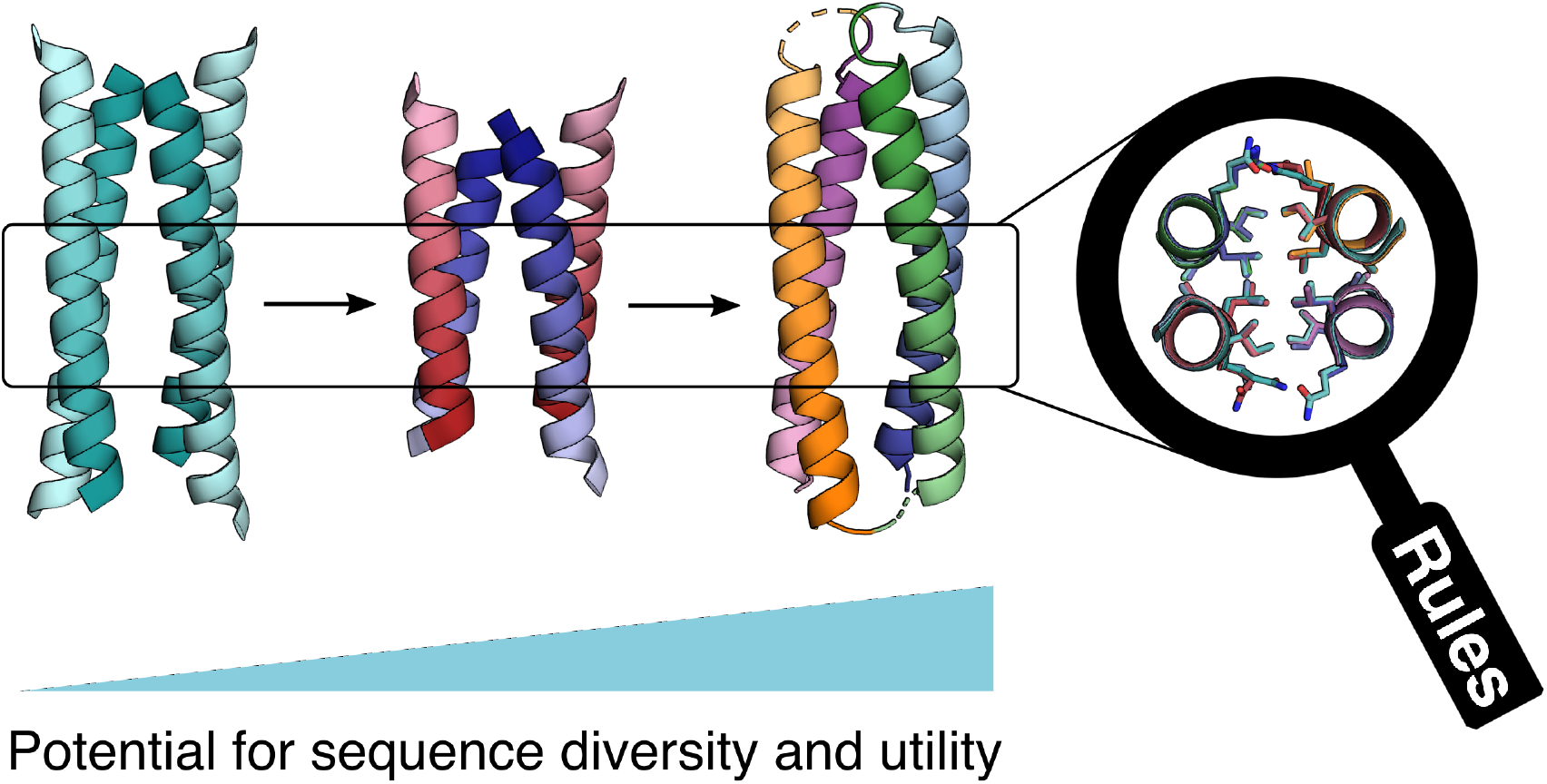

