## Supporting information for "From peptides to proteins: coiled-coil tetramers to single-chain 4-helix bundles"

### 1. Methods

#### 1.1. General

All solvents, chemicals, and reagents were purchased from commercial sources and used without further purification. Fluorenylmethoxycarbonyl(Fmoc)-α-L-amino acids, Rink amide MBHA resin for solid-phase peptide synthesis (SPPS) and N,N-dimethylformamide (DMF) were purchased from Sigma-Aldrich, Fisher Scientific and Cambridge Reagents. Coupling reagents Oxyma Pure and diisopropylcarbodiimide (DIC) were purchased from Fluorochem and Sigma-Aldrich, respectively. Pyridine was purchased from Fisher Scientific; triisopropylsilane (TIPS) was from Acros Organics. Morpholine and trifluoroacetic acid (TFA) as well as all other chemicals (reagent grade) were purchased from Sigma-Aldrich.

#### 1.2. Computational tools

***CC+ search***

Antiparallel coiled-coil tetramers and bundles were analyzed through the online CC+ database.[1] Four-helix coiled coils (CCs) with an antiparallel orientation, homo- or hetero-oligomeric partnering, canonical or non-canonical sequence repeats and comprising many chains of more than 11 residues, were searched with a sequence redundancy cut-off of ≤ 50%. From the database updated the 9$^{th}$ of January 2020, 65 coiled coils were assigned from 179 $\alpha$-helical sequences. Amino-acid profiles were compiled for each position of the heptad repeat, ***abcdefg*** (raw counts in Table S1). The profile was normalized using expected amino-acid frequencies from SWISS-PROT to give propensities for each residue (Figure 2B).

Amino-acid frequencies from SWISS-PROT are the following: Ala (A) 8.25, Arg (R) 5.53, Asn (N) 4.06, Asp (D) 5.46, Cys (C) 1.38, Gln (Q) 3.93, Glu (E) 6.72, Gly (G) 7.07, His (H) 2.27, Ile (I) 5.91, Leu (L) 9.65, Lys (K) 5.80, Met (M) 2.41, Phe (F) 3.86, Pro (P) 4.74, Ser (S) 6.64, Thr (T) 5.35, Trp (W) 1.10, Tyr (Y) 2.92, Val (V) 6.86.

***AlphaFold2 prediction***

AlphaFold2-multimer predictions were performed using the version implemented in ColabFold.[2-4] Different oligomeric states (4, 6, 8 and 12) were tested for each sequence in order to assess if the tetrameric assembly was still predicted. Predicted structures were relaxed using the implemented Amber force fields. Predicted IDDT per position and the predicted alignment error (PAE) were collected to evaluate the predictions.

#### 1.3. Peptide synthesis and purification

***Automated microwave Fmoc/tBu solid-phase peptide synthesis (SPPS)***

Automated microwave SPPS was performed on a Liberty Blue (CEM) synthesizer with in-line UV monitoring. The synthesis was performed on a 0.1 mmol scale on Rink amide MBHA (0.65 mmol/g loading, 100-200 mesh) resins. Side-chain protections of the amino acids were as follows: Gln(Trt), Glu(O*t*Bu), Lys(Boc), Tyr(*t*Bu), Trp(Boc). The coupling of the amino acids was performed in DMF by adding 5 equivalents of amino acids (2.5 mL, 0.2 M), 10 equivalents of DIC in DMF (1.0 mL, 1 M) and 5 equivalents of Oxyma Pure in DMF (1 mL, 0.5 M). Standard couplings were performed at 90 °C for 4 min (100 W for 20 s, 60 W for 10 s, 35 W for 240 s). Standard deprotections were performed using 20% (v/v) morpholine in DMF at 90 °C for 1 min (125 W 30 s, 32 W 60 s). All peptides were manually capped with *N*-terminal acetyl groups through addition of pyridine (0.5 mL) and acetic anhydride (0.25 mL) in DMF (9.25 mL), with shaking at room temperature (rt) for 20 minutes. The resin was washed 3 times with DMF followed by 6 times with DCM before cleavage. Peptides were cleaved from the resin and deprotected with addition of 10 mL of a mixture 95:2.5:2.5 v/v/v TFA/$H_2O$/TIPS, with shaking at rt for 2 hours. The SeMet-containing peptide was cleaved from the resin with addition of 10 mL of a mixture 88.7:5:3.3:3 v/v/v/v TFA/thioanisole/anisole/1,2-ethanedithiol (EDT), with shaking at rt for 2 hours. The TFA solution was filtered to remove the resin beads and was reduced in volume to ≤5 mL using a flow of $N_2$. Cleaved peptide was precipitated with cold diethyl ether (≈45 mL), isolated via centrifugation and dissolved in a 1:1 mixture MeCN/$H_2O$. Crude peptides were lyophilized to yield white powders.

***Semi-preparative High Performance Liquid Chromatography (HPLC)***

All peptides were purified by reverse phase HPLC (JASCO) using a Luna C18 (Phenomenex) column (150 x 10 mm, 5 µM particle size, 100 Å pore size). Crude peptide was dissolved at 7 mg/mL in 20% v/v MeCN in $H_2O$ with 0.1% TFA, injected to the column and eluted with a 3 mL/min linear gradient (20-60%) of MeCN in $H_2O$ with 0.1% TFA each over 30 minutes. Alternatively, acidic peptides were eluted with a linear gradient (20-60%) of MeCN in 25 mM $NH_4HCO_3$ in $H_2O$ over 30 min. Elution of the peptide was detected with in-line UV monitoring at

220 nm and 280 nm simultaneously. When required, a column oven (50 °C) was used to improve separation. Pure fractions were identified by analytical HPLC and matrix-assisted laser desorption/ionization–time of flight (MALDI-TOF) mass spectrometry, then pooled, and freeze-dried.

***Analytical HPLC***

Analytical HPLC traces were obtained using a Jasco 2000 series HPLC system and a Phenomenex Kinetex C18 (100 x 4.6 mm, 5 µm particle size, 100 Å pore size) column. Chromatograms were monitored at 220 nm and 280 nm. The linear gradient was 20–80% MeCN in water (each containing 0.1% TFA) over 25 min at a flow rate of 1 mL/min. Alternatively, acidic peptides were analyzed with the same linear gradient of MeCN but with 25 mM $NH_4HCO_3$ in $H_2O$. When required, a column oven (50 °C) was used to assist peptide elution.

***Mass spectrometry***

Matrix-assisted laser desorption/ionization–time of flight (MALDI-TOF) mass spectra were collected on a Bruker UltraFlex MALDI-TOF mass spectrometer operating in positive-ion reflector mode. Peptides were spotted on a ground steel target plate using $\alpha$-cyano-4-hydroxycinnamic acid dissolved in 1:1 MeCN/$H_2O$ as the matrix. Masses quoted are for the monoisotopic mass as the singly protonated species. Alternatively, full electrospray ionization (ESI) MS spectra were acquired on a Synapt G2S (Waters) mass spectrometer equipped with an IMS-Q-TOF analyzer and using an Advion Nanomate for robot chip-based nanospray ionization in positive mode. Some acidic peptides were analyzed in negative mode. 5 µL of a 50 µM peptide solution in 1:1 MeCN/$H_2O$ were generally injected for the analysis. Masses quoted are for the deconvoluted monoisotopic mass.

#### 1.4. Protein expression and purification

***Cloning and protein sequence***

A synthetic gene for sc-apCC-4 cloned directly into pET28a vector using NdeI and XhoI restriction sites was purchased from Twist Biosciences.

<u>Sc-apCC-4 protein sequence:</u>

*MGSSHHHHHHSSGLVPRGSHM*QLEEIAQQLEEIAKQLKKIAWQLKKIAQGEPSAQGQLEEIAQ
QLEEIAKQLKKIAWQLKKIAQGPDSVQLEEIAQQLEEIAKQLKKIAWQLKKIAQGGTSGGQLEEIA
QQLEEIAKQLKKIAWQLKKIAQ

***Protein expression and purification***

The sc-apCC-4 gene was transformed and then recombinantly expressed in *E. coli* BL21-DE3 Gold (Agilent) cells. Flasks containing 1 L of LB-kanamycin (Thermo Scientific, 50 µg/mL) were inoculated with 5 mL overnight cultures and incubated to an OD600 of ~0.6 at 37 ˚C with 220 rpm shaking. Expression was induced with 0.5 mM IPTG (Neo Biotech), and cultures were incubated at 18 ˚C overnight with 220 rpm shaking (Thermo Scientfic MaxQ 4000).

Following expression, cultures were pelleted at 5000 rpm for 10 minutes (Thermo Scientific Sorvall Lynx 4000). Cell pellets were resuspended in 20 mL lysis buffer (50 mM Tris, pH 7.4, 150 mM NaCl, 30 mM imidazole, 1 mg/mL lysozyme) for 30 min at 37 ˚C. Resuspended pellets were sonicated using a Biologics Model 3000 Ultrasonic homogenizer with settings to 50% power, 90% pulser (1 pulse/second) for five minutes, then clarified at 12,000 rpm for 1 hour. The expressed sc-apCC-4 was first purified with Ni-affinity chromatography. Filtered lysate was loaded onto an ÄKTAprime plus (GE Healthcare) equipped with a HisTrap- 5 mL HP column (GE Healthcare). His-tagged proteins were eluted at 55% Buffer B (Buffer A: 50 mM Tris, 150 mM NaCl, 30 mM imidazole, pH 7.4; Buffer B: 50 mM Tris, 150 mM NaCl, 300 mM imidazole, pH 7.4) using a step gradient. Fractions were combined and purified further by size exclusion chromatography using a HiLoad 16/600 Superdex 75 pg size exclusion column (GE Healthcare) equilibrated in buffer containing 50 mM sodium phosphate, pH 7.4, 150 mM NaCl. Eluted fractions were pooled, concentrated, diluted 1:1 in loading buffer (50mM Tris-Cl (pH 6.8), 10 mM DTT, 2% SDS, 0.1% bromophenol blue, 10% glycerol), and heated to 95 ˚C. 15 µL samples and PageRuler Low Range Unstained Protein Ladder (Thermo Scientific) were separated on a 10-20% SDS-PAGE gel using Tris-Glycine running buffer at 120V (BioRad PowerPac Basic) to confirm identity.

#### 1.5. Solution-phase biophysical characterizations

***Peptide and protein concentration determination***

Peptide and protein concentrations were determined from absorbance at 280 nm measured in a Nanodrop 2000 (Thermo Scientific) spectrometer ($\varepsilon_{280}$(Trp) = 5690 cm$^{-1}$; $\varepsilon_{280}$(Tyr) = 1280 cm$^{-1}$; $\varepsilon_{280}$(sc-apCC-4) = 22000 cm$^{-1}$), or at 214 nm using a Cary-100 (Agilent) UV-Visible spectrometer by measuring the peptide bond.[5]

***Sample annealing***

When necessary, mixtures of acidic and basic peptides were annealed before biophysical measurements. Peptide samples in buffer were heated up to 90 °C in a heat block (Grant), incubated at 90 °C for 10 min and slowly cooled down to room temperature over 90 min.

***Circular dichroism (CD) spectroscopy***

Circular dichroism (CD) data were collected in the far-UV region on a JASCO J-810 or J-815 spectropolarimeter fitted with a Peltier temperature controller. Peptide samples were made up as 50 μM peptide solution in phosphate buffered saline (PBS; 8.2 mM sodium phosphate dibasic, 1.8 mM potassium phosphate monobasic, 137 mM sodium chloride, 2.4 mM potassium chloride), pH 7.4 at 5 °C. For the protein sc-apCC-4, CD spectra were acquired at 25 μM protein concentration in 50 mM sodium phosphate, 150 mM NaCl, pH 7.4 at 5 °C. Data were collected in a 1 mm quartz cuvette between 190 and 260 nm and the instrument was set as follows: band width 1 nm, data pitch 1 nm, scanning speed 100 nm/min, 1 s response time. Every CD curve was obtained by averaging of 8 scans and subtracting the background signal of buffer and cuvette. For thermal profile experiments, CD signals were monitored at 222 nm wavelength. A temperature range of 5–95 °C and a temperature ramp rate of 60 °C per hour were used with the same settings and peptide or protein concentration as above.

CD data were converted from ellipticities (mdeg) to mean residue ellipticities (MRE, ($deg \cdot cm^2 \cdot dmol^{-1} \cdot res^{-1}$)) by normalizing for concentration of peptide bonds and the cell path length using the equation:

$$MRE = \frac{\theta \times 10^6}{c \times l \times n}$$

where the variable $\theta$ is the measured difference in absorbed in left- and right-handed circularly polarized light in millidegrees, $c$ is the micromolar concentration of the compound, $l$ is the path length of the cuvette in mm, and $n$ is the number of amide bonds in the polypeptide, for which the N-terminal acetyl bond was included but not the C-terminal amide.

<u>Chemical denaturation:</u>

The stability against chemical denaturation was examined in presence of the chaotropic agent guanidinium hydrochloride (Gn·HCl). 5 μM or 50 μM of apCC-Tet* or 1 μM of sc-apCC-4 were mixed with increasing concentration of Gn·HCl (0-6 M), and samples were incubated overnight. Data were collected in a 1 or 5 mm quartz cuvette at 5 °C between 190 and 260 nm (100 nm/min,

1 nm interval and bandwidth, 1 s response time). Variable-temperature spectra were collected at 222 nm for the highest concentration of Gn·HCl (i.e., 6 M) using the settings and peptide or protein concentration as above between 5 and 95 °C at a 60 °C/hour ramp rate.

***Analytical ultracentrifugation***

Analytical ultracentrifugation (AUC) was performed on a Beckman Optima X-LA or X-LI analytical ultracentrifuge with an An-50-Ti or An-60-Ti rotor (Beckman-Coulter). Buffer densities, viscosities, and peptide partial specific volumes ($\overline{v}$) were calculated using SEDNTERP (http://rasmb.org/sednterp/).

For sedimentation velocity (SV), sample solutions of 310 or 410 µL were prepared in PBS at 150 µM peptide concentration and placed in a sedimentation velocity cell with an epon or aluminium, respectively, 2-channel centerpiece and quartz windows. The reference channel was loaded with 320 µL (or 420 µL, respectively) of PBS buffer. For SV experiments with the sc-apCC-4 protein, samples were prepared at 25 µM peptide concentration in 50 mM sodium phosphate, pH 7.4, 150 mM NaCl. The samples were centrifuged at 50 krpm at 20 °C, with absorbance scans taken across a radial range of 5.8–7.3 cm at 5 min intervals to a total of 120 scans. Data from a single run were fitted to a continuous c(s) distribution model using SEDFIT,[6] at 95% confidence level. Residuals for sedimentation velocity experiments are shown as a bitmap in which the grayscale shade indicates the difference between the fit and raw data (residuals < -0.05 are black and residuals > 0.05 are white). Good fits are uniformly grey without major dark or light streaks.

Sedimentation equilibrium (SE) experiments were performed at 70 µM peptide concentration or 25 µM protein concentration in 110 µL at 20 °C. Experiments were run in triplicate in a six- channel epon centerpiece. The samples were centrifuged at speeds in the range of 18−48 krpm or 20-45 krpm in increments of 6 and 5 krpm respectively. Scans at each speed were duplicated 6 or 8 hours after the increase in speed to confirm equilibration. Data were fitted using SEDPHAT to a single species model.[7] Monte Carlo analysis was performed to give 95% confidence limits.

***Fluorescence quenching experiments***

Fluorescent quenching experiments were performed following previously published procedure.[8] Mixture were prepared as an equimolar ratio of acidic and basic peptide analogues at 50 µM of the CN-Phe-containing peptide in phosphate buffer (8.2 mM sodium phosphate dibasic, 1.8 mM potassium phosphate monobasic), pH 7.4. Fluorescence emission spectra from 260 nm to 400 nm were recorded on a Jasco Fluorimeter, in a 10 x 10 mm quartz cuvette, and with an excitation wavelength of 240 nm. Every fluorescence emission spectrum was obtained by averaging of 3 scans and subtracting the background signal of buffer and cuvette.

#### 1.6. *In vivo* characterization of peptide assembly

***Plasmids for GFP repression assays***

The plasmids for the GFP repression assays, pBADLacI, pBADLacI*, pBADLacI*-ccDi and pBADLacI*-apCC-Tet, were used as previously described.[9, 10] In order to make pBADLacI*-apCC-Tet*, a DNA fragment encoding apCC-Tet*, with codons optimized for *E. coli*, was designed and was pucharsed from Eurofins Genomics. The DNA fragment was cloned into pBADLacI* at Acc65I/XbaI restriction sites. These plasmids expressed WT LacI or a truncated dimerization mutant LacI* (LacI L251A aa 1-332) with terminal $His_6$, T7, and Xpress tags from the $P_{araBAD}$ arabinose-inducible promoter. CC peptides were fused in frame and C-terminal to LacI* by a flexible linker. The GFP reporter plasmids pVRbLacUV5 and pVRblacO1-lacO1 have also been described previously.[9]

***GFP repression assays***

In order to assay repression of GFP transcription, TB28 cells (MG1655ΔLacIZYA)[11] were transformed with either pVRbLacUV5 (1 lac operator site) or pVRblacO1-lacO1 (2 lac operator sites) and pBADLacI* or its derivatives. Repression assays were carried out essentially as described previously,[9] except that fluorescence intensity was measured using a Clariostar plus microplate reader (BMG Labtech) with GFP settings (i.e., monochromator/filter settings: 470-15/515-20).

#### 1.7. Structural characterization

***Crystal growth***

Diffraction-quality peptide crystals were grown using a sitting-drop vapor-diffusion method. Freeze-dried peptides were dissolved in ultrapure water and diluted to 10 mg/mL. Purified sc-apCC-4 in 50 mM sodium phosphate, 150 mM NaCl was concentrated, then diluted to 10 mg/mL in deionized water, in this case the final buffer concentration was 40 mM phosphate, 120 mM NaCl. Commercially available sparse matrix screens were used (Morpheus®, JCSG-plus™,

Structure Screen 1 and 2, Pact Premier™, ProPlex™; Molecular Dimensions), and the drops were dispensed using a robot (Oryx8; Douglas Instruments). The peptides were screened in MRC 2 drop plates. Drops consisting of 0.3 µL of peptide solution and 0.3 µL of reservoir solution were screened in parallel with drops consisting of 0.4 µL of the peptide solution and 0.2 µL of reservoir solution. For the sc-apCC-4, 0.4 µL of the protein solution and 0.2 µL of reservoir solution were mixed. All plates were incubated at 20 °C. Crystals generally formed within a month, and after looping were soaked in reservoir solution containing 25% glycerol as a cryoprotectant. Crystals of apCC-Tet$^{*3}$-$A_2B_2$ were obtained by optimization around the condition from Structure Screen 1 and 2 C5 comprising 1.4 M sodium citrate tribasic dihydrate, 0.1 M sodium HEPES, pH 7.5. A range of concentrations of sodium citrate tribasic dihydrate from 0.8 M to 1.8 M was screened as well as peptides concentrations (11 mg/mL, 8.5 mg/mL and 6.4 mg/mL). 0.5 µL of the peptide solution and 0.5 µL of reservoir solution were mixed in MRC Maxi 48 Well plates and incubated at 20 °C.

Final crystallization conditions for all peptides are provided in Table S2.

***X-ray crystal structure determination***

Diffraction data for the crystals were obtained at the Diamond Light Source on beamlines I04 or I24 (for apCC-Tet$^{*3}$-$A_2B_2$ and apCC-Tet*, respectively) or at the European Synchrotron Radiation Facility (ESRF) on beamline ID30B (for apCC-Tet$^{*3}$ and sc-apCC-4).

Data were processed using the automated pipelines: Xia2,[12] which ports data through DIALS[13] or MOSFLM[14] to POINTLESS and AIMLESS[15] as implemented in the CCP4 suite,[16] or XDS to XSCALE.[17] Structure of apCC-Tet$^{*3}$-$A_2B_2$ was phased using *ab initio* phasing using ARCIMBOLDO_LITE.[18] Structure of apCC-Tet* was phased using FRAGON,[19] the initial phases were into and refined using BUCCANEER[20] and REFMAC 5.[21] Structure of apCC-Tet$^{*3}$ and sc-apCC-4 were solved by molecular replacement using AlphaFold[2, 4] or apCC-Tet*, respectively, as models using PHASER.[22] Final structures were obtained after iterative rounds of model building with COOT and refinement with PHENIX Refine.[23] Solvent-exposed atoms lacking map density were either deleted or left at full occupancy. Data collection and refinement statistics are provided in Table S3.

### 2. Supplementary data

#### 2.1. Supplementary tables

**Table S1. Raw amino-acid counts for antiparallel four-helix coiled coils found by the CC+ database (over 179 α-helical sequences).**[a]

| - | *a* | *b* | *c* | *d* | *e* | *f* | *g* | Sum |
|---|---|---|---|---|---|---|---|---|
| **A** | 103 | 41 | 42 | 77 | **92** | 69 | 74 | 498 |
| **C** | 7 | 2 | 5 | 0 | 4 | 0 | 1 | 19 |
| **D** | 5 | 67 | 44 | 8 | 31 | 58 | 25 | 238 |
| **E** | 17 | 128 | 183 | 24 | 71 | 148 | 72 | 643 |
| **F** | 31 | 5 | 7 | 14 | 4 | 10 | 12 | 83 |
| **G** | 4 | 27 | 18 | 5 | 20 | 25 | 21 | 120 |
| **H** | 13 | 15 | 12 | 10 | 5 | 11 | 12 | 78 |
| **I** | 81 | 14 | 13 | **194** | 33 | 18 | 65 | 418 |
| **K** | 11 | 74 | 76 | 5 | 60 | 75 | 38 | 339 |
| **L** | **226** | 37 | 18 | **224** | 98 | 30 | **158** | 791 |
| **M** | 33 | 11 | 12 | 32 | 17 | 4 | 16 | 125 |
| **N** | 9 | 51 | 33 | 14 | 23 | 52 | 23 | 205 |
| **P** | 0 | 0 | 2 | 5 | 0 | 0 | 6 | 13 |
| **Q** | 21 | 66 | 68 | 33 | 58 | 55 | **67** | 368 |
| **R** | 8 | 82 | 70 | 6 | 77 | 75 | 39 | 357 |
| **S** | 40 | 37 | 40 | 36 | 56 | 43 | 46 | 298 |
| **T** | 36 | 25 | 31 | 22 | 37 | 39 | 34 | 224 |
| **V** | 126 | 17 | 17 | 82 | 43 | 13 | 39 | 337 |
| **W** | 0 | 0 | 4 | 3 | 2 | 5 | 8 | 22 |
| **Y** | 5 | 7 | 9 | 8 | 20 | 11 | 4 | 64 |
| **Sum** | 780 | 706 | 706 | 804 | 758 | 741 | 766 | 5240 |

[a]Residues identified for the design of the new antiparallel tetramer sequences are highlighted in red and bold.

**Table S2. Sequences of *de novo* peptides and proteins in this study.**

| Peptide/protein names | Sequences<br>***gabcdef gabcdef gabcdef gabcdef loop*** |
|---|---|
| pLLL | Ac-G LLEELAQ LLEELAK LLKKLAW LLKKLAQ G-$NH_2$ |
| pLLI | Ac-G LLEEIAQ LLEEIAK LLKKIAW LLKKIAQ G-$NH_2$ |
| pQLL | Ac-G QLEELAQ QLEELAK QLKKLAW QLKKLAQ G-$NH_2$ |
| pQLI<br>**apCC-Tet*** | Ac-G QLEEIAQ QLEEIAK QLKKIAW QLKKIAQ G-$NH_2$ |
| pQLL$^{3}$ | Ac-G QLEELAK QLQQLAW QLKKLAQ G-$NH_2$ |
| pQLI$^{3}$<br>**apCC-Tet*$^{3}$** | Ac-G QLEEIAK QLQQIAW QLKKIAQ G-$NH_2$ |
| pQLI-A<br>**apCC-Tet*-A** | Ac-G QLEEIAQ QLEEIAK QLEEIAW QLEEIAQ G-$NH_2$ |
| pQLI-B<br>**apCC-Tet*-B** | Ac-G QLKKIAQ QLKKIAK QLKKIAY QLKKIAQ G-$NH_2$ |
| pQLI$^{3}$-A<br>**apCC-Tet*$^{3}$-A** | Ac-G QLEEIAK QLEEIAW QLEEIAQ G-$NH_2$ |
| pQLI$^{3}$-B<br>**apCC-Tet*$^{3}$-B** | Ac-G QLKKIAK QLKKIAY QLKKIAQ G-$NH_2$ |
| pQLI$^{3}$-A-nMSE<br>**apCC-Tet*$^{3}$-A-nMSE** | Ac-G QLXEIAK QLEEIAK QLEEIAQ G-$NH_2$ |
| pQLI$^{3}$-B-c4CF<br>**apCC-Tet*$^{3}$-B-c4CF** | Ac-G QLKKIAK QLKKIAK QLKKIZQ G-$NH_2$ |
| pQLI$^{3}$-B-n4CF<br>**apCC-Tet*$^{3}$-B-n4CF** | Ac-G QLKZIAK QLKKIAK QLKKIAQ G-$NH_2$ |
| **sc-apCC-4** | QLEEIAQ QLEEIAK QLKKIAW QLKKIAQ GEPSAQG<br>QLEEIAQ QLEEIAK QLKKIAW QLKKIAQ GPDSV<br>QLEEIAQ QLEEIAK QLKKIAW QLKKIAQ GGTSGG<br>QLEEIAQ QLEEIAK QLKKIAW QLKKIAQ |
| pQLL-A | Ac-G QLEELAQ QLEELAK QLEELAW QLEELAQ G-$NH_2$ |
| pQLL-B | Ac-G QLKKLAQ QLKKLAK QLKKLAY QLKKLAQ G-$NH_2$ |
| pQLL$^{3}$-A | Ac-G QLEELAK QLEELAW QLEELAQ G-$NH_2$ |
| pQLL$^{3}$-B | Ac-G QLKKLAK QLKKLAY QLKKLAQ G-$NH_2$ |

X = Selenomethionine (MSE); Z = 4-cyanophenyalanine (4CF)

**Table S3. Crystallization conditions used to obtain the structures discussed in this article.**

| Peptide or protein | Molecular dimensions screen | Crystallization conditions of the reservoir | Dilution sample/reservoir (in the drop) |
|---|---|---|---|
| **apCC-Tet* (8A3G)** | Proplex, G2 | 2.0 M Ammonium sulfate, 0.1 M Sodium acetate, pH 5.0 | 1:1 |
| **apCC-Tet*$^3$ (8A3I)** | Structure Screen 1 and 2, C5 | 1.4 M sodium citrate tribasic dihydrate, 0.1 M sodium HEPES, pH 7.5 | 2:1 |
| **apCC-Tet*$^3$-$A_2B_2$ (8A3J)** | Optimization from Structure Screen 1 and 2, C5 | 1.8 M sodium citrate tribasic dihydrate, 0.1 M sodium HEPES, pH 7.5 | 1:1 |
| **sc-apCC-4 (8A3K)** | Morpheus, D9 | 10% w/v PEG 20 000, 20% v/v PEG MME 550, 0.02 M 1,6-hexanediol, 0.02 M 1-butanol, 0.02 M (RS)-1,2-propanediol, 0.02 M 2-propanol, 0.02 M 1,4-butanediol, 0.2 M 1,3-propanediol, 0.1 M bicine/Trizma base pH 8.5 | 2:1 |

**Table S4. Merging and refinement statistics for all X-ray crystal structures.**

| | apCC-Tet* | apCC-Tet*3 | apCC-Tet*3-$A_2B_2$ | sc-apCC-4 |
|---|---|---|---|---|
| **PDB ID** | 8A3G | 8A3I | 8A3J | 8A3K |
| **Data collection** | | | | |
| Source | Diamond I24 | ESRF ID30B | Diamond I04 | ESRF ID30B |
| Detector | PILATUS 6M | PILATUS3 6M | EIGER2 XE 16M | PILATUS3 6M |
| Wavelength (Å) | 0.85 | 0.9253 | 0.9795 | 0.9253 |
| Resolution range | 26.86 - 0.96 (0.9943 - 0.96) | 42.46 - 1.42 (1.471 - 1.42) | 31.58 - 2.1 (2.175 - 2.1) | 27.48 - 2.0 (2.071 - 2.0) |
| Space group | $P\ 4_2\ 2_1\ 2$ | $P\ 3_2\ 2\ 1$ | $P\ 1\ 2_1\ 1$ | $P\ 1\ 2_1\ 1$ |
| Unit cell: *a, b, c (Å)* | 34.01 34.01 87.53 | 49.03 49.03 32.62 | 27.66 63.16 47.30 | 28.08 38.86 56.83 |
| *α, β, γ (°)* | 90 90 90 | 90 90 120 | 90 91.6752 90 | 90 101.884 90 |
| Total reflections | 645077 (19488) | 28332 (2768) | 64987 (6508) | 18274 (1751) |
| Unique reflections | 31036 (2293) | 8690 (831) | 9430 (923) | 7660 (751) |
| Multiplicity | 20.8 (8.5) | 3.3 (3.2) | 6.9 (7.1) | 2.4 (2.3) |
| Completeness (%) | 95.51 (72.54) | 97.45 (93.53) | 98.43 (97.88) | 92.85 (92.36) |
| Mean I/sigma(I) | 30.55 (1.32) | 38.83 (2.30) | 4.37 (0.62) | 13.21 (1.29) |
| Wilson B-factor | 8.99 | 16.65 | 22.57 | 47.29 |
| R-merge | 0.0429 (0.7173) | 0.1061 (0.6371) | 0.2294 (1.152) | 0.03111 (0.8826) |
| R-meas | 0.04392 (0.7643) | 0.1268 (0.7589) | 0.2486 (1.243) | 0.03907 (1.101) |
| R-pim | 0.009184 (0.2529) | 0.06726 (0.4024) | 0.09484 (0.4646) | 0.02326 (0.6503) |
| CC1/2 | 1 (0.94) | 0.973 (0.666) | 0.986 (0.783) | 0.999 (0.602) |
| CC* | 1 (0.984) | 0.993 (0.894) | 0.997 (0.937) | 1 (0.867) |
| **Refinement** | | | | |
| Reflections used in refinement | 30910 (2280) | 8592 (809) | 9413 (922) | 7651 (749) |
| Reflections used for R-free | 1514 (114) | 427 (51) | 389 (42) | 345 (24) |
| R-work | 0.1603 (0.2220) | 0.1989 (0.3187) | 0.2297 (0.2879) | 0.2363 (0.3773) |
| R-free | 0.1688 (0.2054) | 0.2348 (0.4990) | 0.2672 (0.3055) | 0.2756 (0.3643) |
| CC(work) | 0.969 (0.939) | 0.498 (0.135) | 0.907 (0.814) | 0.969 (0.695) |
| CC(free) | 0.976 (0.946) | 0.752 (0.236) | 0.918 (0.848) | 0.987 (0.535) |
| Number of non-hydrogen atoms | 577 | 379 | 1299 | 823 |
| macromolecules | 533 | 370 | 1275 | 823 |
| ligands | 5 | 0 | 0 | 0 |
| solvent | 39 | 9 | 24 | 0 |
| Protein residues | 64 | 50 | 186 | 127 |
| RMS(bonds) | 0.017 | 0.010 | 0.009 | 0.006 |
| RMS(angles) | 1.50 | 1.08 | 1.06 | 0.83 |
| Ramachandran favored (%) | 100.00 | 100.00 | 99.38 | 96.69 |
| Ramachandran allowed (%) | 0.00 | 0.00 | 0.62 | 3.31 |
| Ramachandran | 0.00 | 0.00 | 0.00 | 0.00 |
| Rotamer outliers (%) | 0.00 | 0.00 | 3.09 | 0.00 |
| Clashscore | 0.90 | 2.68 | 4.84 | 4.52 |
| Average B-factor | 12.89 | 28.54 | 29.38 | 55.19 |
| macromolecules | 11.97 | 28.00 | 29.16 | 55.19 |
| ligands | 26.99 | - | - | - |
| solvent | 20.69 | 38.03 | 33.90 | - |
| Number of TLS groups | - | - | 2 | 4 |

**Table S5. Socket2 analysis of crystallographic structures and related AlphaFold2 predictions.[24]**

| Structures | | Number of observed knobs at different cutoffs | | | | | | | | | | | |
|---|---|---|---|---|---|---|---|---|---|---|---|---|---|
| | | 7 Å | | | | 7.5 Å | | | | 8 Å | | | |
| | | *g* | *a* | *d* | *e* | *g* | *a* | *d* | *e* | *g* | *a* | *d* | *e* |
| X-ray structures | apCC-Tet* | - | - | - | - | - | 12 | 12 | 10 | 2 | 12 | 12 | 12 |
| | apCC-Tet*[3] | - | - | - | - | - | 8 | 8 | 4 | - | 8 | 8 | 8 |
| | apCC-Tet*[3]-$A_2B_2$[a] | - | - | - | - | - | 8 | 7 | 10 | 1 | 8 | 9 | 11 |
| | sc-apCC-4 | - | - | - | - | - | 12 | 8 | 13 | - | 12 | 9 | 13 |
| AlphaFold2 predictions | apCC-Tet* | - | 12 | 12 | 4 | - | 12 | 12 | 12 | - | 12 | 12 | 12 |
| | apCC-Tet*[3] | - | 8 | 8 | 8 | - | 8 | 8 | 8 | - | 8 | 8 | 8 |
| | apCC-Tet*[3]-$A_2B_2$ | - | 8 | 8 | 8 | - | 8 | 8 | 8 | - | 8 | 8 | 8 |
| | sc-apCC-4 | - | 12 | 12 | 2 | - | 12 | 12 | 14 | 3 | 12 | 12 | 14 |

[a]The numbers of knobs correspond to the average per coiled-coil assembly as the crystal structure of apCC-Tet*[3]-$A_2B_2$ was composed of 2 coiled coils in the asymmetric unit.

**Table S6. Sequences comparison of historical *de novo* antiparallel 4-helix bundles.** Structurally resolved antiparallel 4-helix bundles with identified knobs-into-holes packing by SOCKET2[24] were visually inspected. These were mainly detected from the CC+ database.[1] Amino acids found at ***g***, ***a***, ***d*** and ***e*** positions are given below. Antiparallel 4-helix bundles without full structural characterization were not considered.

| PDB ID | Composition | | | | Reference |
|---|---|---|---|---|---|
| | ***g*** | ***a*** | ***d*** | ***e*** | |
| **8A3G, 8A3I, 8A3J, 8A3K** | Q | L | I | A | This study |
| **1EC5** | Q, L, D, W | L, E, Y | L, A, I, H | L, I | Lombardi, et al. (2000)[25] |
| **1MFT, 1U7J, 1U7M, 1Y47, 1JMB, 1JM0** | Y, L, W, D | L, E, Y | L, A, I, H | Y, M, S, L, I | Lahr, et al. (2005)[26]<br>Di Costanzo, et al. (2001)[27] |
| **1W5H, 2CCN, 2CCF** | K, E, L | L | I | E, L, C (or S), K | Yadav, et al. (2006)[28] |
| **1W5J, 1W5K** | R, K, E, L | L | I | E, L, C, R | Yadav, et al. (2006)[28] |
| **2B1F** | R, H, E, Q | L | A, L | V, N | Deng, et al. (2006)[29] |
| **2B22** | H, V | V, L | L, V | L, N | Deng, et al. (2006)[29] |
| **2NRN** | Q, E, H, R | L | V, K, E, N, L | V, N, K, E, Y, A | Deng, et al. (2007)[30] |
| **3S0R** | A | L | A | E, Q, R | Grigoryan, et al. (2011)[31] |
| **4HB1** | L, A | L, A | A, L | K, Q | Schafmeister, et al. (1997)[32] |
| **5J0K, 6EGC** | L, S | L, A, I, V | L, Q, V, I | E, K, R | Boyken, et al. (2016)[33]<br>Chen, et al. (2019)[34] |
| **5J10** | L, S, I | V, L, A, I, M | I, Q, L, M | A, K, E, L | Boyken, et al. (2016)[33] |
| **5TGY** | A, L, I, V, Q | F, I, L, G | L, I, A, W, F | E, R, A, V | Polizzi, et al. (2017)[35] |
| **5VJS, 5VJU** | L, Q, G, T, D | R, A, L, G, E, Y, V | S, H, F, L, A, I, G | Q, L, I, S | Unpublished |
| **6C52** | K | L, A | L, A | R, E, Q | Zhang, et al. (2018)[36] |
| **6DLC** | L, I, N, R, Q, V, A | L, S, V, F, I, A | F, H, I, A, S, V, L, Y | L, I, V, E, T, N, R | Chen, et al. (2019)[37] |
| **6DMP** | L, I, A | L, I, H, V, S | A, Q, L, M, S, I | Q, L, E, K, R | Chen, et al. (2019)[37] |
| **6Q5S, 6Q5R** | K, E | L | L | A | Rhys, et al (2019)[38] |
| **6W70** | E, V, A, Y, L, K, T, I | F, L, A, G, Y, M, T | V, A, M, L, H, I, F, G | E, A, K, V, L, T | Polizzi, et al. (2020)[39] |

#### 2.2. Supplementary figures

##### 2.2.1. Rational and computational design

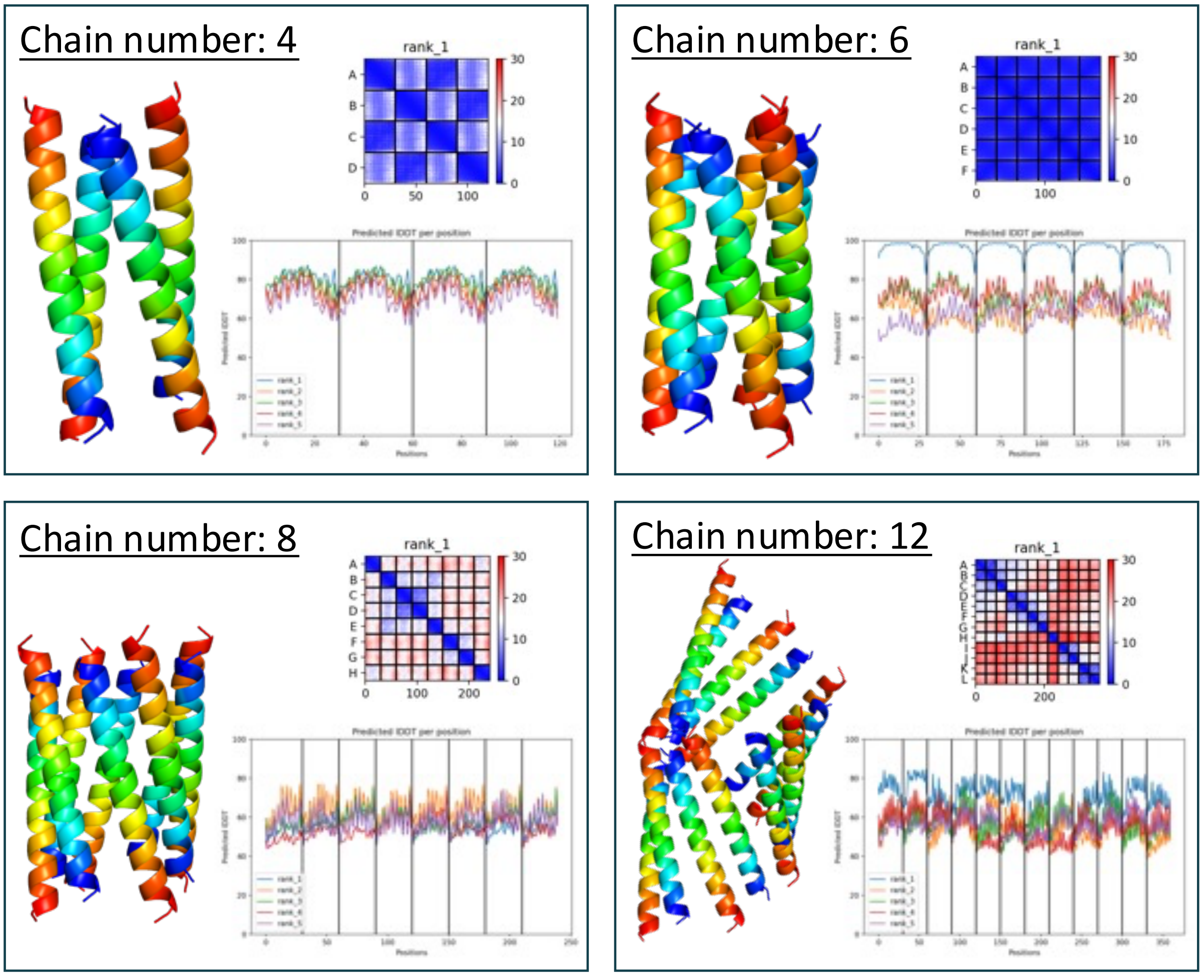

**Figure S1. AlphaFold2 predictions for pLLL.** Assembly predictions for the pLLL sequence with different numbers of chains as the input for AlphaFold2-multimer (ColabFold). Each panel shows the top-ranked prediction (left) with the associated Predicted Alignment Error (PAE) graph (top right) to assess the confidence about the interface, and the predicted IDDT graph (bottom right) to assess model confidence at each position. In this case, AlphaFold2-multimer predicted with high confidence a 6-helix bundle for the pLLL sequence with 6 peptide chains as input.

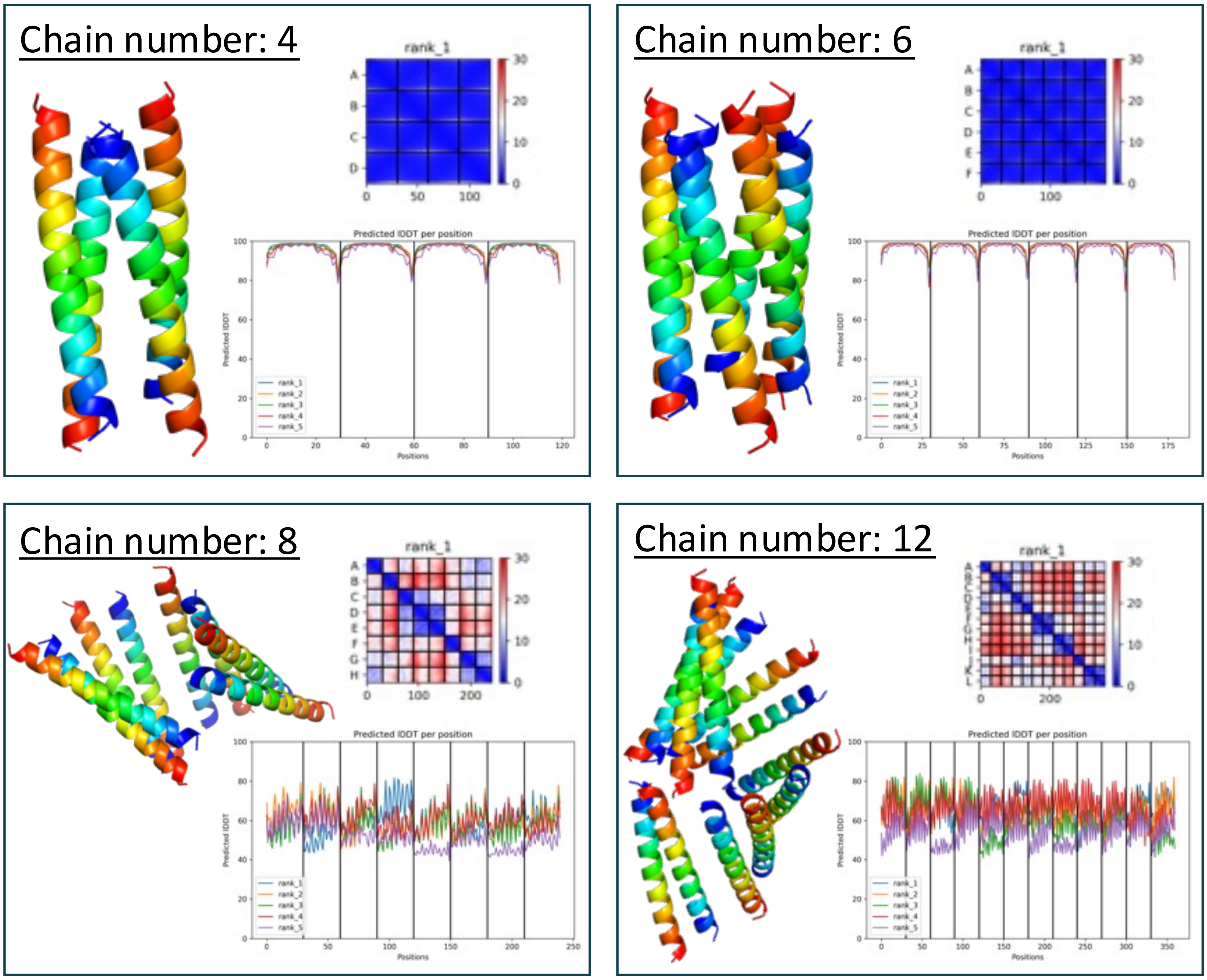

**Figure S2. AlphaFold2 predictions for pLLI**. Assembly predictions for the pLLI sequence with different numbers of chains as the input for AlphaFold2-multimer (ColabFold). Each panel shows the top-ranked prediction (left) with the associated Predicted Alignment Error (PAE) graph (top right) to assess the confidence about the interface, and the predicted lDDT graph (bottom right) to assess model confidence at each position. In this case, AlphaFold2 predicted with high confidence 4- or 6-helix bundle for pLLI sequence when the requested oligomeric state was 4 or 6, respectively.

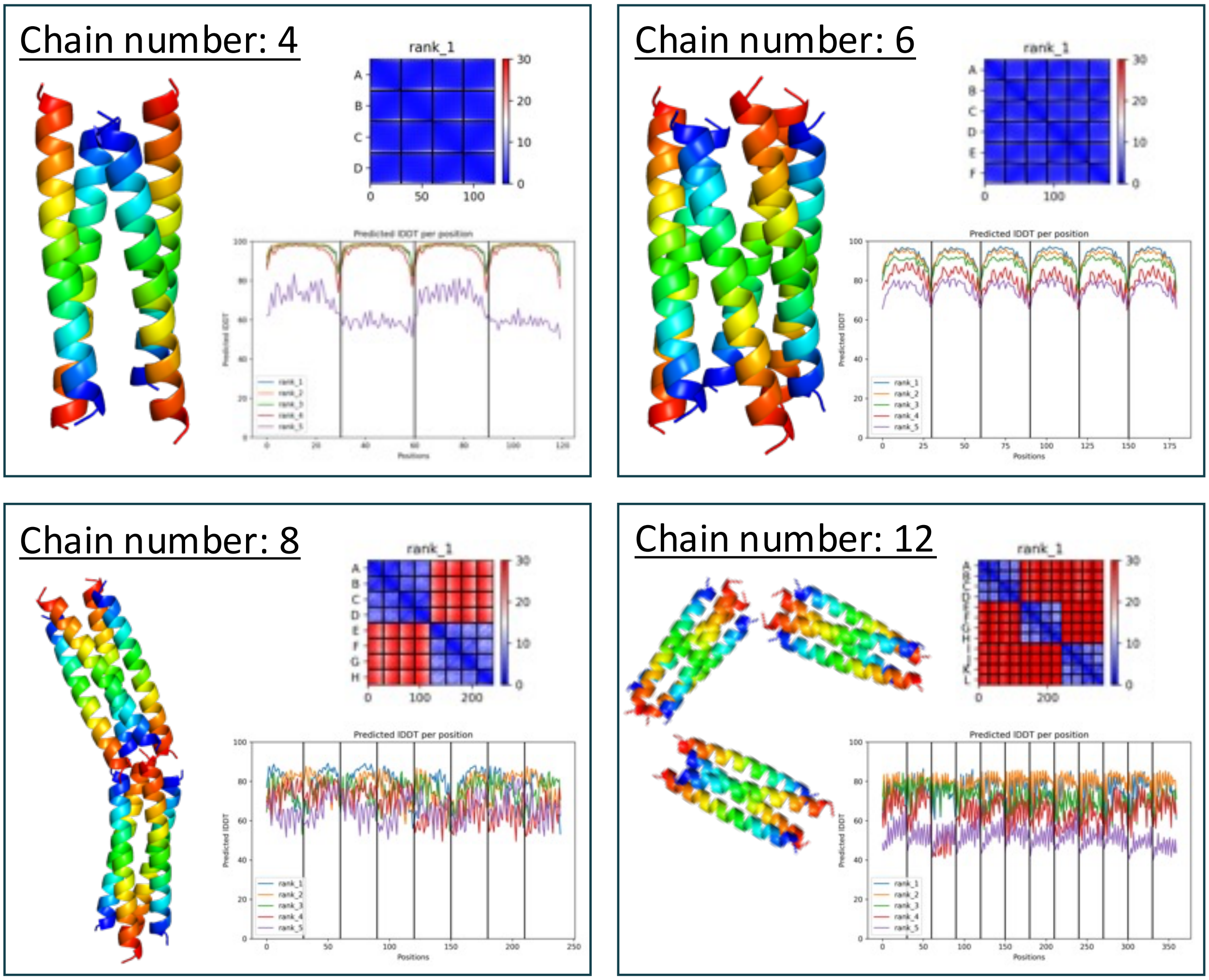

**Figure S3. AlphaFold2 predictions of pQLI (apCC-Tet*).** Assembly predictions for the pQLI sequence with different numbers of chains as the input for AlphaFold2-multimer (ColabFold). Each panel shows the top-ranked prediction (left) with the associated Predicted Alignment Error (PAE) graph (top right) to assess the confidence about the interface, and the predicted lDDT graph (bottom right) to assess model confidence at each position. In this case, AlphaFold2 slightly predicted better and with high confidence a 4-helix bundle for the pQLI sequence with 4 peptide chains as input. Interestingly, when the requested oligomeric was higher (i.e., 8 or 12), AlphaFold2 predicted the formation of several tetramers with plDDT around 80%.

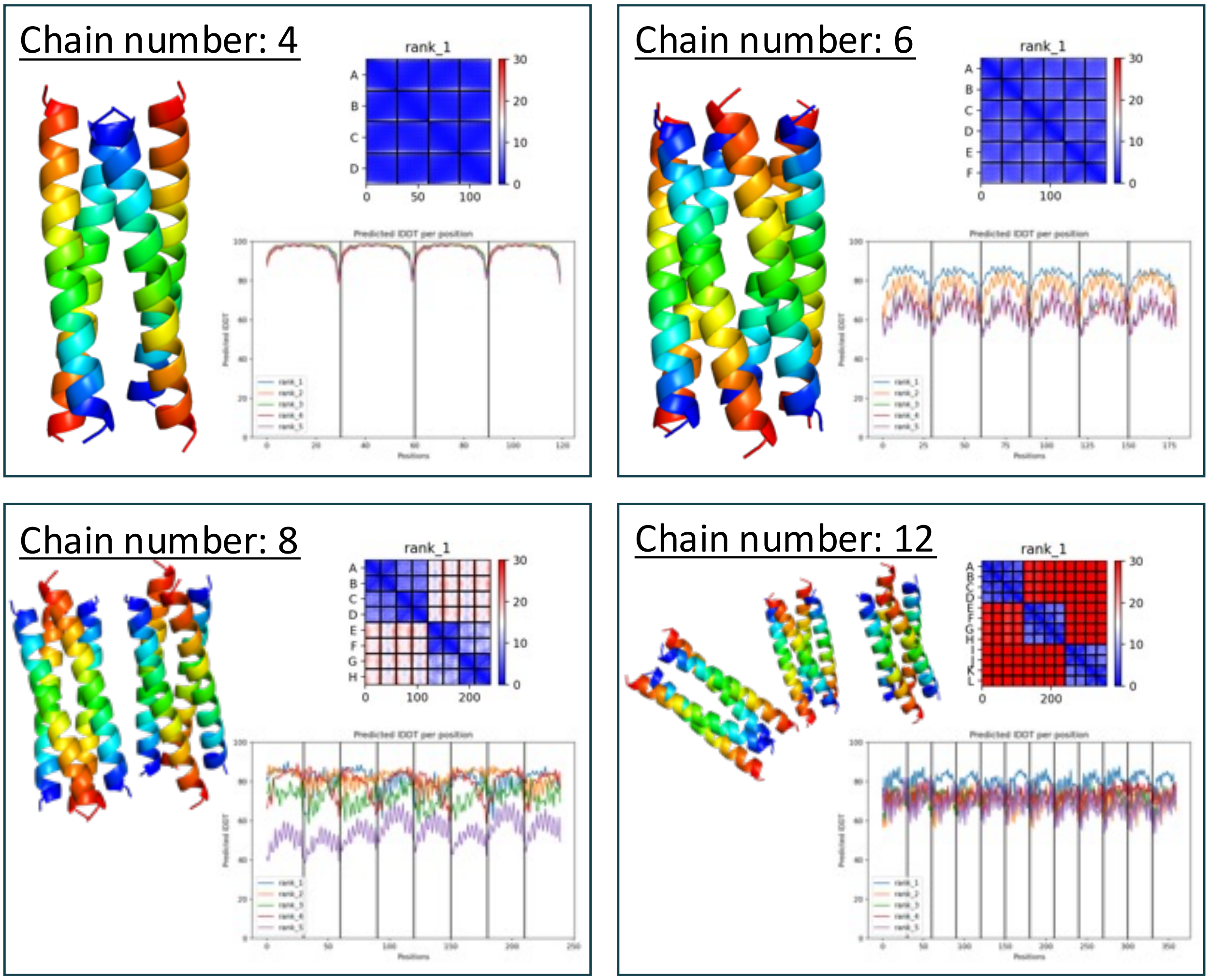

**Figure S4. AlphaFold2 predictions of pQLL** Assembly predictions for the pQLL sequence with different numbers of chains as the input for AlphaFold2-multimer (ColabFold). Each panel shows the top-ranked prediction (left) with the associated Predicted Alignment Error (PAE) graph (top right) to assess the confidence about the interface, and the predicted lDDT graph (bottom right) to assess model confidence at each position. In this case, AlphaFold2 predicted with high confidence a 4-helix bundle for pQLL sequence when the requested oligomeric state was 4. The hexameric assembly was predicted with lower confidence for oligomeric state of 6 as input. Interestingly, when the requested oligomeric was higher (i.e., 8 or 12), AlphaFold2 predicted the formation of several tetramers with plDDT around 80%.

#### 2.2.2. Characterization of homopeptides

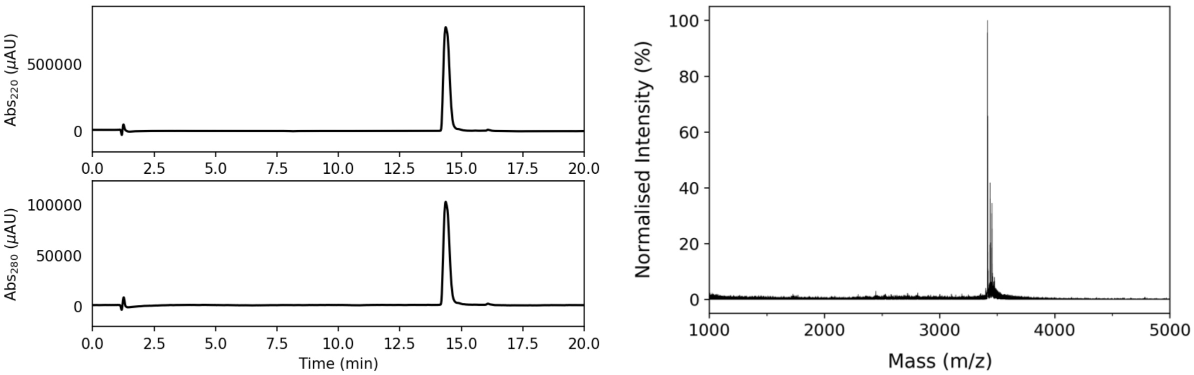

**Figure S5. Characterization of pLLL. Left:** Analytical HPLC chromatogram at 220 nm (top) and 280 nm (bottom) from a gradient of 40 to 100% MeCN (0.1% TFA) in $H_2O$ (0.1% TFA) with column oven at 50 °C. **Right:** MALDI-TOF MS. Calculated mass: 3416.2 Da $[M+H]^+$. Observed mass: 3417.1 Da $[M+H]^+$.

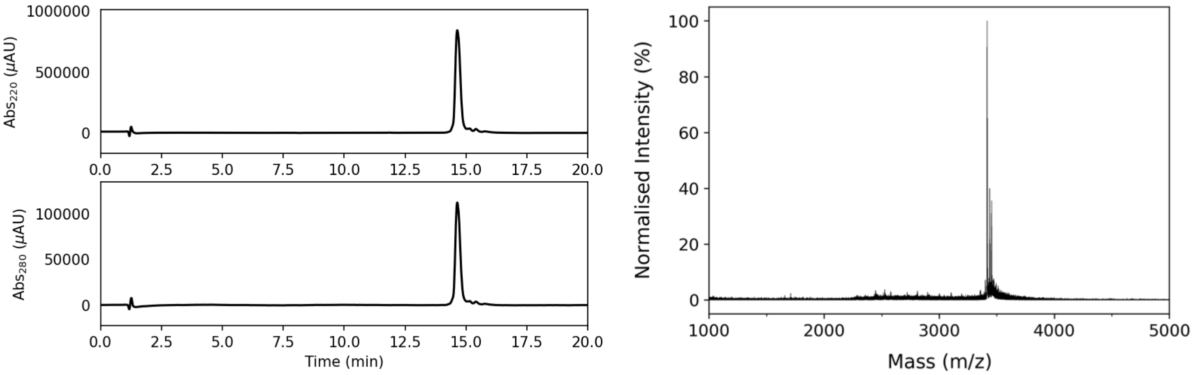

**Figure S6. Characterization of pLLI. Left:** Analytical HPLC chromatogram at 220 nm (top) and 280 nm (bottom) from a gradient of 40 to 100% MeCN (0.1% TFA) in $H_2O$ (0.1% TFA) with column oven at 50 °C. **Right:** MALDI-TOF MS. Calculated mass: 3416.2 Da $[M+H]^+$. Observed mass: 3417.2 Da $[M+H]^+$.

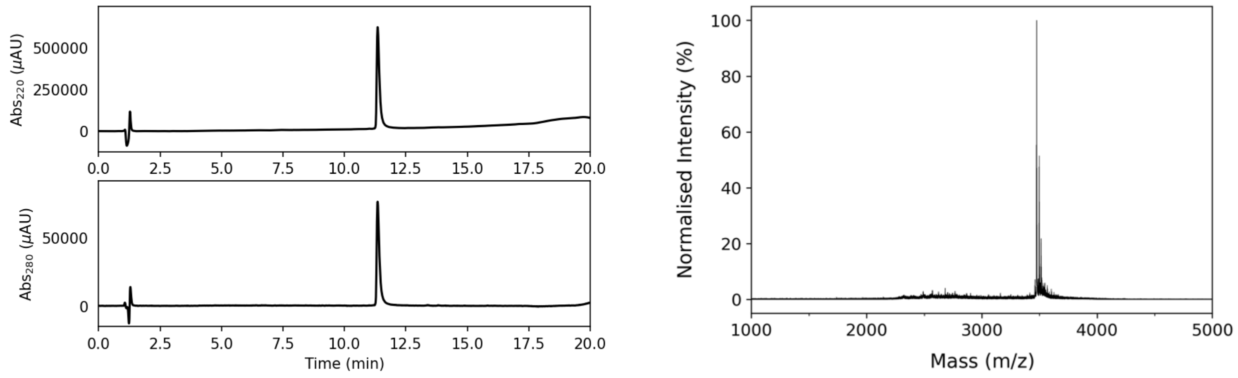

**Figure S7. Characterization of pQLI (apCC-Tet*). Left:** Analytical HPLC chromatogram at 220 nm (top) and 280 nm (bottom) from a gradient of 20 to 80% MeCN (0.1% TFA) in $H_2O$ (0.1% TFA). **Right:** MALDI-TOF MS. Calculated mass: 3476.1 Da $[M+H]^+$. Observed mass: 3476.0 Da $[M+H]^+$.

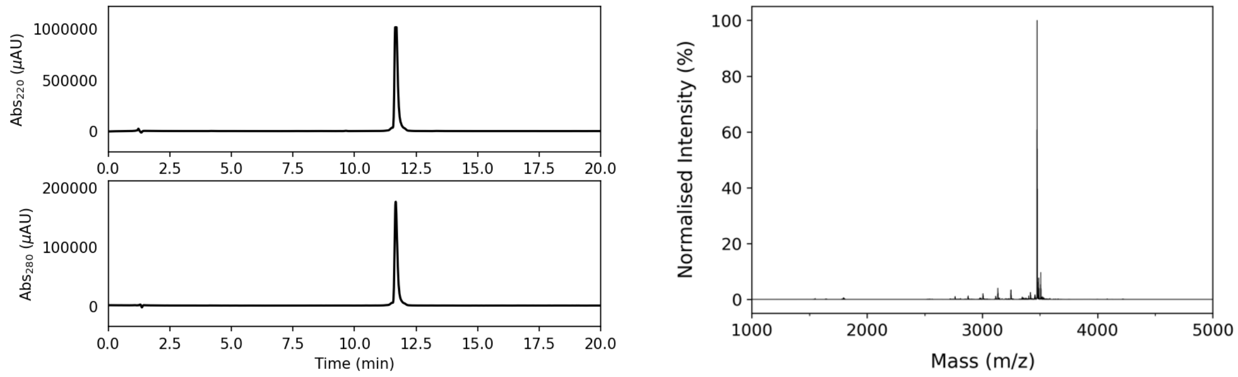

**Figure S8. Characterization of pQLL. Left:** Analytical HPLC chromatogram at 220 nm (top) and 280 nm (bottom) from a gradient of 20 to 80% MeCN (0.1% TFA) in $H_2O$ (0.1% TFA). **Right:** Deconvoluted ESI MS. Calculated mass: 3476.1 Da $[M+H]^+$. Observed mass: 3476.6 Da $[M+H]^+$.

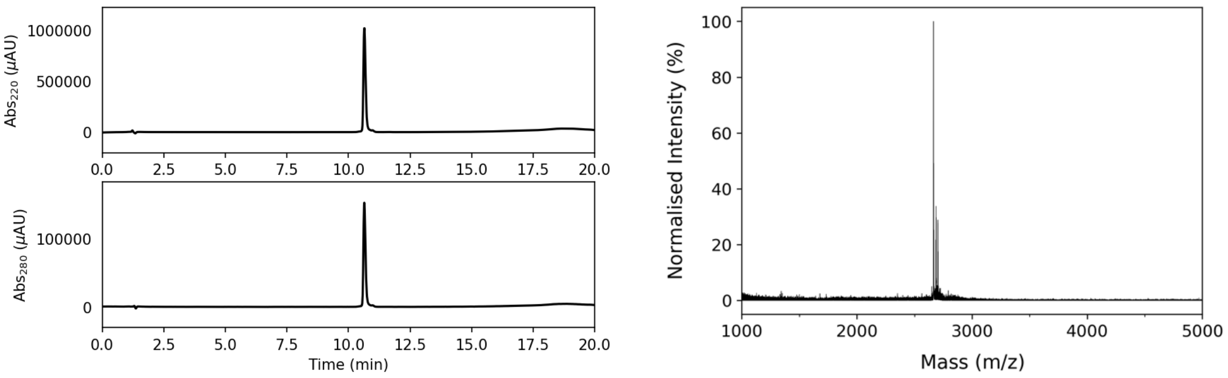

**Figure S9. Characterization of pQLI$^3$ (apCC-Tet*$^3$). Left:** Analytical HPLC chromatogram at 220 nm (top) and 280 nm (bottom) from a gradient of 20 to 80% MeCN (0.1% TFA) in $H_2O$ (0.1% TFA). **Right:** MALDI-TOF MS. Calculated mass: 2664.1 Da $[M+H]^+$. Observed mass: 2664.4 Da $[M+H]^+$.

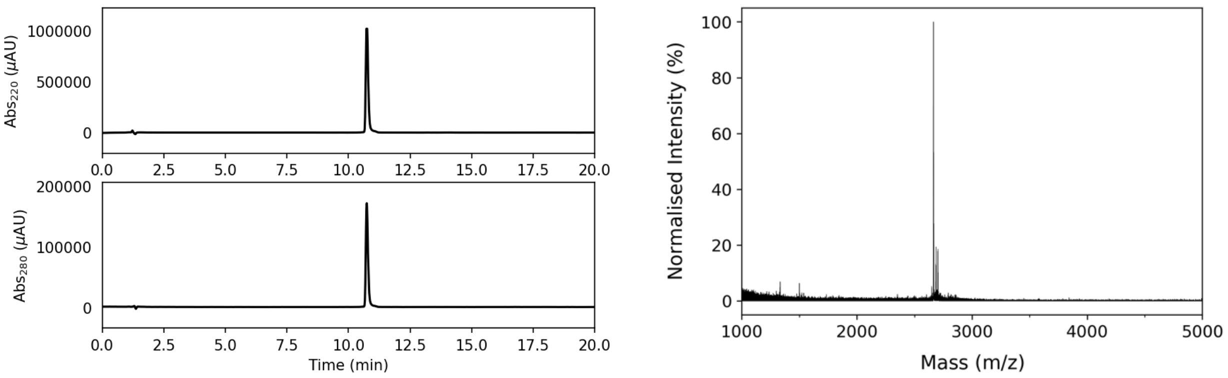

**Figure S10. Characterization of pQLL$^3$. Left:** Analytical HPLC chromatogram at 220 nm (top) and 280 nm (bottom) from a gradient of 20 to 80% MeCN (0.1% TFA) in $H_2O$ (0.1% TFA). **Right:** MALDI-TOF MS. Calculated mass: 2664.1 Da $[M+H]^+$. Observed mass: 2664.4 Da $[M+H]^+$.

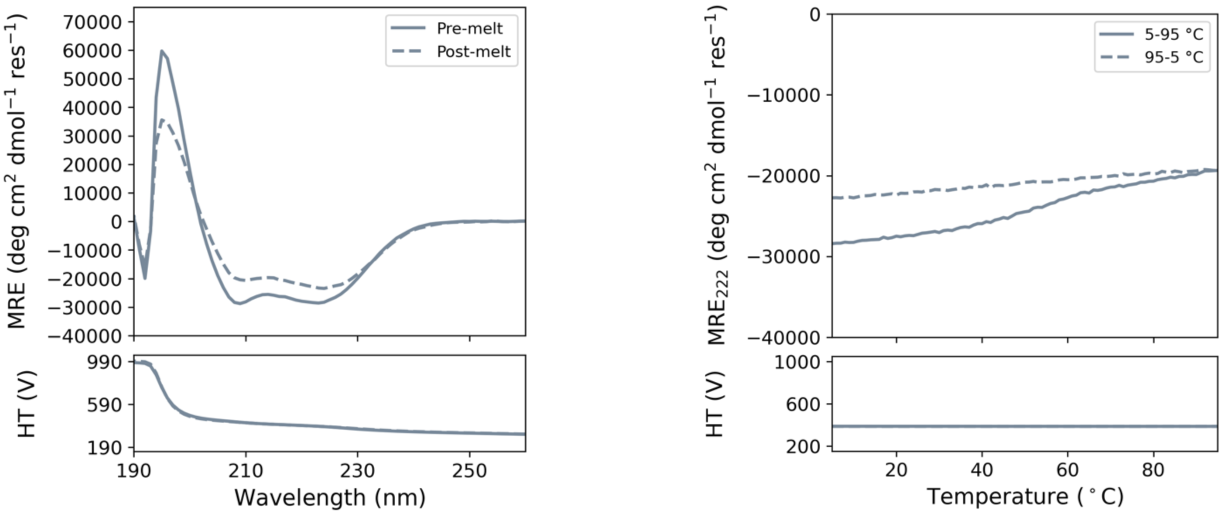

**Figure S11. CD spectroscopy of pLLL. Left:** CD spectra at 5 °C before thermal ramp up (solid line) and after thermal ramp down (dashed line). **Right:** Thermal profiles (ramping up and ramping down) monitored at 222 nm. Bottom panels show corresponding high-tension (HT) voltages applied to the detector during measurements. Conditions: 50 μM peptide concentration, PBS, pH 7.4.

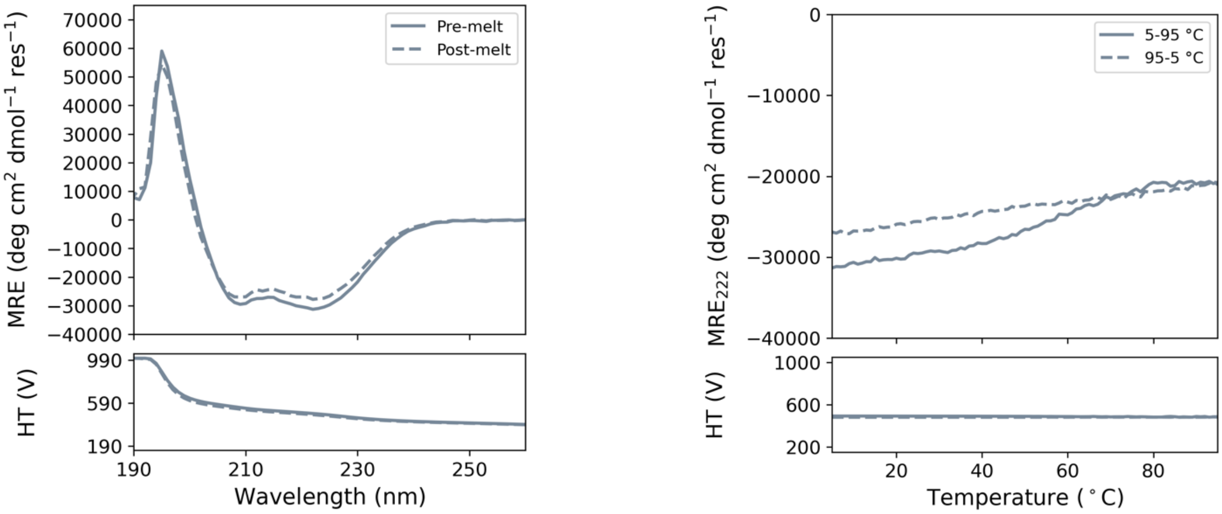

**Figure S12. CD spectroscopy of pLLI. Left:** CD spectra at 5 °C before thermal ramps up (solid line) and after thermal ramps down (dashed line). **Right:** Thermal profiles (ramping up and ramping down) monitored at 222 nm. Bottom panels show corresponding high-tension (HT) voltages applied to the detector during measurements. Conditions: 50 μM peptide concentration, PBS, pH 7.4.

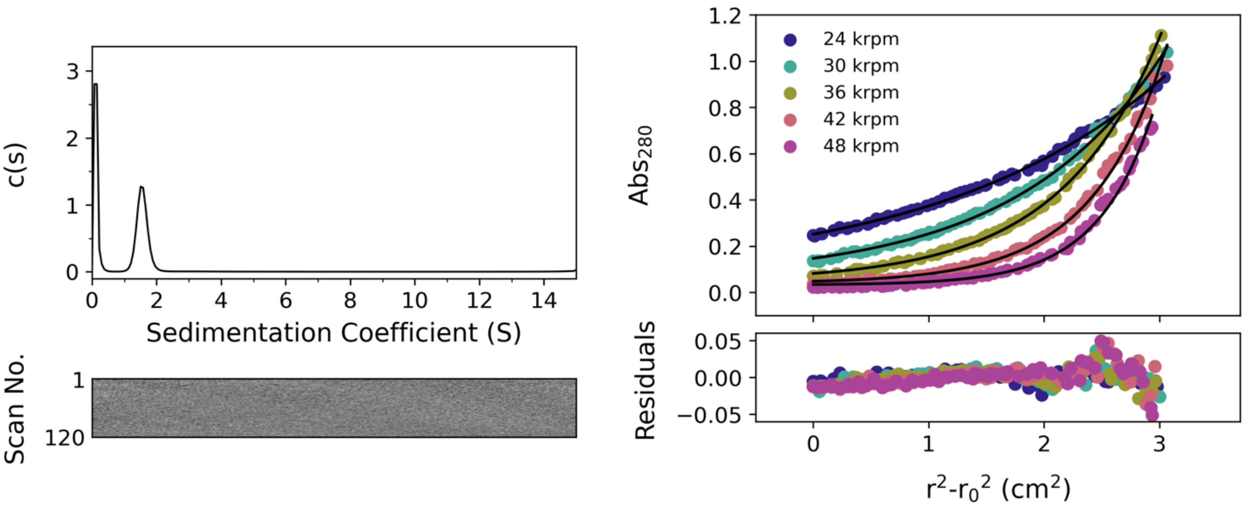

**Figure S13. AUC data for pLLL. Left:** Sedimentation velocity (SV) AUC data for pLLL peptide at 50 krpm ($\bar{v}$ = 0.799 cm$^3$ g$^{-1}$). Residuals are shown as a bitmap below the fitted data. Continuous c(s) distribution returned a molecular mass of 20583 Da corresponding to 6.0 x monomer mass at 95% confidence level (f/f0 = 1.205, s = 1.556 S, $s_{20,w}$ = 1.627S). **Right:** Sedimentation equilibrium (SE) AUC data for pLLL between 24 and 48 krpm at 6 krpm intervals. Fitted single-ideal species model curves are overlaid in black and gave a molecular mass 20049 Da corresponding to 5.9 x monomer mass, 95% confidence limits 19945-20268 Da. Conditions: 150 $\mu$M peptide, PBS, pH 7.4, 20 °C.

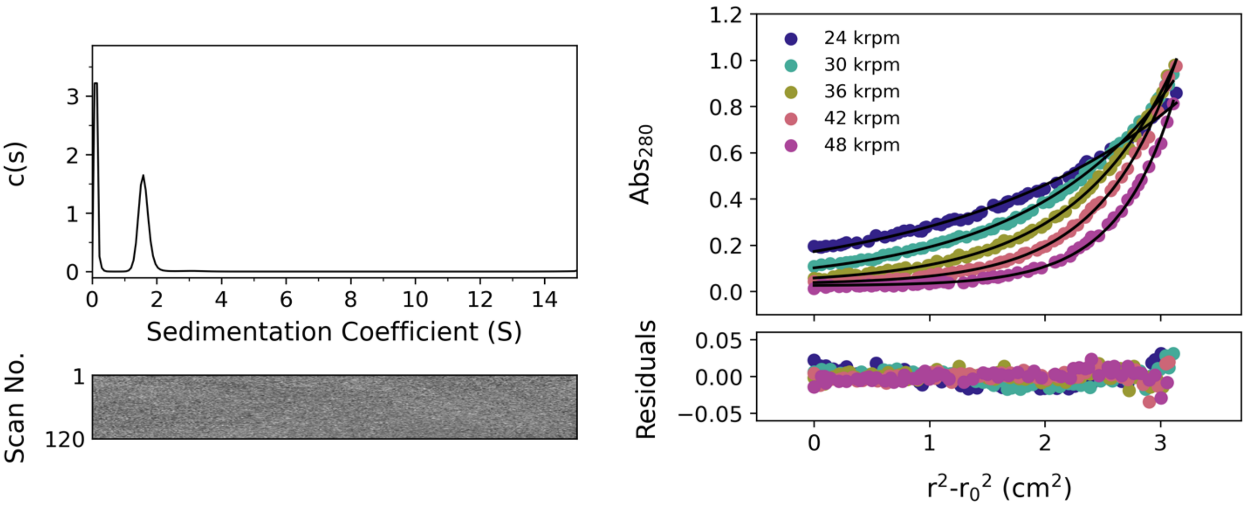

**Figure S14. AUC data for pLLI. Left:** Sedimentation velocity (SV) AUC data for pLLI peptide at 50 krpm ($\bar{v}$ = 799 cm$^3$ g$^{-1}$). Residuals are shown as a bitmap below the fitted data. Continuous c(s) distribution returned a molecular mass of 21537 Da corresponding to 6.3 x monomer mass at 95% confidence level (f/f0 = 1.212, s = 1.595 S, $s_{20,w}$ = 1.668 S). **Right:** Sedimentation equilibrium (SE) AUC data for pLLI between 24 and 48 krpm at 6 krpm intervals. Fitted single-ideal species model curves are overlaid in black and gave a molecular mass 20079 Da corresponding to 5.9 x monomer mass, 95% confidence limits 19889-20287 Da. Conditions: 150 $\mu$M peptide, PBS, pH 7.4, 20 °C.

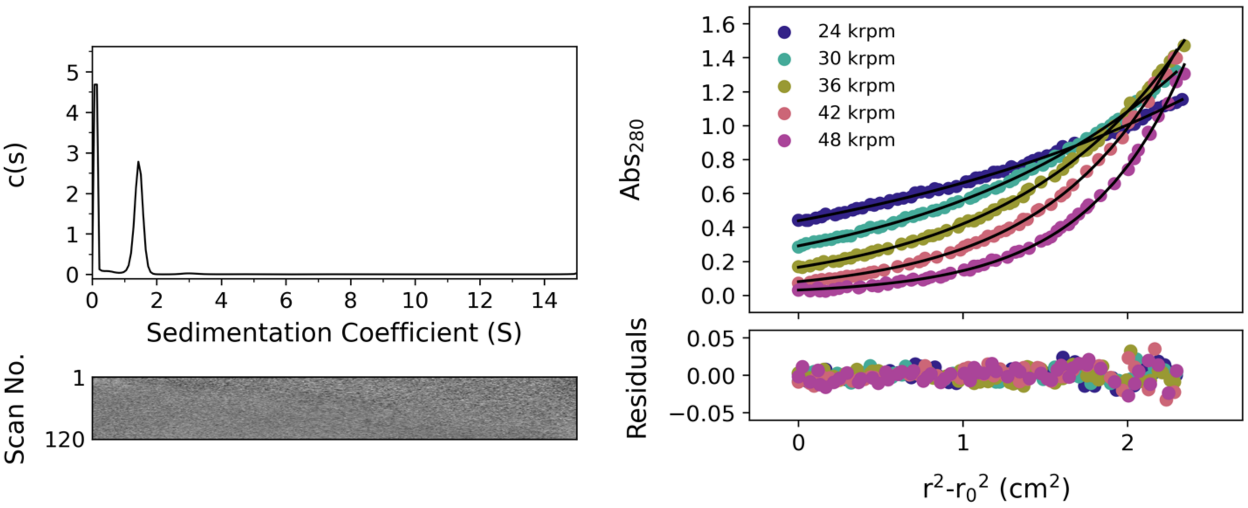

**Figure S15. AUC data for pQLI (apCC-Tet*). Left:** Sedimentation velocity (SV) AUC data for apCC-Tet* at 50 krpm ($\bar{v}$ = 0.766 $cm^3$ $g^{-1}$). Residuals are shown as a bitmap below the fitted data. Continuous c(s) distribution returned a molecular mass of 15800 Da corresponding to 4.5 x monomer mass at 95% confidence level (f/f0 = 1.298, s = 1.434 S, $s_{20,w}$ = 1.492 S). **Right:** Sedimentation equilibrium (SE) AUC data for apCC-Tet* between 24 and 48 krpm at 6 krpm intervals. Fitted single-ideal species model curves are overlaid in black and gave a molecular mass 14134 Da corresponding to 4.1 x monomer mass, 95% confidence limits 14053-14306 Da. Conditions: 150 μM and 75 μM peptide for SV and SE experiments respectively, PBS, pH 7.4, 20 °C.

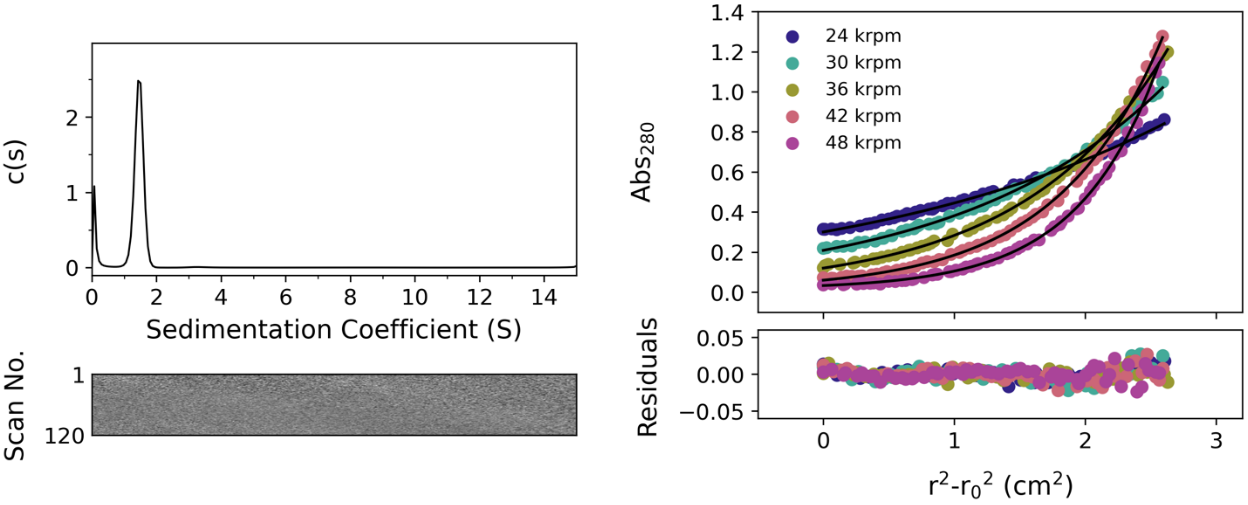

**Figure S16. AUC data for pQLL Left:** Sedimentation velocity (SV) AUC data for pQLL peptide at 50 krpm ($\bar{v}$ = 0.766 $cm^3$ $g^{-1}$). Residuals are shown as a bitmap below the fitted data. Continuous c(s) distribution returned a molecular mass of 15064 Da corresponding to 4.3 x monomer mass at 95% confidence level (f/f0 = 1.238, s = 1.457 S, $s_{20,w}$ = 1.516 S). **Right:** Sedimentation equilibrium (SE) AUC data for pQLL between 24 and 48 krpm at 6 krpm intervals. Fitted single-ideal species model curves are overlaid in black and gave a molecular mass 13698 Da corresponding to 3.9 x monomer mass, 95% confidence limits 13454-13740 Da. Conditions: 150 μM and 75 μM peptide for SV and SE experiments respectively, PBS, pH 7.4, 20 °C.

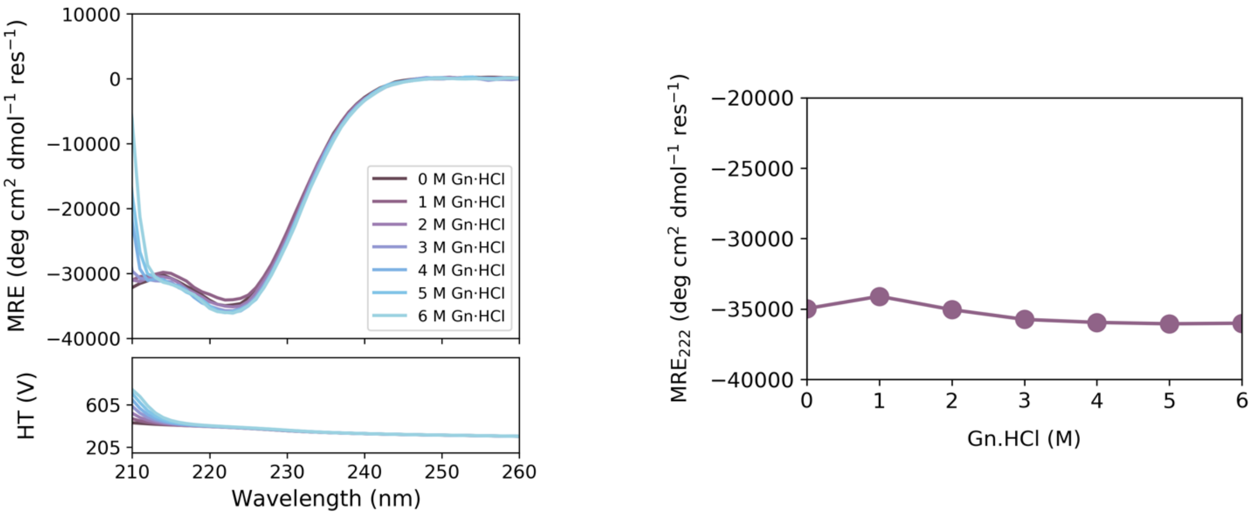

**Figure S17. Chemical denaturation of pQLI (apCC-Tet*). Left:** CD full scan from 210 to 260 nm at 5 °C of apCC-Tet* incubated with increasing concentration of guanidinium hydrochloride (Gn·HCl). **Right:** Chemical denaturation curve measured at 222 nm for apCC-Tet* (deduced from left spectra). Conditions: 50 µM peptide concentration, ± Gn·HCl, PBS, pH 7.4 and incubation overnight before measurements.

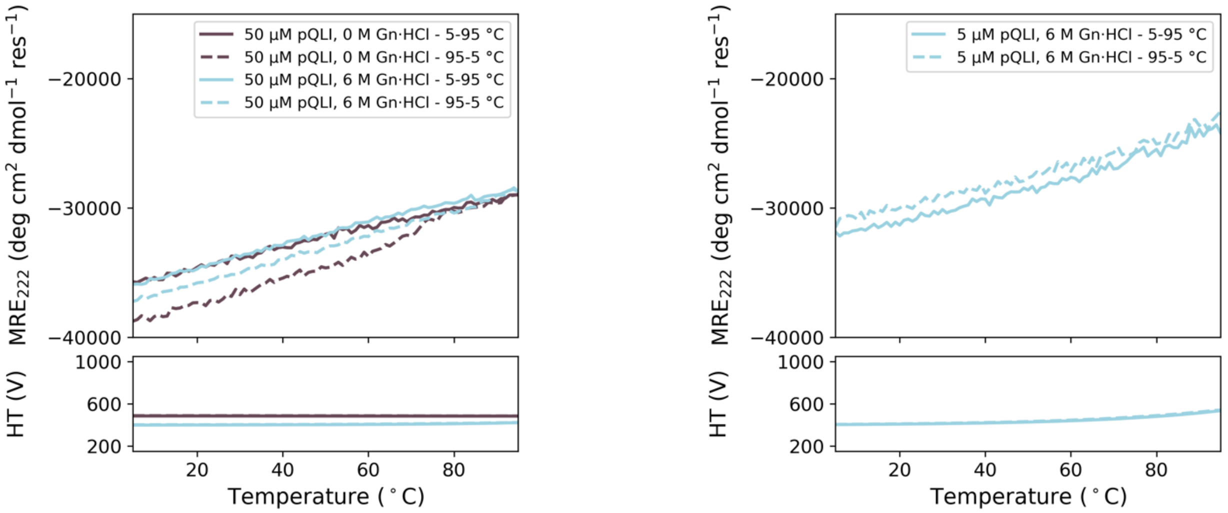

**Figure S18. Temperature profile of pQLI (apCC-Tet*) in presence of high concentration of denaturant. Left:** Thermal profiles (ramping up and ramping down) monitored at 222 nm for 50 µM of apCC-Tet* with or without 6 M Gn·HCl. **Right:** Thermal profiles (ramping up and ramping down) monitored at 222 nm in presence of 6 M Gn·HCl for lower concentration of apCC-Tet* (5 µM). Peptide samples were incubated overnight with guanidinium before measurements.

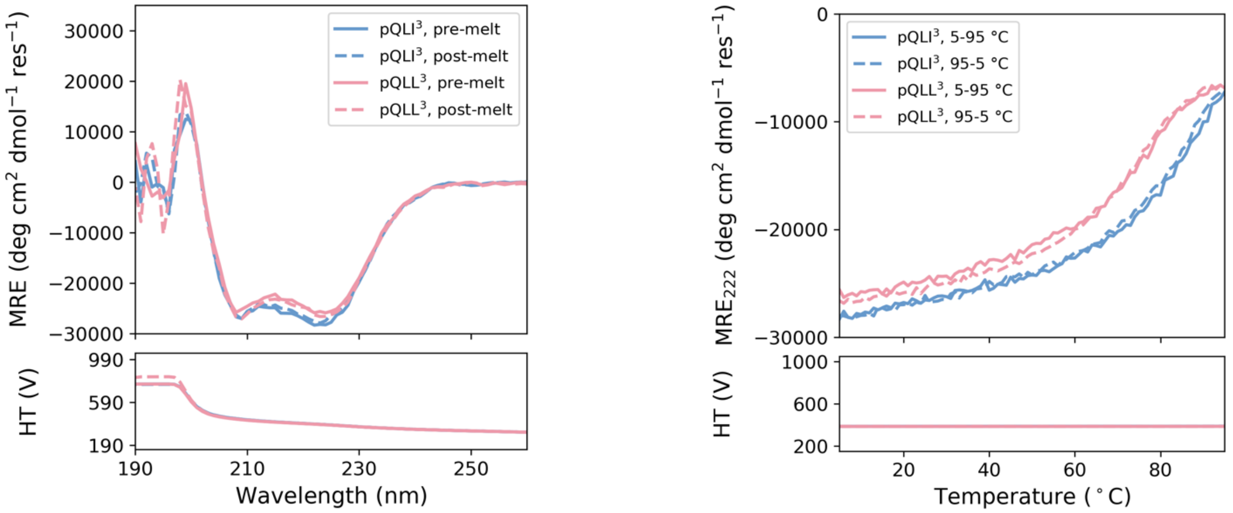

**Figure S19. CD spectroscopy of 3-heptad homotetramers $pQLI^3$ ($apCCTet^{*3}$) and $pQLL^3$ at low concentration. Left:** CD spectra at 5 °C before thermal ramp up (solid line) and after thermal ramp down (dashed line) for $pQLI^3$ (pink) and $pQLL^3$ (blue). **Right:** Thermal profiles (ramping up and ramping down) monitored at 222 nm. Bottom panels show corresponding high-tension (HT) voltages applied to the detector during measurements. Conditions: 5 µM peptide concentration, PBS, pH 7.4.

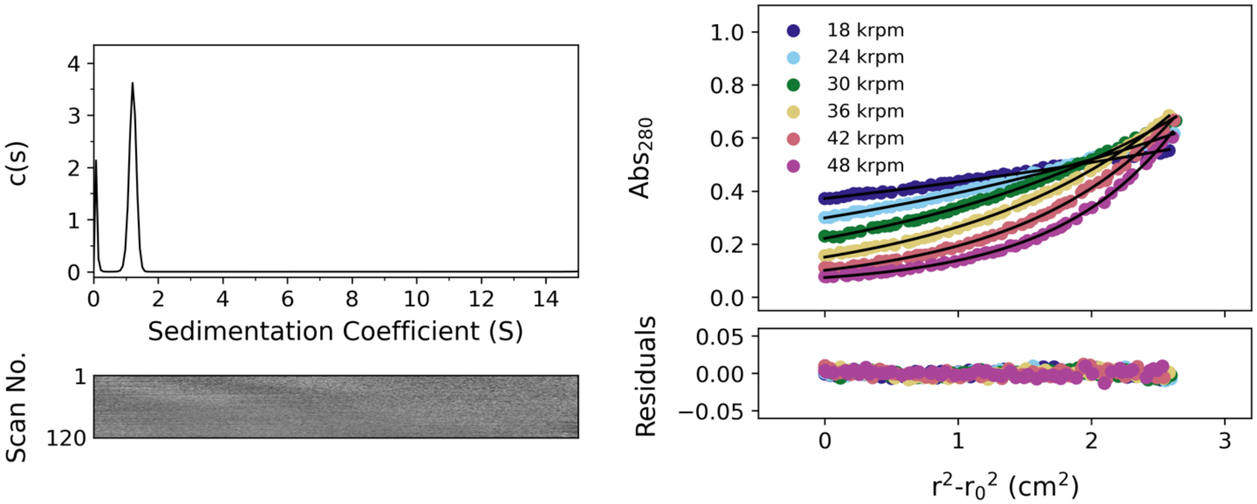

**Figure S20. AUC data for $pQLI^3$ ($apCC\text{-}Tet^{*3}$). Left:** Sedimentation velocity (SV) AUC data for $apCC\text{-}Tet^{*3}$ at 50 krpm ($\bar{v}$ = 0.761 $cm^3$ $g^{-1}$). Residuals are shown as a bitmap below the fitted data. Continuous c(s) distribution returned a molecular mass of 11431 Da corresponding to 4.3 x monomer mass at 95% confidence level (f/f0 = 1.273, s = 1.211 S, $s_{20,w}$ = 1.259 S). **Right:** Sedimentation equilibrium (SE) AUC data for $apCC\text{-}Tet^{*3}$ between 18 and 48 krpm at 6 krpm intervals. Fitted single-ideal species model curves are overlaid in black and gave a molecular mass 9217 Da corresponding to 3.5 x monomer mass, 95% confidence limits 9052-9298 Da. Conditions: 150 µM and 75 µM peptide for SV and SE experiments respectively, PBS, pH 7.4, 20 °C.

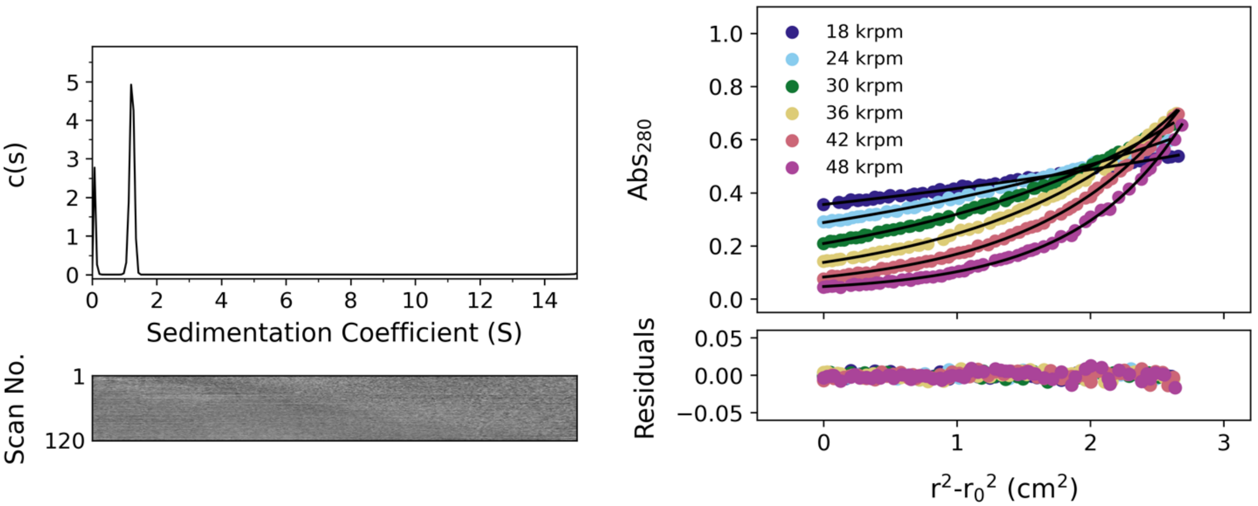

**Figure S21. AUC data for pQLL$^3$. Left:** Sedimentation velocity (SV) AUC data for pQLL$^3$ at 50 krpm ($\bar{v}$ = 0.761 cm$^3$ g$^{-1}$). Residuals are shown as a bitmap below the fitted data. Continuous c(s) distribution returned a molecular mass of 10758 Da corresponding to 4.0 x monomer mass at 95% confidence level (f/f0 = 1.205, s = 1.229 S, $s_{20,w}$ = 1.277 S). **Right:** Sedimentation equilibrium (SE) AUC data for pQLL$^3$ between 18 and 48 krpm at 6 krpm intervals. Fitted single-ideal species model curves are overlaid in black and gave a molecular mass 10152 Da corresponding to 3.8 x monomer mass, 95% confidence limits 10039-10227 Da. Conditions: 150 μM and 75 μM peptide for SV and SE experiments respectively, PBS, pH 7.4, 20 °C.

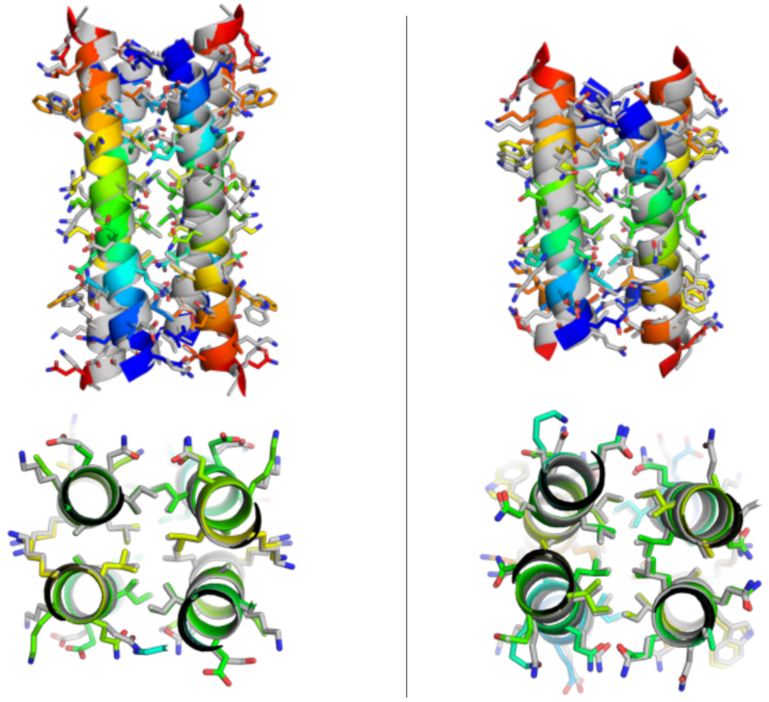

**Figure S22. Structural overlay of crystal structures and related AlphaFold2 predictions for pQLI (apCC-Tet*) analogues. Left:** Overlay of the crystal structure of pQLI (apCC-Tet*, chainbow coloring) and the corresponding AlphaFold2 prediction (grey) with a

RMSD$_{all-atom}$ = 0.359 Å and C$\alpha$ RMSD = 0.337 Å. **Right:** Overlay of the crystal structure of pQLI[3] (apCC-Tet*[3], chainbow coloring) and the corresponding AlphaFold2 prediction (grey) with a RMSD$_{all-atom}$ = 0.584 Å and C$\alpha$ RMSD = 0.590 Å.

##### 2.2.3. Characterization of heterotetramers

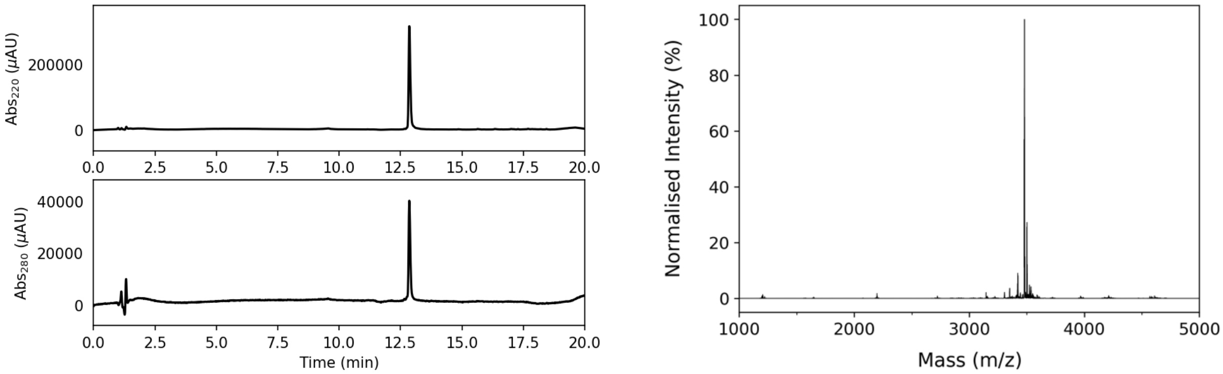

**Figure S23. Characterization of apCC-Tet*-A. Left:** Analytical HPLC chromatogram at 220 nm (top) and 280 nm (bottom) from a gradient of 20 to 80% MeCN in 25 mM $NH_4HCO_3$ in $H_2O$. **Right:** Deconvoluted ESI MS, negative mode. Calculated mass: 3478.9 Da [M]. Observed mass: 3479.8 Da [M].

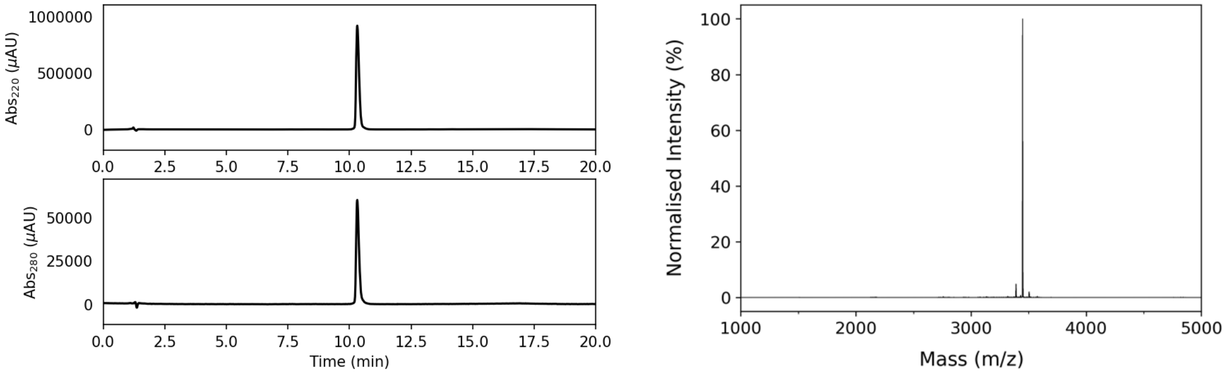

**Figure S24. Characterization of apCC-Tet*-B. Left:** Analytical HPLC chromatogram at 220 nm (top) and 280 nm (bottom) from a gradient of 20 to 80% MeCN (0.1% TFA) in $H_2O$ (0.1% TFA). **Right:** Deconvoluted ESI MS. Calculated mass: 3449.3 Da $[M+H]^+$. Observed mass: 3448.8 Da $[M+H]^+$.

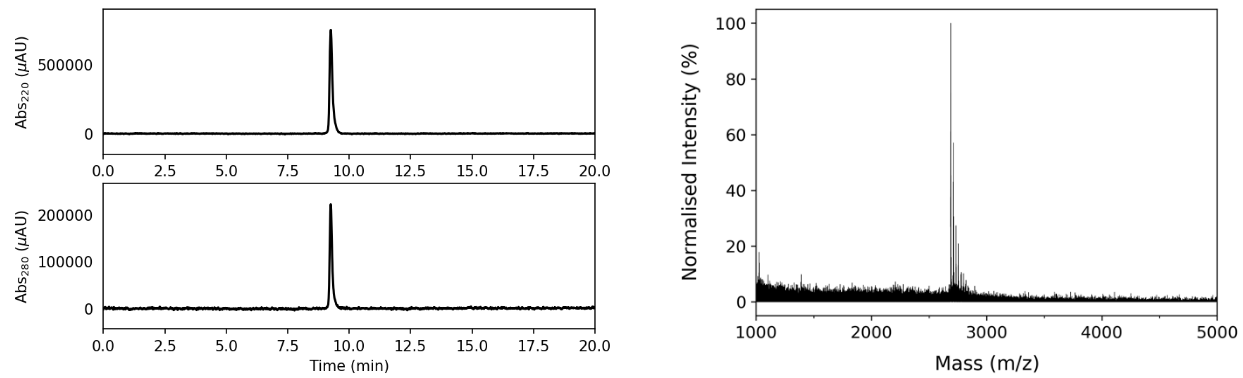

**Figure S25. Characterization of apCC-Tet*[3]-A. Left:** Analytical HPLC chromatogram at 220 nm (top) and 280 nm (bottom) from a gradient of 20 to 80% MeCN in 25 mM $NH_4HCO_3$ in $H_2O$. **Right:** MALDI-TOF MS. Calculated mass: 2668.0 Da $[M+H]^+$, 2690.0 Da $[M+Na]^+$. Observed mass: 2689.9 Da $[M+Na]^+$.

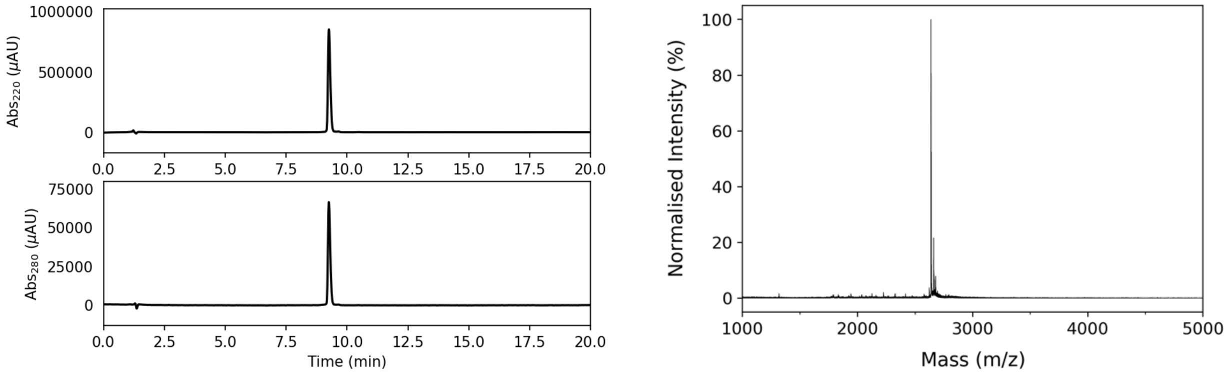

**Figure S26. Characterization of apCC-Tet*[3]-B. Left:** Analytical HPLC chromatogram at 220 nm (top) and 280 nm (bottom) from a gradient of 20 to 80% MeCN (0.1% TFA) in $H_2O$ (0.1% TFA). **Right:** MALDI-TOF MS. Calculated mass: 2639.3 Da $[M+H]^+$. Observed mass: 2639.4 Da $[M+H]^+$.

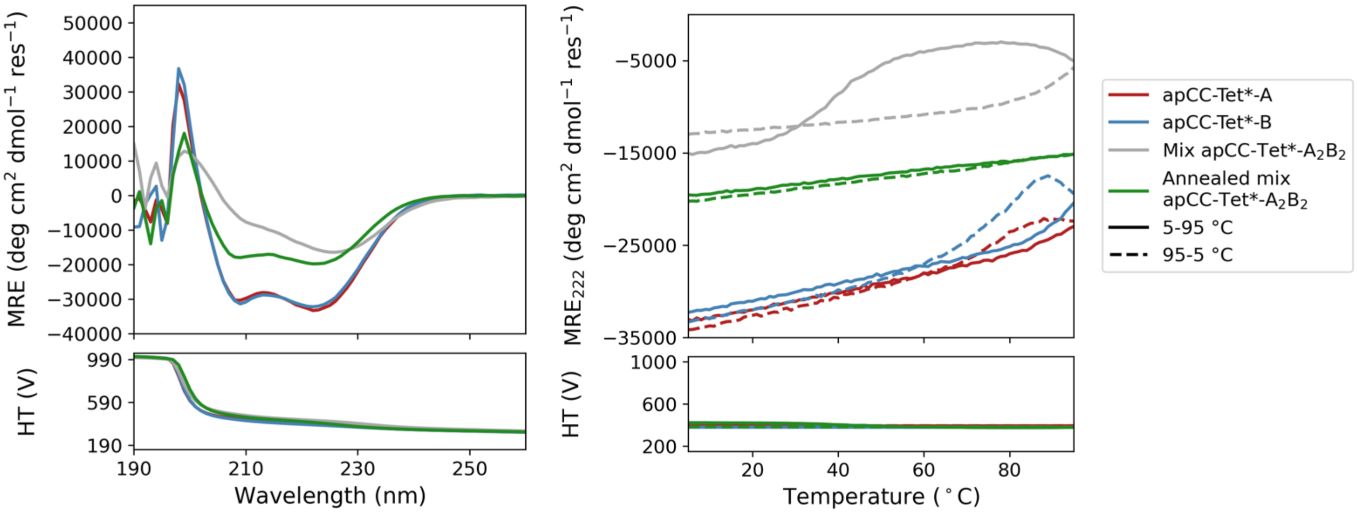

**Figure S27. CD spectroscopy of isolated peptides and mixture for apCC-Tet*-$A_2B_2$. Left:** CD spectra at 5 °C before thermal ramp up. **Right:** Thermal profiles (ramping up and ramping down) monitored at 222 nm. Bottom panels show corresponding high-tension (HT) voltages applied to the detector during measurements. Conditions: 10 μM peptide concentration, PBS (pH 6.5).

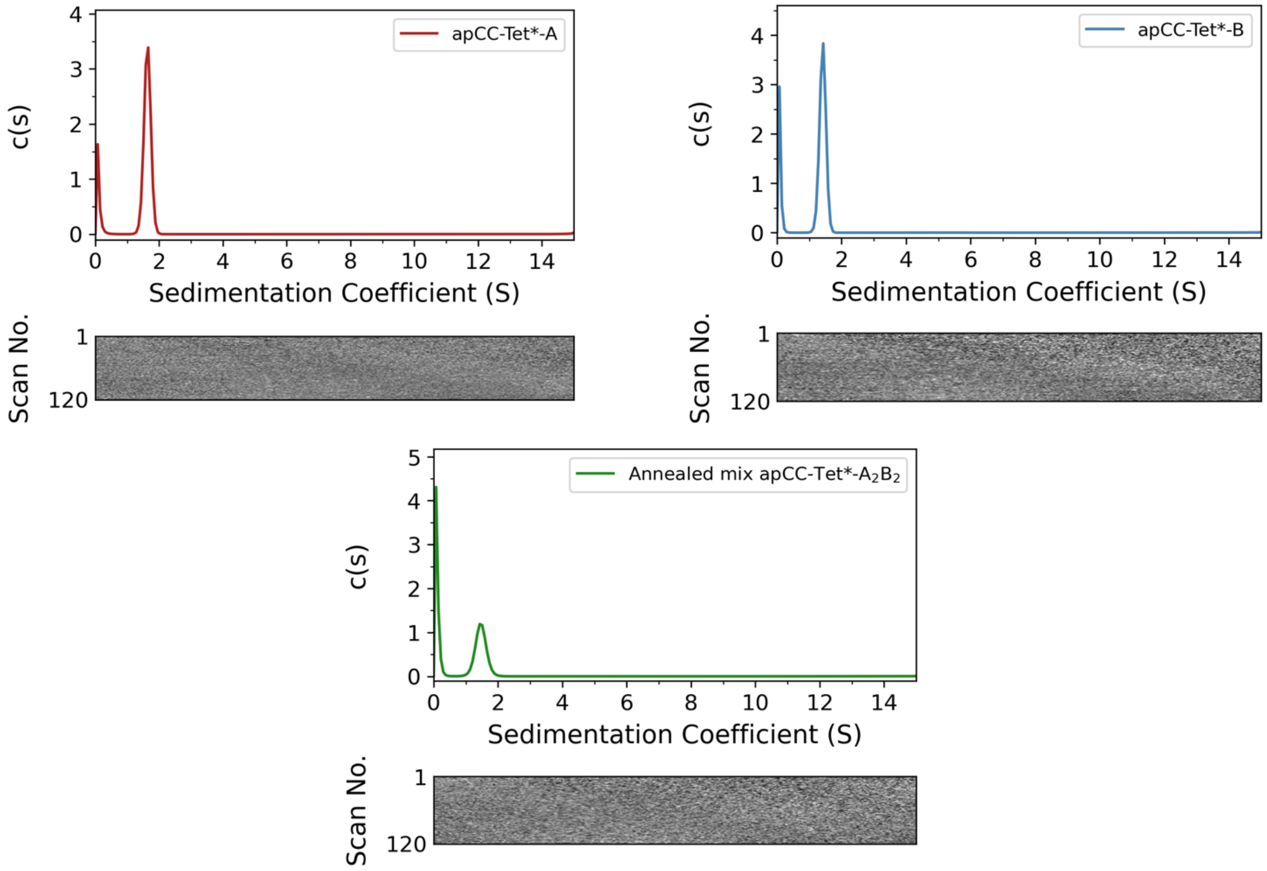

**Figure S28. AUC data isolated peptides and mixture for apCC-Tet*-$A_2B_2$. Top left:** Sedimentation velocity (SV) AUC data for apCC-Tet*-A at 50 krpm ($\overline{v}$ = 0.742 $cm^3$ $g^{-1}$). Continuous c(s) distribution returned a molecular mass of 15128 Da corresponding to 4.3 x monomer mass at 95% confidence level (f/f0 = 1.238, s = 1.632 S, $s_{20,w}$ = 1.693 S). **Top right:** Sedimentation velocity (SV) AUC data for apCC-Tet*-B at 50 krpm ($\bar{v}$ = 0.789 $cm^3$ $g^{-1}$). Continuous c(s) distribution returned a molecular mass of 16248 Da corresponding to 4.7 x monomer mass at 95% confidence level (f/f0 = 1.192, s = 1.416 S, $s_{20,w}$ = 1.478 S). **Bottom middle:** Sedimentation velocity (SV) AUC data for apCC-Tet*-$A_2B_2$ at 50 krpm ($\bar{v}$ = 0.766 $cm^3$ $g^{-1}$). Continuous c(s) distribution returned a molecular mass of 14030 Da corresponding to 4.1 x average monomer mass at 95% confidence level (f/f0 = 1.176, s = 1.467 S, $s_{20,w}$ = 1.525 S). Conditions: 150 μM peptide for SV experiments, PBS, pH 6.8, 20 °C. Residuals are shown as a bitmap below the fitted data.

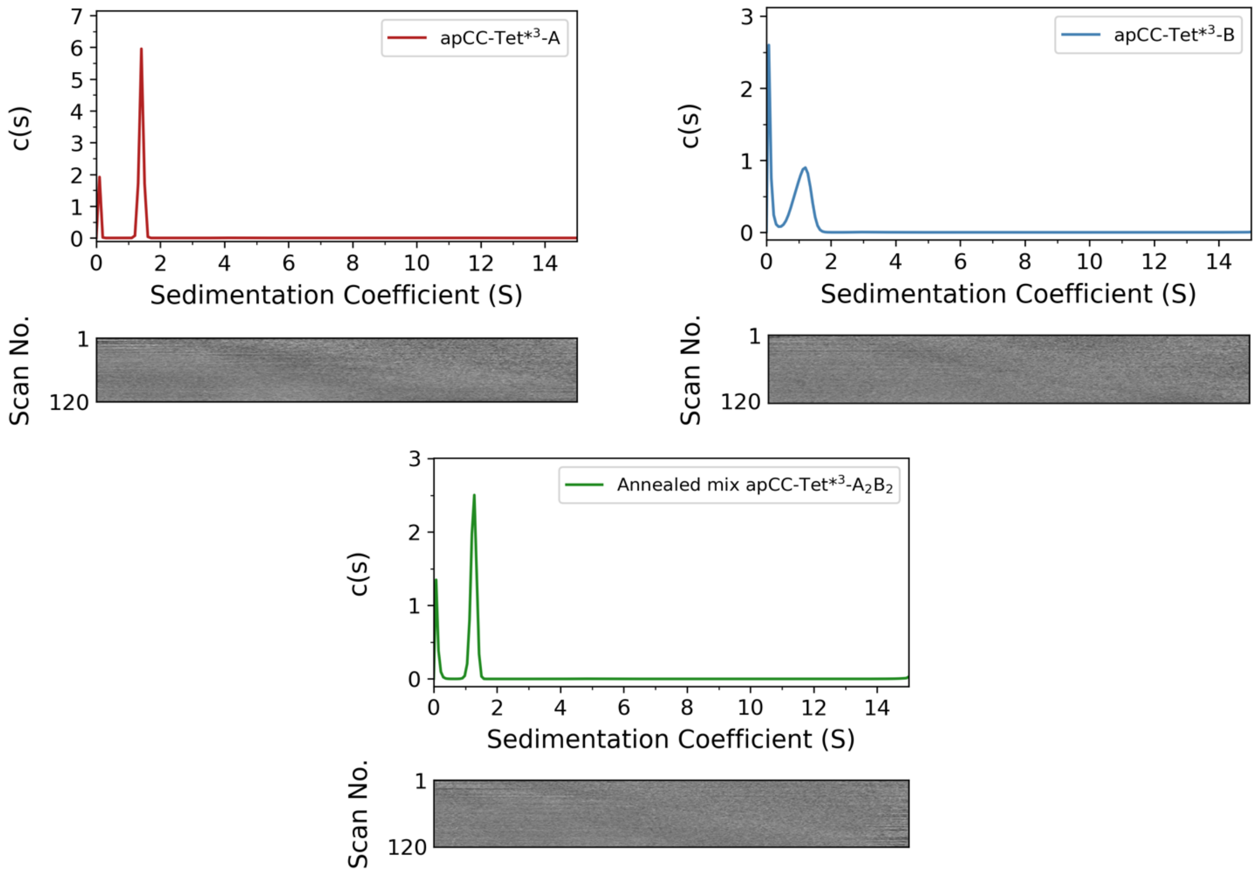

**Figure S29. AUC data isolated peptides and mixture for apCC-Tet*3-$A_2B_2$. Top left:** Sedimentation velocity (SV) AUC data for apCC-Tet*3-A at 50 krpm ($\bar{v}$ = 0.744 $cm^3$ $g^{-1}$). Continuous c(s) distribution returned a molecular mass of 11362 Da corresponding to 4.3 x monomer mass at 95% confidence level (f/f0 = 1.179, s = 1.406 S, $s_{20,w}$ = 1.459S). **Top right:** Sedimentation velocity (SV) AUC data for apCC-Tet*3-B at 50 krpm ($\bar{v}$ = 0.790 $cm^3$ $g^{-1}$). Continuous c(s) distribution returned a molecular mass of 10869 Da corresponding to 4.1 x monomer mass at 95% confidence level (f/f0 = 1.180, s = 1.092 S, $s_{20,w}$ = 1.140 S). **Bottom middle:** Sedimentation velocity (SV) AUC data for apCC-Tet*3-$A_2B_2$ at 50 krpm ($\bar{v}$ = 0.767 $cm^3$ $g^{-1}$). Continuous c(s) distribution returned a molecular mass of 10660 Da corresponding to 4.0 x average monomer mass at 95% confidence level (f/f0 = 1.135, s = 1.259 S, $s_{20,w}$ = 1.310 S). Conditions: 150 μM peptide for SV experiments, PBS, pH 7.4, 20 °C. Residuals are shown as a bitmap below the fitted data.

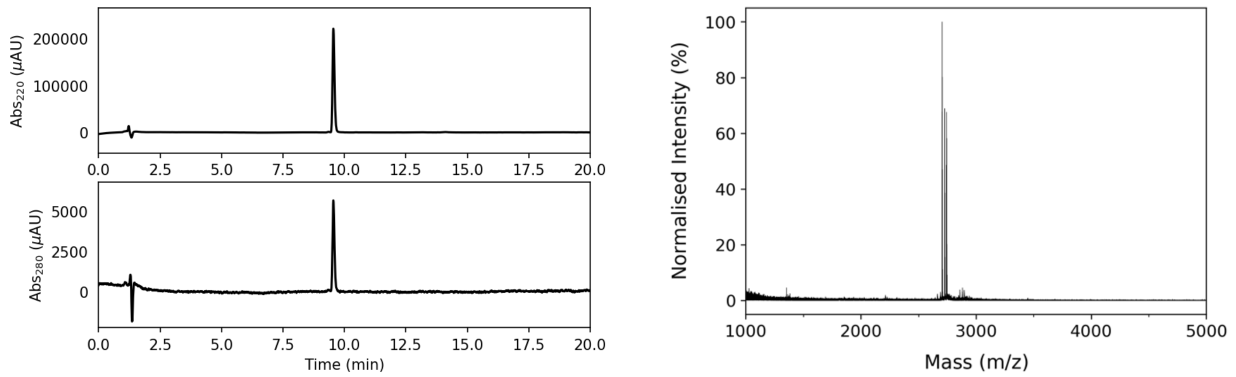

**Figure S30. Characterization of apCC-Tet$^{*3}$-B-c4CNF. Left:** Analytical HPLC chromatogram at 220 nm (top) and 280 nm (bottom) from a gradient of 20 to 80% MeCN (0.1% TFA) in $H_2O$ (0.1% TFA). **Right:** MALDI-TOF MS. Calculated mass: 2705.4 Da $[M+H]^+$. Observed mass: 2705.5 Da $[M+H]^+$.

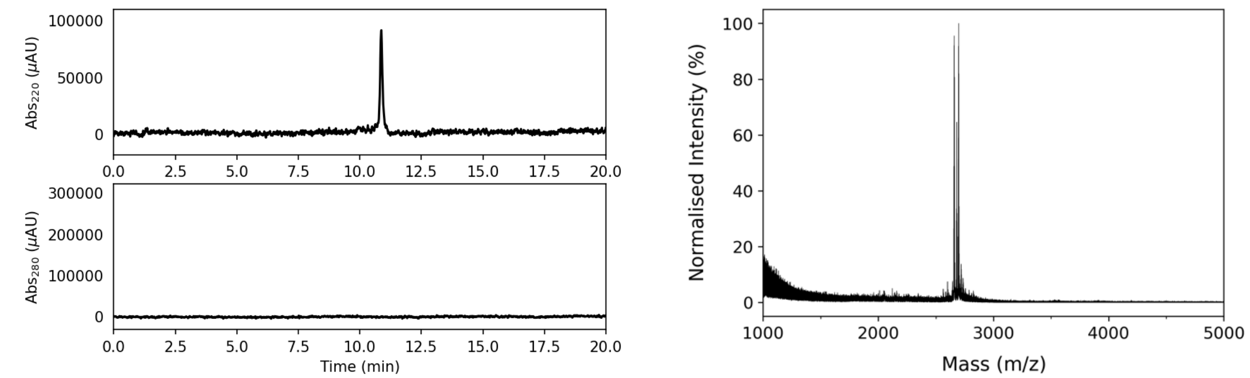

**Figure S31. Characterization of apCC-Tet$^{*3}$-A-nMSE. Left:** Analytical HPLC chromatogram at 220 nm (top) and 280 nm (bottom) from a gradient of 20 to 80% MeCN in 25 mM $NH_4HCO_3$ in $H_2O$. No chromophore was present in the sequence of apCC-Tet$^{*3}$-A-nMSE to absorb at 280 nm. **Right:** MALDI-TOF MS. Calculated mass: 2658.9 Da $[M+H]^+$. Observed mass: 2659.0 Da $[M+H]^+$.

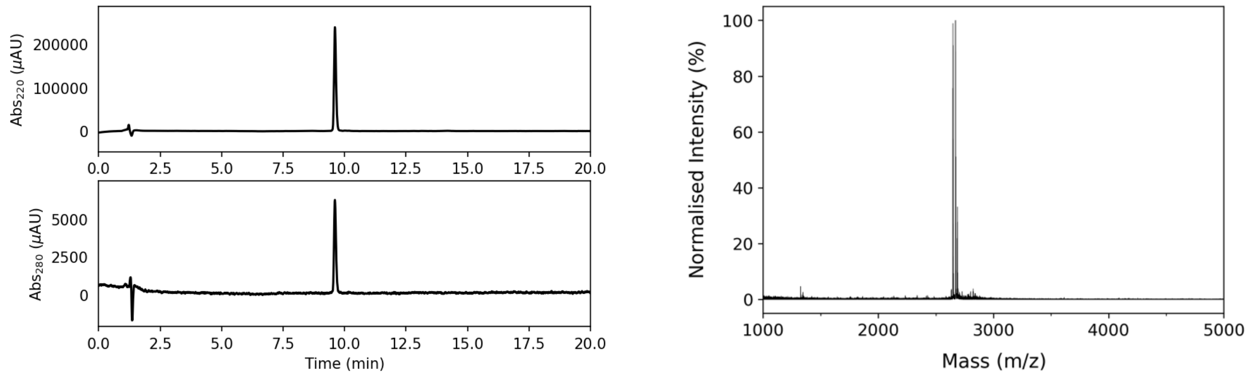

**Figure S32. Characterization of apCC-Tet$^{*3}$-B-n4CNF. Left:** Analytical HPLC chromatogram at 220 nm (top) and 280 nm (bottom) from a gradient of 20 to 80% MeCN (0.1% TFA) in $H_2O$ (0.1%

TFA). **Right:** MALDI-TOF MS. Calculated mass: 2648.3 Da [M+H]$^+$. Observed mass: 2648.6 Da [M+H]$^+$.

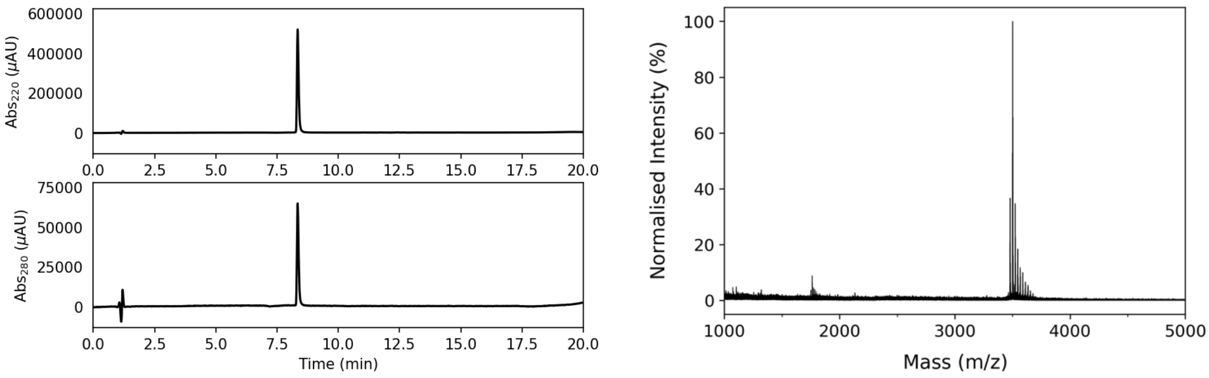

**Figure S33. Characterization of pQLL-A. Left:** Analytical HPLC chromatogram at 220 nm (top) and 280 nm (bottom) from a gradient of 20 to 80% MeCN in 25 mM $NH_4HCO_3$ in $H_2O$. **Right:** MALDI-TOF MS. Calculated mass: 3479.9 Da [M+H]$^+$, 3501.9 Da [M+Na]$^+$. Observed mass: 3479.4 Da [M+H]$^+$, 3501.4 Da [M+Na]$^+$.

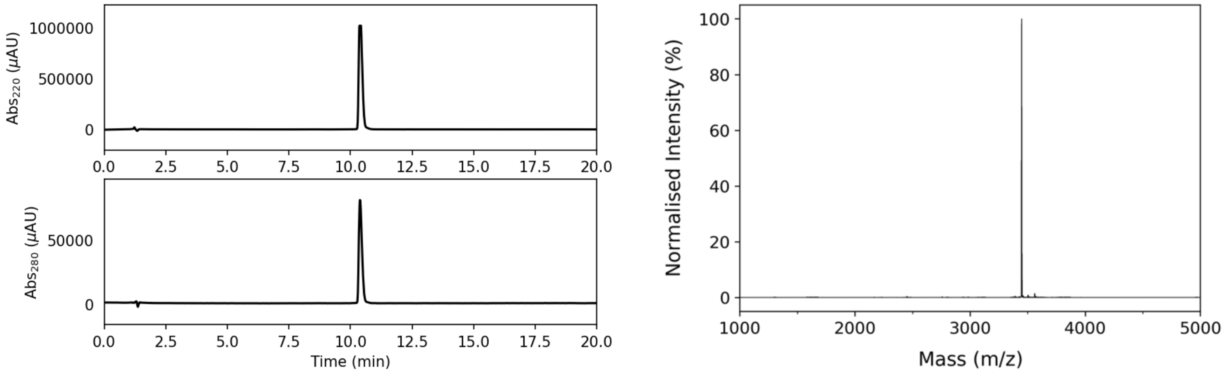

**Figure S34. Characterization of pQLL-B. Left:** Analytical HPLC chromatogram at 220 nm (top) and 280 nm (bottom) from a gradient of 20 to 80% MeCN (0.1% TFA) in $H_2O$ (0.1% TFA). **Right:** Deconvoluted ESI MS. Calculated mass: 3449.3 Da [M+H]$^+$. Observed mass: 3449.0 Da [M+H]$^+$.

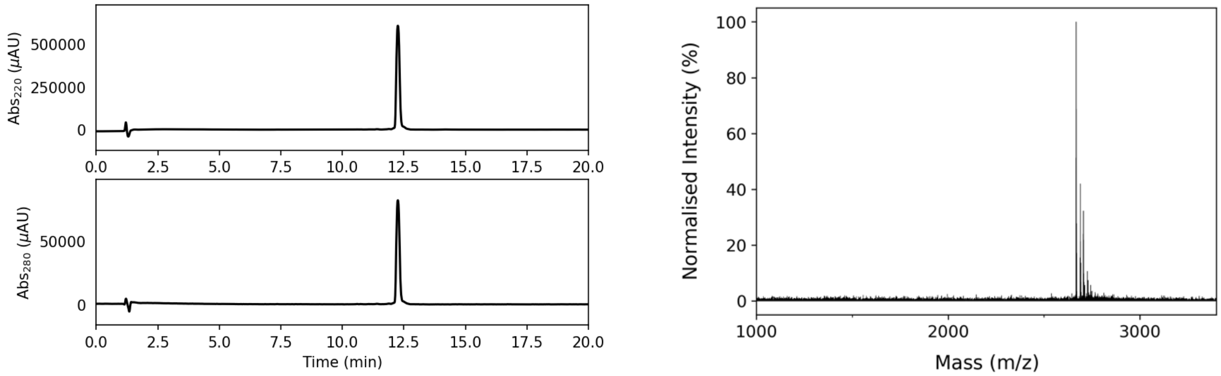

**Figure S35. Characterization of pQLL$^3$-A. Left:** Analytical HPLC chromatogram at 220 nm (top) and 280 nm (bottom) from a gradient of 20 to 80% MeCN (0.1% TFA) in $H_2O$ (0.1% TFA). **Right:** MALDI-TOF MS. Calculated mass: 2668.0 Da [M+H]$^+$. Observed mass: 2667.9 Da [M+H]$^+$.

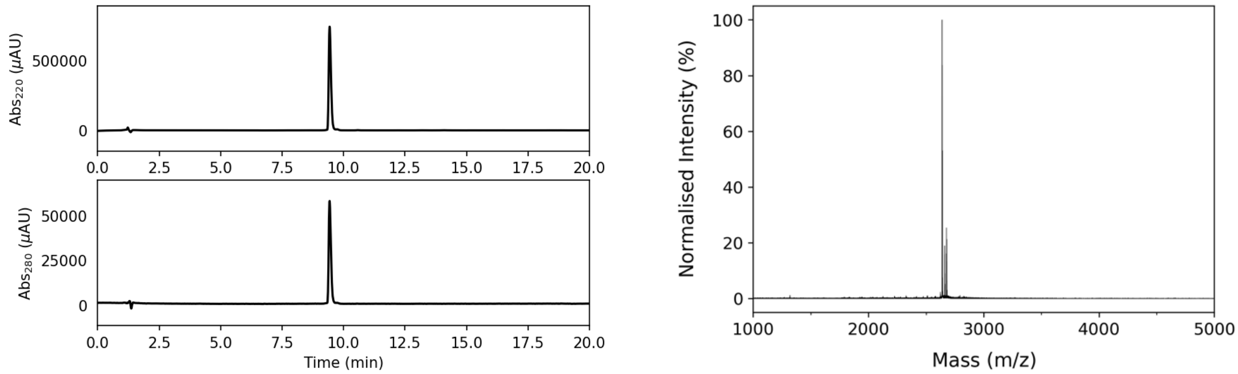

**Figure S36. Characterization of pQLL[3]-B. Left:** Analytical HPLC chromatogram at 220 nm (top) and 280 nm (bottom) from a gradient of 20 to 80% MeCN (0.1% TFA) in $H_2O$ (0.1% TFA). **Right:** MALDI-TOF MS. Calculated mass: 2639.3 Da $[M+H]^+$. Observed mass: 2639.3 Da $[M+H]^+$.

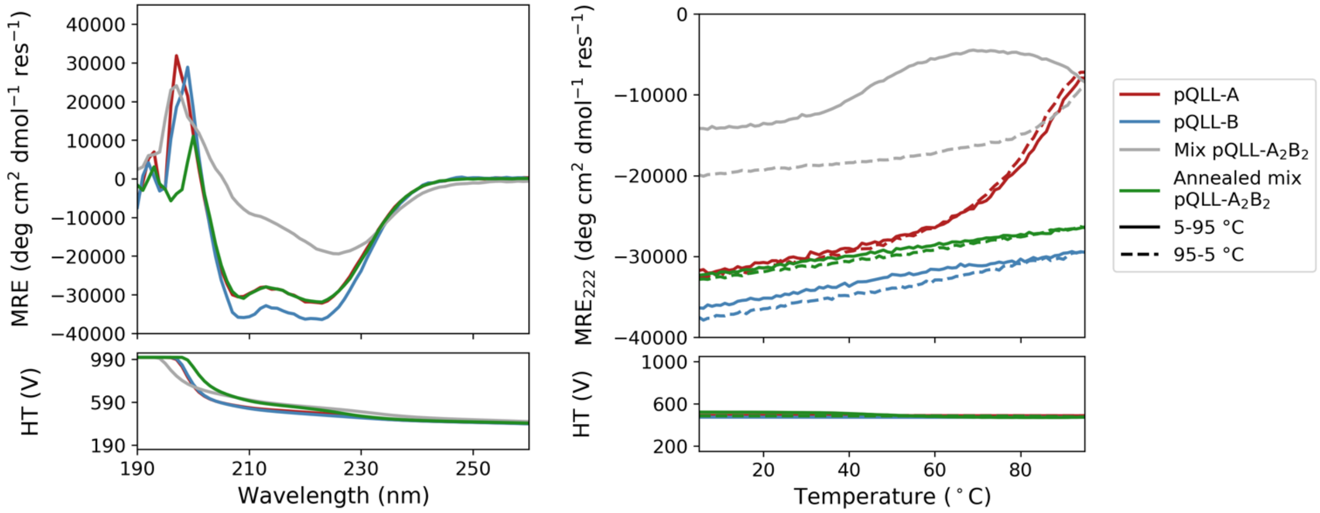

**Figure S37. CD spectroscopy of isolated peptides and mixture for pQLL-$A_2B_2$. Left:** CD spectra at 5 °C before thermal ramp up. **Right:** Thermal profiles (ramping up and ramping down) monitored at 222 nm. Bottom panels show corresponding high-tension (HT) voltages applied to the detector during measurements. Conditions: 10 μM peptide concentration, PBS (pH 7.4).

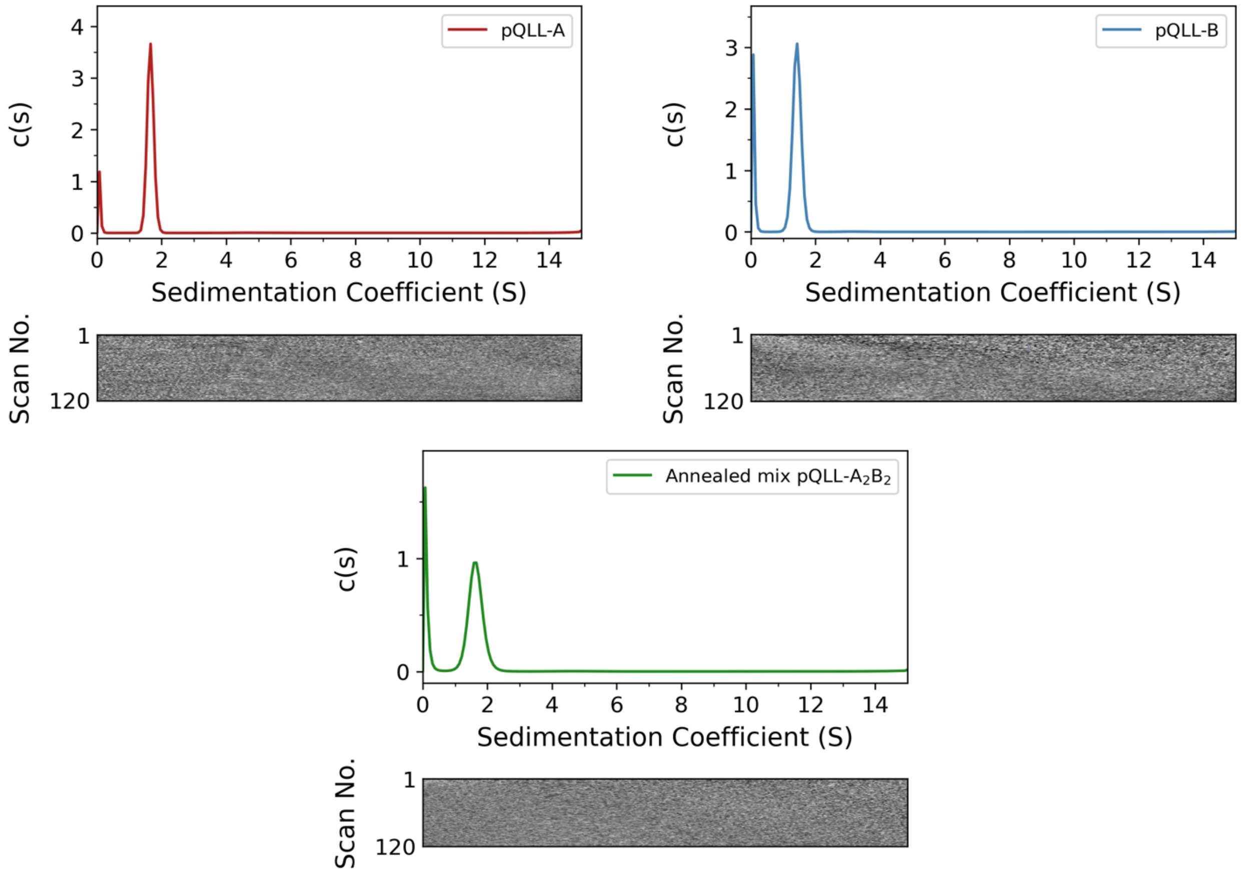

**Figure S38. AUC data isolated peptides and mixture for pQLL-$A_2B_2$. Top left:** Sedimentation velocity (SV) AUC data for pQLL-A at 50 krpm ($\bar{v}$ = 0.742 $cm^3$ $g^{-1}$). Continuous c(s) distribution returned a molecular mass of 15464 Da corresponding to 4.4 x monomer mass at 95% confidence level (f/f0 = 1.240, s = 1.654 S, $s_{20,w}$ = 1.715 S). **Top right:** Sedimentation velocity (SV) AUC data for pQLL-B at 50 krpm ($\bar{v}$ = 0.789 $cm^3$ $g^{-1}$). Continuous c(s) distribution returned a molecular mass of 18432 Da corresponding to 5.3 x monomer mass at 95% confidence level (f/f0 = 1.288, s = 1.425 S, $s_{20,w}$ = 1.488 S). **Bottom middle:** Sedimentation velocity (SV) AUC data for pQLL-$A_2B_2$ at 50 krpm ($\bar{v}$ = 0.766 $cm^3$ $g^{-1}$). Continuous c(s) distribution returned a molecular mass of 16795 Da corresponding to 4.8 x monomer mass at 95% confidence level (f/f0 = 1.188, s = 1.636 S, $s_{20,w}$ = 1.702 S). Conditions: 150 μM peptide for SV experiments, PBS, pH 7.4, 20 °C. Residuals are shown as a bitmap below the fitted data.

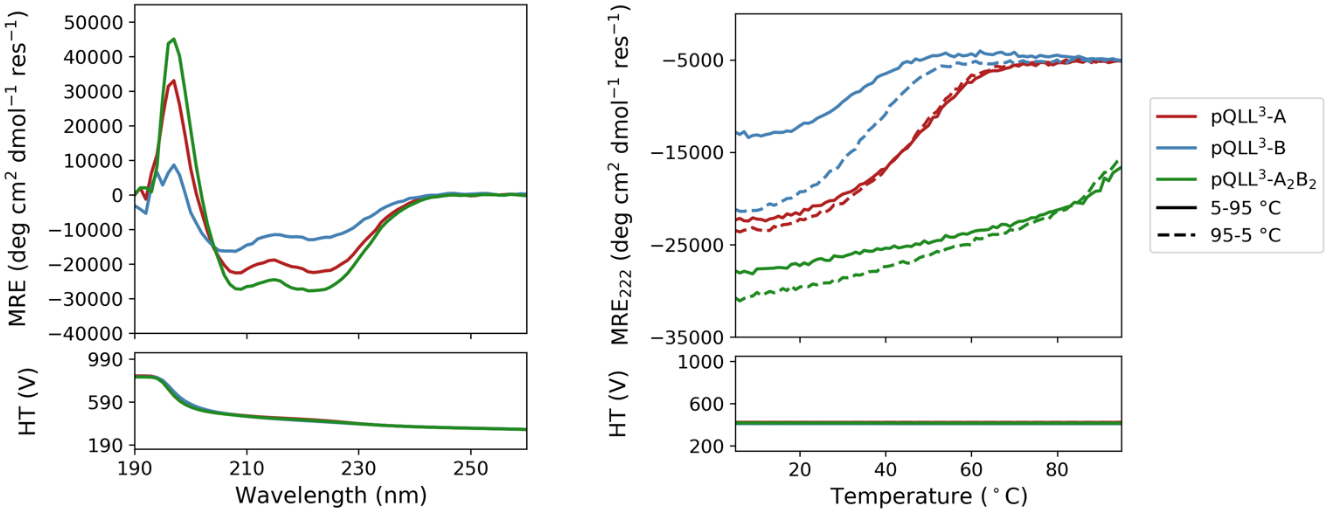

**Figure S39. CD spectroscopy of isolated peptides and mixture for pQLL$^3$-A$_2$B$_2$. Left:** CD spectra at 5 °C before thermal ramp up. **Right:** Thermal profiles (ramping up and ramping down) monitored at 222 nm. Bottom panels show corresponding high-tension (HT) voltages applied to the detector during measurements. Conditions: 10 μM peptide concentration, PBS (pH 7.4).

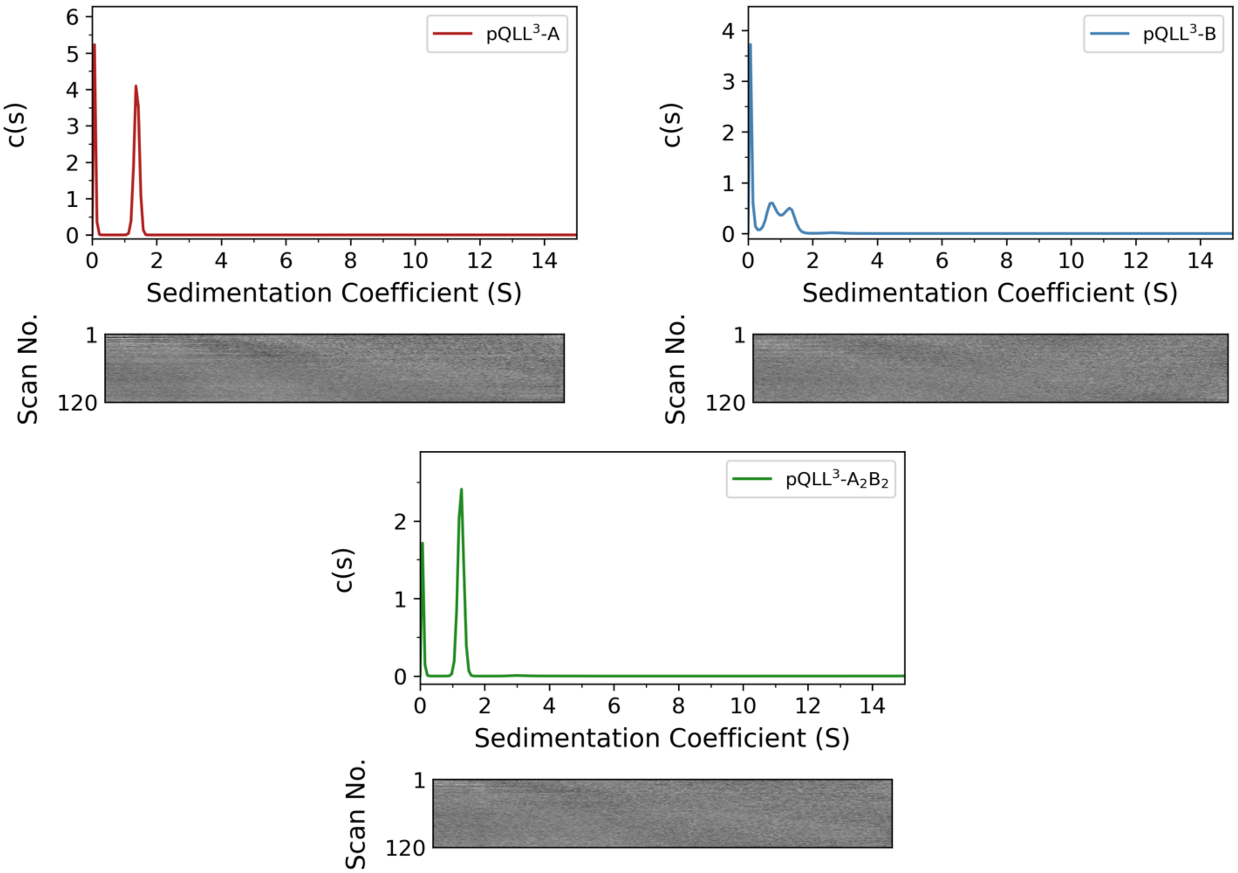

**Figure S40. AUC data isolated peptides and mixture for pQLL$^{3}$-$A_2B_2$. Top left:** Sedimentation velocity (SV) AUC data for pQLL$^{\mathbf{3}}$-A at 50 krpm ($\bar{v}$ = 0.744 cm$^3$ g$^{-1}$). Continuous c(s) distribution returned a molecular mass of 11150 Da corresponding to 4.2 x monomer mass at 95% confidence level (f/f0 = 1.187, s = 1.379 S, $s_{20,w}$ = 1.431 S). **Top right:** Sedimentation velocity (SV) AUC data for pQLL$^{\mathbf{3}}$-B at 50 krpm ($\bar{v}$ = 0.790 cm$^3$ g$^{-1}$). Continuous c(s) distribution returned a molecular mass of 7970 Da corresponding to 3.0 x monomer mass at 95% confidence level (f/f0 = 1.341, s = 0.781 S, $s_{20,w}$ = 0.815 S). **Bottom middle:** Sedimentation velocity (SV) AUC data for pQLL$^{\mathbf{3}}$-$A_2B_2$ at 50 krpm ($\bar{v}$ = 0.767 cm$^3$ g$^{-1}$). Continuous c(s) distribution returned a molecular mass of 11556 Da corresponding to 4.4 x monomer mass at 95% confidence level (f/f0 = 1.197, s = 1.260S, $s_{20,w}$ = 1.311 S). Conditions: 150 $\mu$M peptide for SV experiments, PBS, pH 7.4, 20 °C. Residuals are shown as a bitmap below the fitted data.

##### 2.2.4. Characterization of the single-chain protein

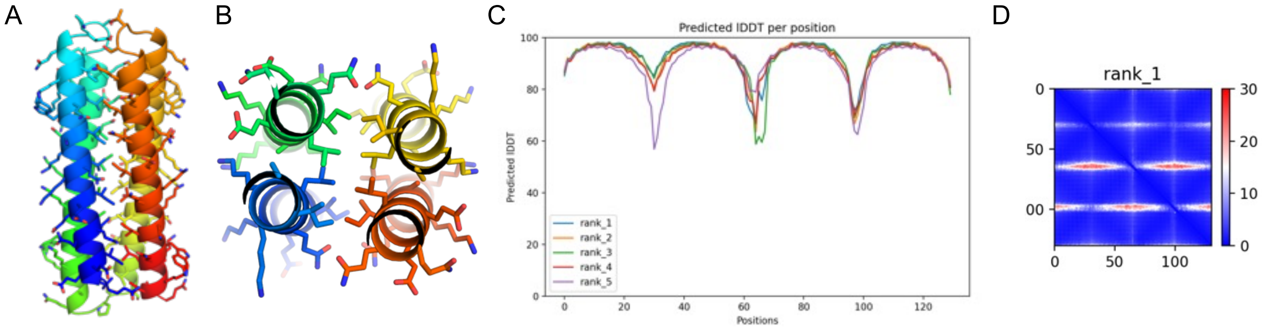

**Figure S41.** AlphaFold2 prediction of sc-apCC-4. (A&B) Orthogonal views of the top-ranked AlphaFold2 prediction for the sequence of sc-apCC-4. It is in good agreement with the narrow and wide interface organization of the parent apCC-Tet*. (C) Associated predicted lDDT and (D) Predicted Alignment Error (PAE) graphs for the sc-apCC-4 prediction.

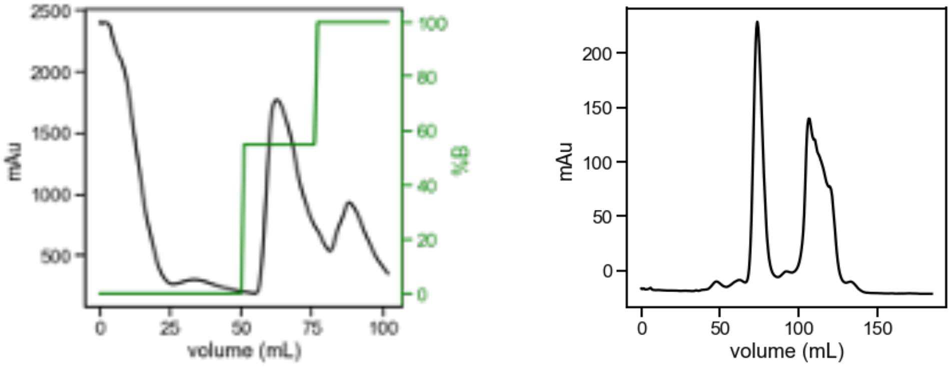

**Figure S42. FPLC of scapCC4 purification. Left:** sc-apCC-4 Ni-affinity chromatography**.** Fractions eluting at 55% B were pulled and concentrated then further purified by size exclusion chromatography. **Right:** sc-apCC-4 size exclusion chromatography. Fractions eluting at 60 mL were pooled and concentrated to be analyzed in further experiments.

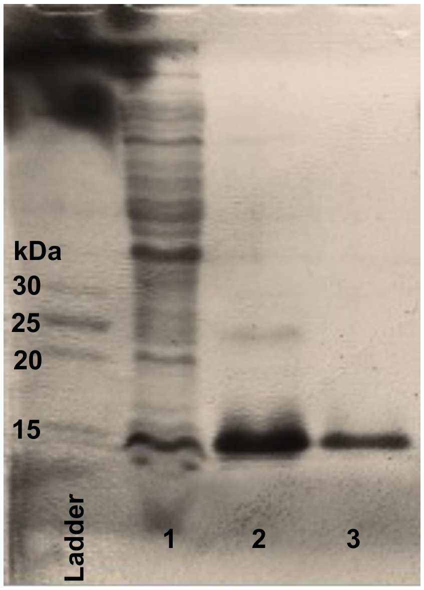

**Figure S43. SDS-PAGE of sc-apCC-4 purification.** Ladder: PageRuler Low Range Unstained Protein Ladder (Thermo Scientific), (1) sc-apCC-4 flow through from Ni-affinity chromatography, (2) combined fractions from Ni-affinity chromatography, (3) combined fractions after SEC. Expected molecular weight: 17 kDa.

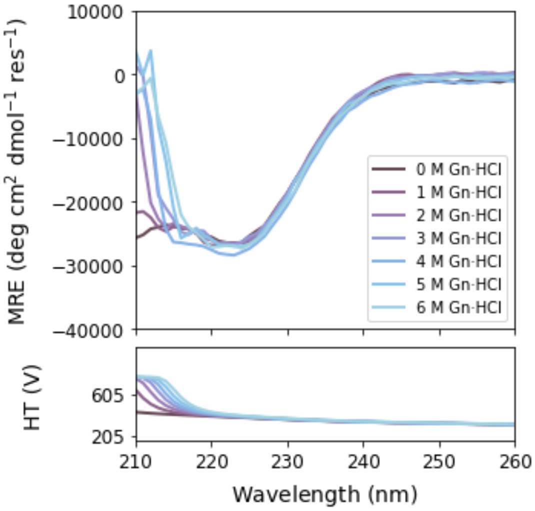

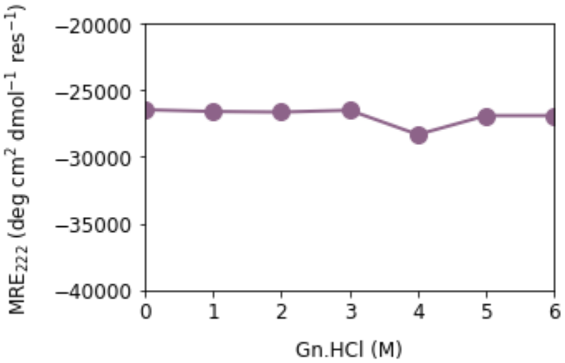

**Figure S44. Chemical denaturation of sc-apCC-4. Left:** CD full scan from 210 to 260 nm at 5 °C of sc-apCC-4 incubated with increasing concentration of guanidinium hydrochloride (Gn·HCl). **Right:** Chemical denaturation curve measured at 222 nm for sc-apCC-4 (deduced from left spectra). Conditions: 1 µM peptide concentration, ± Gn·HCl, phosphate buffer, pH 7.4 and incubation overnight before measurements.

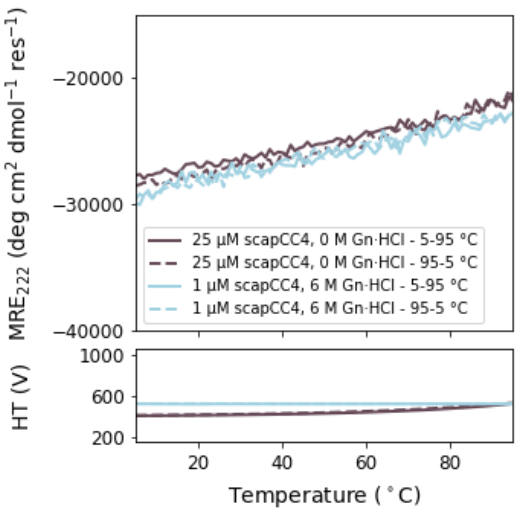

**Figure S45. Temperature profile of sc-apCC-4 in presence of high concentration of denaturant.** Thermal profiles (ramping up and ramping down) monitored at 222 nm for 1 μM of sc-apCC-4 with 6 M Gn·HCl after incubation overnight.

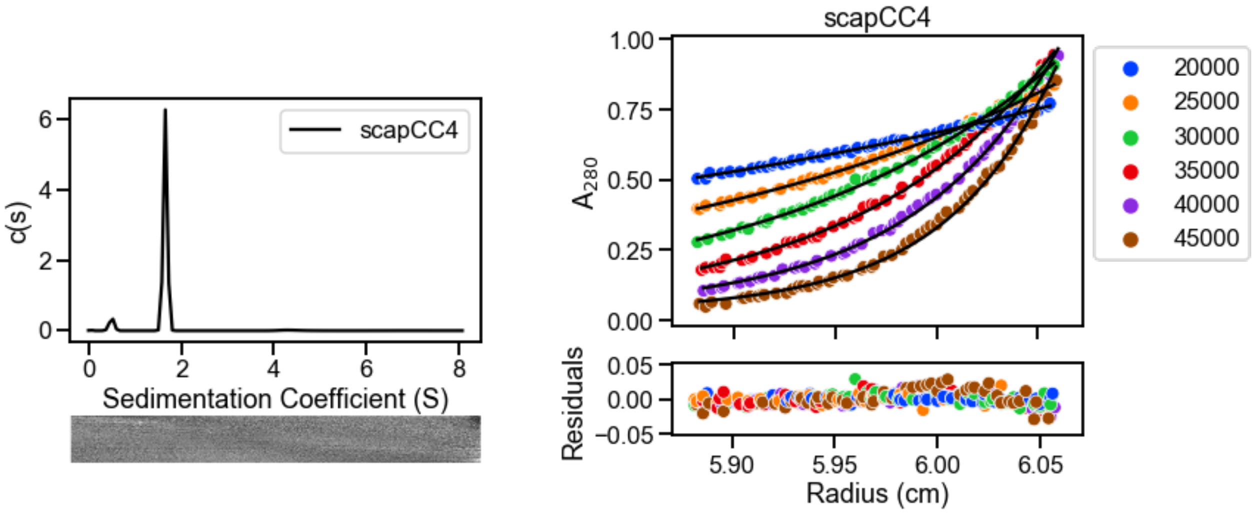

**Figure S46. AUC data for sc-apCC-4. Left:** Sedimentation velocity (SV) AUC data for sc-apCC-4 at 50 krpm ($\bar{v}$ = 0.752 $cm^3$ $g^{-1}$). Residuals are shown as a bitmap below the fitted data. Continuous c(s) distribution returned a molecular mass of 16042 Da corresponding to 0.9 x monomer mass at 95% confidence level (f/f0 = 1.21, s = 1.658 S, $s_{20,w}$ = 1.722 S). **Right:** Sedimentation equilibrium (SE) AUC data for sc-apCC-4 between 20 and 45 krpm at 5 krpm intervals. Fitted single-ideal species model curves are overlaid in black and gave a molecular mass 17009 Da corresponding to 1.0 x monomer mass, 95% confidence limits 16745-17263 Da. Conditions: 25 μM protein for SV and SE experiments, 50 mM sodium phosphate, 150 mM NaCl, pH 7.4, 20 °C

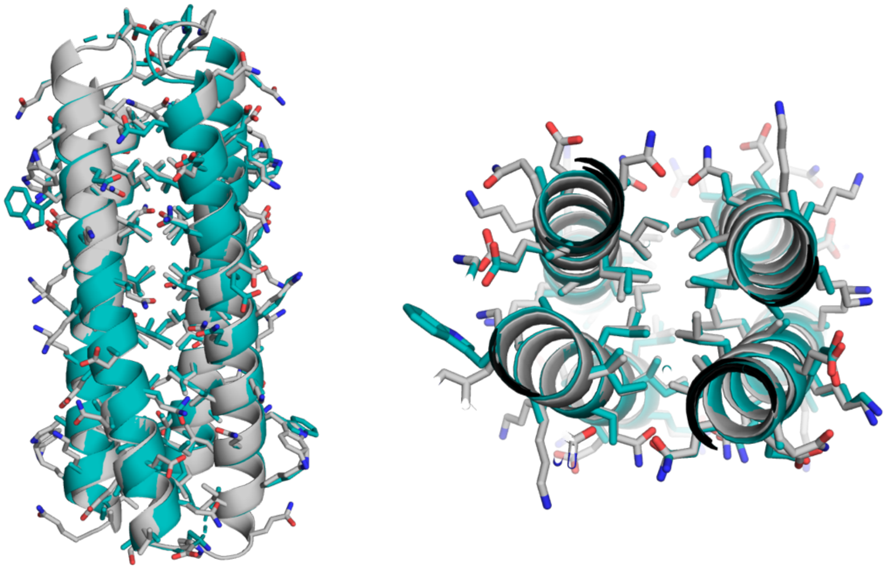

**Figure S47. Structural overlay of the crystal structure and related AlphaFold2 prediction for sc-apCC-4 protein.** Overlay of the crystal structure of sc-apCC-4 (teal) and the corresponding AlphaFold2 prediction (grey) with a $RMSD_{all\text{-}atom}$ = 0.475 Å and Cα RMSD = 0.463 Å.
